# TFAP2 paralogs facilitate chromatin access for MITF at pigmentation genes but inhibit expression of cell-cell adhesion genes independently of MITF

**DOI:** 10.1101/2021.11.23.469757

**Authors:** Colin Kenny, Ramile Dilshat, Hannah Seberg, Eric Van Otterloo, Gregory Bonde, Annika Helverson, Christopher M. Franke, Eiríkur Steingrímsson, Robert A. Cornell

## Abstract

In developing melanocytes and in melanoma cells, multiple paralogs of the Activating-enhancer-binding Protein 2 family of transcription factors (TFAP2) contribute to expression of genes encoding pigmentation regulators, but their interaction with Microphthalmia transcription factor (MITF), a master regulator of these cells, is unclear. Supporting the model that Tfap2 facilitates MITF’s ability to activate expression of pigmentation genes, single-cell seq analysis of zebrafish embryos revealed that pigmentation genes are only expressed in the subset of *mitfa*- expressing cells that also express Tfap2 paralogs. To test this model in SK-MEL-28 melanoma cells we deleted the two *TFAP2* paralogs with highest expression, *TFAP2A* and *TFAP2C,* creating *TFAP2* knockout (*TFAP2*-KO) cells. We then assessed gene expression, chromatin accessibility, binding of TFAP2A and of MITF, and the chromatin marks H3K27Ac and H3K27Me3 which are characteristic of active enhancers and silenced chromatin, respectively. Integrated analyses of these datasets indicate TFAP2 paralogs directly activate enhancers near genes enriched for roles in pigmentation and proliferation, and directly repress enhancers near genes enriched for roles in cell adhesion. Consistently, compared to WT cells, *TFAP2*-KO cells proliferate less and adhere to one another more. TFAP2 paralogs and MITF co-operatively activate a subset of enhancers, with the latter necessary for MITF binding and chromatin accessibility. By contrast, TFAP2 paralogs and MITF do not appear to co-operatively inhibit enhancers. These studies reveal a mechanism by which TFAP2 profoundly influences the set of genes activated by MITF, and thereby the phenotype of pigment cells and melanoma cells.

## Introduction

Gene expression in developing melanocytes and melanoma, a cancer derived from the melanocyte lineage, is regulated by transcription factors including Microphthalmia-associated transcription factor (MITF) and members of the SOXE, PAX and TFAP2 families [1–11]. MITF is required for differentiation of melanocytes during development, and its activity is regulated at both the transcriptional and post-translational levels [3]. In melanoma cells, high levels of MITF activity promote cell proliferation and pigmentation, while lower levels promote an invasive phenotype [12, 13]. Mass spectroscopy revealed that MITF interacts with components of both the PBAF chromatin remodeling complex, including BRG1 and CDH7, and the NURF remodeling complex, including RBBP4 [14, 15]. Furthermore, chromatin immunoprecipitation of BRG1 in cells depleted of *MITF* revealed that MITF recruits BRG1 to the promoters of specific genes, including *TYR*, which encodes the rate-limiting enzyme in melanin synthesis Tyrosinase [15]. Similar analysis suggested that SOX10 also recruits BRG1 to chromatin, and at some loci it does so in co-operation with MITF [15]. Conversely, there is evidence that PAX3 inhibits the activity of MITF at the *DCT* promoter [16]. Furthermore, low MITF activity is associated with an invasive phenotype, and deletion or knockdown of *MITF* results in upregulation of genes that promote migration and invasion [17]. MITF CUT&RUN peaks are found near some genes whose expression is upregulated in *MITF* mutant cells, implying that MITF directly represses their expression [17]. This set of MITF peaks is enriched for the binding site of FOXC1, a transcriptional repressor [18], suggesting MITF has co-factors in its repressive function as well as its activating one.

The activating enhancer-binding family of transcription factors, comprising five members, TFAP2A-E, regulate development of many cell types and organs including neural crest, placodes, epidermis, trophectoderm, heart, kidney, and brain [19–28]. In several contexts, including melanocyte differentiation, TFAP2 paralogs function redundantly [10, 29–31]. For instance, in zebrafish *tfap2a* loss-of-function mutant embryos the number of melanocytes is lower than normal and pigmentation is profoundly delayed relative to in wild-type embryos; this phenotype is exacerbated if *tfap2a* mutant embryos are also depleted of *tfap2e* expression with antisense morpholinos [10]. In zebrafish melanoma Tfap2a and Tfap2e also appear to act redundantly to promote proliferation and, interestingly, to suppress cell adhesion and cell migration [32]. Consistent with this, in the skin of mouse embryos with neural-crest specific knockout of *Tfap2a* and *Tfap2b*, the two paralogs with highest expression, fewer-than-normal cells express markers of melanocytes [33]. In addition, evidence exists for sub-functionalization among Tfap2 paralogs [34]. Of note, while *tfap2e* is the paralog with highest expression in zebrafish embryonic melanophores, it’s clear ortholog in mice, *Tfap2e,* is not expressed in embryonic melanocytes [35]. This suggests that during evolution there has been some shuffling of the function of individual TFAP2 paralogs among species.

TFAP2 paralogs and MITF appear to co-activate certain genes. For instance, in a human melanoma cell line, the *in vitro* enhancer activity of an element within an *IRF4* intron depended on the simultaneous binding of MITF and TFAP2 [36]. Further, in zebrafish *tfap2a* and *mitfa* double mutant embryos there is a greater-than-additive reduction in the number, and level of pigmentation, of melanocytes in comparison to single mutants [33]. Supporting parallel function of Tfap2 paralogs and Mitfa, the promoters of MITF target genes are enriched for TFAP2 consensus binding sites [15, 37], and ChIP-seq experiments in primary melanocytes indicate that TFAP2A and MITF bind overlapping regions of chromatin near genes encoding regulators of pigmentation [33]. Collectively, these observations indicate that TFAP2 paralogs co-activate a subset of MITF target genes by binding at the same enhancers. Still unclear, however, is whether TFAP2 paralogs and MITF act cooperatively or independently at enhancers they co-regulate. It is also unclear whether they co-repress enhancers.

TFAP2 paralogs may serve as pioneer factors for MITF, although not all evidence supports this possibility. *Pioneer* or *initiating* TFs can bind and penetrate nucleosome-bound DNA and recruit other TFs that lack this property called *settler TFs* [reviewed in 38, 39, 40]. Evidence that TFAP2 paralogs are pioneer factors includes, first, that TFAP2 binding site is over-represented within DNase1-protected “footprints” in mouse embryonic stem cells induced to differentiate [41]. Second, TFAP2A catalyzes assisted loading of androgen receptor (AR) in epididymis cells [42] and estrogen receptor in MCF-7 cells [43]. Third, the TFAP2 binding site is enriched for at the center of ATAC-seq peaks, implying it has a strong effect on chromatin accessibility [44]. Fourth, ATAC-seq peaks in naïve-stated human ESC showed reduced openness in TFAP2C-KO cells [45], and forcing expression of TFAP2C in human ESC is sufficient to open chromatin at loci where it binds [46]. Finally, TFAP2A, TFAP2B and TFAP2C can bind nucleosomes [47]. Together these findings support the possibility that TFAP2 displaces nucleosomes and thereby facilitates chromatin binding by MITF. However, it is not clear that MITF needs a pioneer factor to bind chromatin. In the *dynamic-assisted-loading model*, all classes of TFs have short residency on chromatin [reviewed in 38]. In this model, *initiating TFs* can recruit ATP-dependent chromatin remodelers (nBAF, SWI/SNF, INO80, ISWI, NURD) and thereby make chromatin accessible to other TFs, i.e., the *assisted* TFs [48]. Unlike the pioneer factor model, the assisted-loading-model predicts that initiation factors are interchangeable, and that the recruitment of chromatin remodeling complexes by initiation factors is dependent on the local chromatin structure and accessibility [48]. As mentioned above, MITF binds various components of the SWI/SNF complex [14, 49, 50] and the chromatin remodeler CHD7 [15] and so meets the criteria for an initiating factor. If the dynamic-assisted-loading model holds in this situation, MITF would have no need for a pioneer factor like TFAP2 to assist its binding to chromatin.

To address these questions, we used single cell RNA-sequencing (scRNA-seq) to investigate the temporal order of gene expression of *mitfa*, *tfap2* paralogs, and pigmentation regulators in cells of the melanocyte lineage which were included among GFP-expressing cells isolated from *Tg(mitfa-GFP)* zebrafish embryos at 28 hours post fertilization (hpf). Furthermore, to test how TFAP2 paralogs affect enhancer activity and MITF binding, we deleted TFAP2 paralogs from SK-MEL-28 melanoma cells and assessed: nucleosome positioning, using the assay for transposase-accessible chromatin with next generation sequencing (ATAC-seq); enhancer activity, using cleavage under targets and release using nuclease (CUT&RUN) with anti-H3K27Ac, anti-H3K4me3, and anti-H3K27me3; and binding of MITF, using CUT&RUN with anti-MITF. We similarly assessed binding of TFAP2A in cells harboring loss of function mutations in *MITF*. Our results support the notion that TFAP2 factors behave like the canonical pioneer factor FOXA1: at many chromatin elements bound by TFAP2A, loss of TFAP2 led to loss of enhancer activity, and in a large subset, it also resulted in chromatin becoming condensed. At both subsets of TFAP2-activated enhancers, MITF binding was TFAP2-dependent. In addition, we find evidence that TFAP2 paralogs can also inhibit enhancers, and at a subset of those, they exclude binding of MITF. Finally, the analyses suggest that TFAP2 directly inhibits many of the same genes that MITF inhibits, but we do not find evidence that TFAP2 and MITF co-repress the same enhancers. Together these findings illuminate the mechanisms by which TFAP2 and MITF coordinately regulate differentiation of melanocytes and the phenotype of melanoma cells.

## Results

### Expression of *tfap2* paralogs in the melanocyte lineage precedes activation of Mitfa transcriptional programs *in-vivo*

If Tfap2 paralogs cooperate with MITF to promote expression of melanocyte differentiation genes *in-vivo*, then high level expression of known Mitfa*-*target genes, including *dct, pmel, mreg (zgc:91968)* and *trpm1b*, should only occur in cells that express both *mitfa* and *tfap2* paralogs. To test this prediction, we performed scRNA-seq on GFP^+^ cells sorted from *Tg(mitfa:GFP)* transgenic zebrafish embryos at 28 hours post fertilization (hpf) using the 10x Chromium system. We sequenced 11,217 cellular transcriptomes and visualized the data in two-dimensional space using Seurat [51] and uniform and manifold approximation and projection (UMAP) [52] (**Fig. 1A**). We assigned cell-type annotations using previously identified marker genes [53–56] and information from the Zebrafish Information Network (ZFIN) [57]. The remaining 28 clusters comprised 11 cell types (**Fig. 1A**); neural crest and basal cell lineages were the most strongly represented (**Supplemental Fig. S1A-C**). Interestingly, only a minority of the clusters expressed *mitfa.* Sorted cells that express GFP but not *mitfa* presumably reflect either non-specific activity of the *mitfa* promoter used in the transgene or earlier transient expression of *mitfa* revealed by the long half-life of GFP; importantly, for the upcoming analysis, such cells did not express pigmentation genes. We focused on *sox10-*positive clusters 6-12 as these comprised all stages of the melanocyte lineage, including all *mitfa*-expressing cells (**Supplemental Fig. S1D**). Re-clustering of *sox10*-positive cells identified ten clusters (**Fig. 1B**). Two of these were neural crest (NC) cell populations which shared expression of bona fide NC markers (*foxd3* and *sox10*), but were distinguished by expression in one cluster of members of the Notch signaling pathway (*her6, her4* and *her12*); this cluster may represent migrating cardiac or cranial NC populations [58] (**Fig. 1C**, **Supplemental Fig. S2A, S2D**). In addition, two clusters expressed markers of a tripotent precursor of melanoblasts, iridoblasts, and xanthoblasts (*cdkn1ca, slc15a2*, *ino80e, id3, mycn, tfec*), called MIX cells [59–62] (**Supplemental Fig. S2B**). Interestingly, while both MIX clusters express *tfap2a*, the clusters differed by one and not the other expressing *tfap2e* and *tfap2b* - indeed these genes were ranked first and third, respectively, among those differentially expressed between the two MIX clusters (**Fig. 1G; Supplemental Fig. S3A; Supplemental Data Table 1**). Therefore, we will refer to the two MIX clusters as *tfap2*-low and *tfap2*-high (**Fig. 1B**). An additional cluster expressed high levels of melanoblast/ xanthoblast markers (*mitfa, impdh1b, gch2, id3*) suggesting it corresponds to the previously described MX cluster [59–62]. Another cluster was similar to the MX cluster but also expressed markers of differentiated melanophores (*dct, pmela, tyrp1b, mreg, tyr*) (**Fig. 1F-G; Supplemental Fig. S4A-B**). Other cluster pairs corresponding to xanthoblasts and xanthophores, and to iridoblasts and iridophores, were identified (**Fig. 1B, Supplemental Fig. S2C-D**).

**Figure 1:**
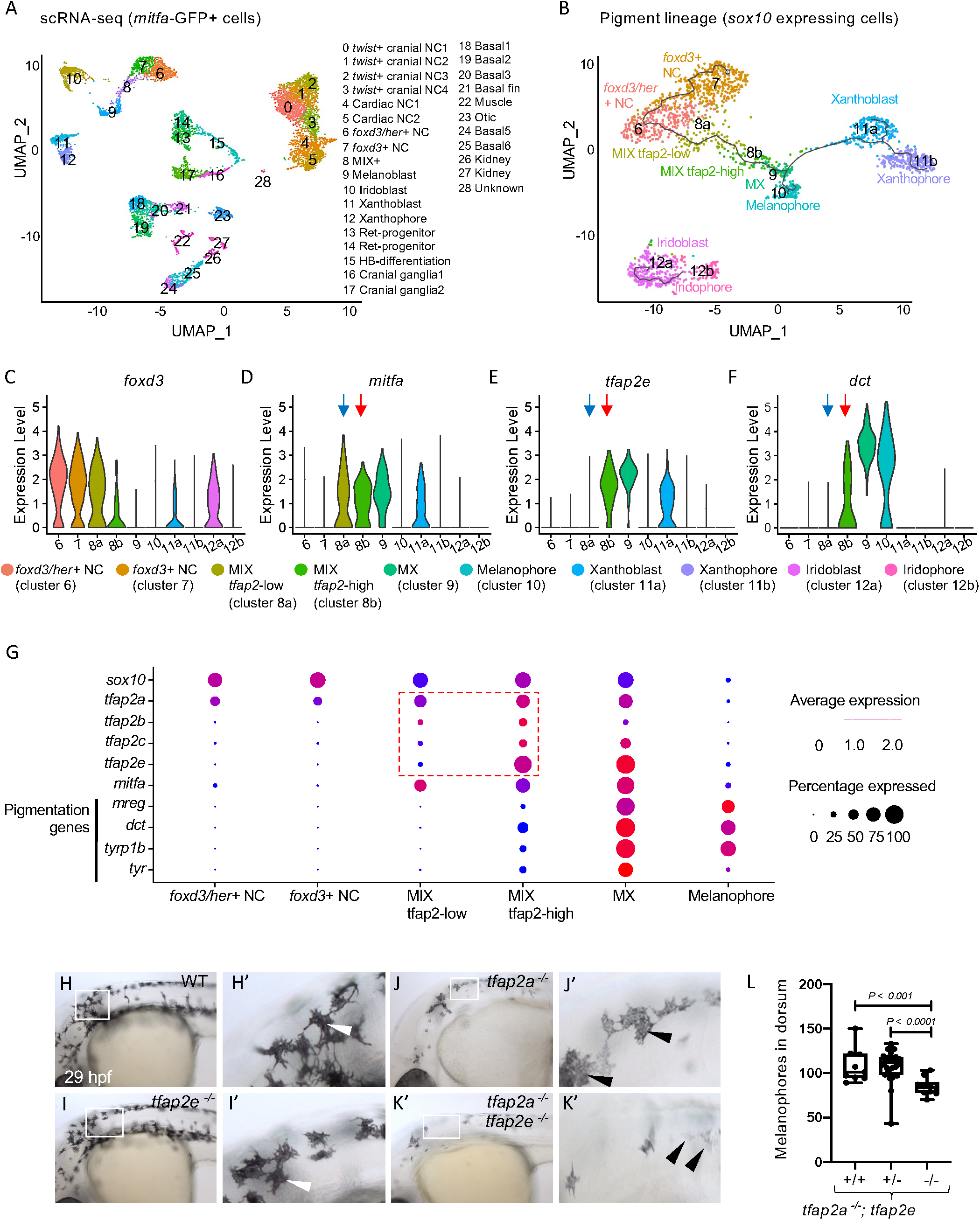
Expression of *tfap2* paralogs in the melanophore lineage precedes activation of Mitfa-target genes *in-vivo*: (**A**) Uniform Manifold Approximation and Projection (UMAP) obtained after clustering (dimensions, dims = 30, resolution = 1.2) GFP+ cells (n = 11,217 cells) sorted from *Tg(mitfa:GFP)* zebrafish embryos at 28 hours post fertilization (hpf). Annotated cell clusters as labelled. (**B**) UMAP obtained after re-clustering *sox10*-expressing clusters 6-12 (n = 1918 cells). Black line - Monocle pseudotime trajectory analysis starting at *foxd3*+ neural crest cells showing the progression through different pigment cell clusters as shown. (**C-F**) Violin plots showing expression of select genes *foxd3, mitfa, tfap2e* and *dct* for each cell cluster represented in B. *Blue arrows* point to the MIX-*tfap2*-low cluster, and show that *mitfa*, but not pigmentation genes, are expressed in this cluster (*tfap2a* is also expressed in this cluster). *Red arrows* point to the MIX-*tfap2*-high cluster, and show that *mitfa*, *tfap2e*, and pigmentation genes are expressed in this cluster. (**G**) Dot plot representing the expression of *sox10*, *tfap2* paralogs, *mitfa* and Mitfa-target genes, and the percentage of cells expressing these genes in neural crest, MIX, MX and melanophore clusters. Expression of *tfap2* paralogs are highlighted in MIX *tfap2*-low and MIX *tfap2*-high clusters (red box). Size of dots represents percentage of cells expressing the gene, and blue-red scale (low-high) represents relative average expression among cells. (**H-K**) Lateral views of head and trunk of live embryos at 29 hpf, anterior to the left and dorsal to the top. Genotype as shown. Boxes, regions magnified in accompanying panels H’-K.’ (**H-H’**) A wild-type or heterozygous mutant (sibling) embryo with normal melanophores (white arrowheads). (**I-I’**) A *tfap2e^ui157ui/157^* embryo, with melanophores that are normal in terms of number, differentiation and pigmentation (white arrowheads). (**J-J’**) A *tfap2a^low/low^* homozygous mutant embryo, with fewer melanophores than *tfap2e^ui157ui/157^* and WT sibling embryos. (**K-K’**) A *tfap2a^low/low^; tfap2e ^ui157ui/157^* double-mutant embryo, with fewer and paler melanophores than in *tfap2a^low/low^* siblings. (**L**) Box plot illustrating the number of pigmented melanophores in the dorsum of *tfap2a^low/low^*, *tfap2a^low/low^; tfap2e^+/ ui157^*, and *tfap2a^low/low^*; *tfap2e^ui157ui/157^* double mutant embryos at 36 hpf. Center line, mean; box limits, upper and lower quartiles; whiskers, minimum and maximum values; black dots, number of melanocytes per individual embryo (*tfap2a^low/low^*; n=9, *tfap2a^low/low^; tfap2e^+/ui157^;* n=32, *tfap2a^low/low^; tfap2e^ui157/ui157^*, n=10). P-value according to the Student’s t-test.

Pseudotime analysis supports a lineage trajectory for the melanophore lineage leading from NC, to *tfap2*-low MIX cells, to *tfap2*-high MIX cells, to MX, to melanophores (**Fig. 1B**). Within this trajectory, despite expression of *mitfa* in both *tfap2*-low and *tfap2*-high MIX clusters, expression of select pigmentation genes (*dct, pmel, mreg* and *trpm1b*) is first detected in *tfap2*-high MIX cells, and becomes higher in MX cells concomitant with higher level expression of *mitfa* and of *tfap2e* (**Fig. 1C-G; Supplemental Fig. S4A-B**). These data support a cooperative role for Mitfa and Tfap2 paralogs in activation of pigmentation genes.

### Tfap2a and Tfap2e redundantly promote the differentiation of zebrafish embryonic melanocytes

The scRNA-seq analysis revealed expression of four *tfap2* paralogs at some point within the embryonic melanophore lineage, i.e., in NC cells (*tfap2a*), in *tfap2*-low MIX cells (*tfap2a*), *tfap2*-high MIX cells (*tfap2a, tfap2b, tfap2c*, and *tfap2e*), and in MX (*tfap2a, tfap2c* and *tfap2e*) (**Fig. 1G; Supplemental Fig. S3A**); expression of all *tfap2* paralogs is reduced upon terminal differentiation of melanophores (cluster 10). An overt phenotype of reduced melanocyte numbers and delayed melanization is evident in *tfap2a* mutants [23, 63, 64] and morphants [65], but not in *tfap2b* morphants [56], *tfap2c* mutants [29, 66], or *tfap2e* morphants [10]. However, depletion of *tfap2e* from *tfap2a* mutants yields a more severe phenotype of reduced melanocyte numbers and delayed melanization in comparison to control-MO-injected *tfap2a* mutants [10]. Here we used zinc finger nucleases to generate a 157bp deletion of exon 2 in *tfap2e*. Similar to the findings with morpholinos, homozygous mutants were phenotypically normal (**Supplemental Fig. S5A-C**) while *tfap2a/tfap2e* double mutants had fewer embryonic melanophores than *tfap2a* single mutants, and pigmentation was further delayed (**Fig. 1H-L**). Nonetheless, by 48 hpf the intensity of pigmentation within individual melanophores was not overtly different between wild-type and *tfap2a/tfap2e* double mutants (**Supplemental Fig. S5G-J**).

Importantly, the phenotypes in single or double mutants thus far examined may be suppressed by compensatory upregulation of other Tfap2 paralogs. Indeed, we detected upregulated expression of *tfap2c* in *tfap2a* mutants and in *tfap2a/tfap2e* double mutants, presumably driven by non-sense mediated decay (**Supplemental Fig. S6**). In conclusion, the *in-vivo* consequence of loss of Tfap2 function upon melanophore differentiation is difficult to assess because of the potential for redundant function among Tfap2 paralogs and because of pleiotropy (see Discussion), nonetheless mutant embryos with lower-than-normal expression of *tfap2* paralogs exhibit fewer melanophores than wild-types, and these melanophores are delayed in pigmentation.

### TFAP2A binds open and closed chromatin

To test how TFAP2 paralogs interact with MITF we switched to a cell line model. We chose the SK-MEL-28 melanoma cell line because we have deleted all alleles of *MITF* from this cell line and have evaluated MITF binding genome-wide using CUT&RUN [17]. Here we carried out (1) CUT&RUN using antibodies to TFAP2A (i.e., TFAP2A peaks), (2) CUT&RUN using antibodies to chromatin marks indicative of active regulatory elements (H3K27Ac and H3K4Me3) [67, 68] and indicative of inactive chromatin (H3K27Me3) [69], and (3) ATAC-seq to distinguish between open and closed chromatin [70]. We used IgG as a background control and the MACS2 software to call peaks in each CUT&RUN dataset (**Supplemental Fig. S7A-B**). Based on proximity to transcriptional start sites (TSS), about one-third of TFAP2A peaks appeared proximal to promoters (within 3kb of a TSS). As expected, these elements had strong H3K4Me3 signal (**Supplemental Fig. S7B**). At promoter-proximal TFAP2A peaks, the H3K27Ac signal in WT cells was relatively consistent, whereas at promoter-distal TFAP2A peaks (greater than 3 kb but within 100 kb of a TSS) the H3K27Ac signal ranged from high to background level (**Supplemental Fig. S7B-C**). About two-thirds of TFAP2A peaks overlapped ATAC-seq peaks, indicating that they were in open chromatin (**Supplemental Fig. S7D-E**). Of note, the read depth (height) of a peak approximates the number of chromosome molecules where TFAP2A binds. The average read depth of the TFAP2A peaks in closed chromatin was only about 50% of that in open chromatin but was nonetheless 80-fold higher than the IgG background read depth (**Supplemental Fig. S7D-E, Supplemental Fig. S8A-B for example loci**). Importantly, the TFAP2 binding site was strongly enriched for in both TFAP2A-bound elements where the local ATAC-seq signal was called as a peak and in counterparts where it was not (*p* < 1 x 10^-1785^ and *p* < 1 x 10^-4375^, respectively), supporting the idea that TFAP2A binds DNA directly even when the DNA is occupied by nucleosomes (**Supplemental Fig. S8C-D**). These results indicate that TFAP2A binds at both open and closed chromatin, consistent with it being a pioneer factor, and at enhancers and promoters with a range of activity levels.

### At a subset of TFAP2-activated enhancers, TFAP2 is necessary for chromatin accessibility

Having shown that TFAP2A can bind closed chromatin, we next asked whether TFAP2 factors play a role in opening chromatin [39]. We used CRISPR/Cas9 methods to introduce frame-shift mutations into *TFAP2A* and *TFAP2C*, the *TFAP2* genes with high expression in SK-MEL-28 cells (**Supplemental Fig. S9A-D**). We then carried out RNA-seq, ATAC-seq, and CUT&RUN with antibodies to H3K27Ac, H3K4me3, and H3K27me3, in two independent knockout clones (hereafter, *TFAP2*- KO cells). Western blot analysis showed an absence of immunoreactivity for both proteins (**Supplemental Fig. 9E**). Control clones (hereafter WT cells) were derived from the parental SK-MEL-28 line transiently transfected with Cas9 but not with guide RNAs. RNA-seq revealed that expression of 532 genes was downregulated, and expression of 609 genes was upregulated, in *TFAP2*-KO cells (i.e., in both clones) versus in WT cells. We will refer to these sets as TFAP2-activated genes and TFAP2-inhibited genes, respectively.

We defined TFAP2-activated enhancers as TFAP2A peaks a) that overlap H3K27Ac and ATAC-seq peaks in WT cells (21,745/ 36,948, or 59% of all TFAP2A peaks), b) that are greater than 3 kb from a transcription start site (11,005/ 21,745, or 51% of TFAP2A peaks at H3K27Ac/ATAC peaks), and c) where the H3K27Ac signal was significantly lower in *TFAP2*-KO cells relative to in WT cells (adj p < 0.05, log2FC <- 1; 3,858/11,005 or 35% of TFAP2A-bound enhancers). Interestingly, at about half the TFAP2-activated enhancers (2002/3858), the ATAC-seq signal was significantly lower in *TFAP2*-KO versus in WT cells (adj. p < 0.05, log2FC <-1) (**Fig. 3A, E-E’**). The open status of these loci depends on TFAP2 paralogs, and we therefore infer that TFAP2 paralogs directly or indirectly recruit the nucleosome-displacing machinery. We further infer that TFAP2 paralogs bound these loci as closed chromatin, but it is also possible they bound to another pioneer factor present at these loci. In either case, it participates in a pioneering function, and we refer to this subset of enhancers as TFAP2-pioneered-and-activated enhancers (**Fig. 2 Box 1A**). We refer to the subset where ATAC-seq signal is unchanged between *TFAP2*-KO and WT cells as non-pioneered TFAP2-activated enhancers **(Fig. 2 Box 1B; Fig. 3B, F-F’).**

**Figure 2:**
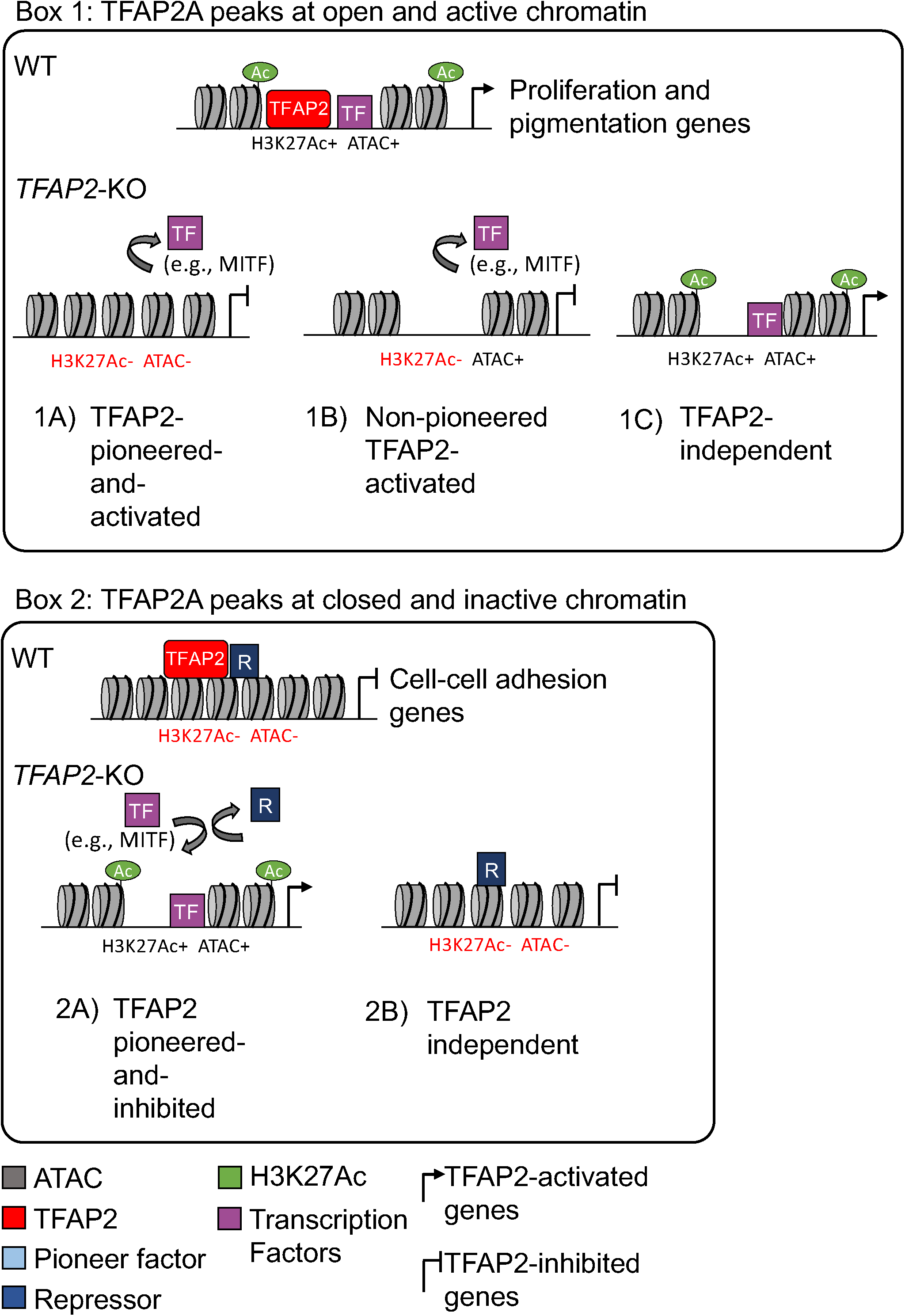
Categories of TFAP2-regulated enhancers. Box 1: TFAP2A peaks at open and active chromatin. (**1A**) TFAP2-pioneered-and- activated enhancers show reduced nucleosome accessibility (ATAC-seq) and reduced levels of active chromatin marks (H3K27Ac and H3K4Me3) in *TFAP2-KO* cells compared to in WT cells. We infer that TFAP2 paralogs pioneer chromatin access for transcriptional co-activators, like MITF and SOX10 (purple box), that in turn recruit chromatin remodelling enzymes and histone modifying enzymes. (**1B**) Non-pioneered TFAP2-activated enhancers show loss of active chromatin marks but unchanged nucleosome accessibility in *TFAP2*-KO cells compared to in WT cells. At these enhancers, we infer that TFAP2 paralogs recruit the binding of transcription factors that, in turn, recruit histone modifying enzymes. TFAP2 paralogs also may recruit such enzymes. It is possible that these elements are stably pioneered by TFAP2 paralogs [94]. (**1C**) At TFAP2-independent elements, neither the nucleosome accessibility nor active histone marks are altered in *TFAP2*-KO cells relative to in WT cells. Both types of TFAP2-activated enhancer are significantly enriched near genes whose expression is reduced in *TFAP2*-KO cells relative to in WT cells (i.e., TFAP2- activated genes). Such genes are associated with the gene ontology (GO) terms cell proliferation and pigmentation. TFAP2-independent elements are associated with neither TFAP2-activated nor TFAP2-inhibited genes. **Box 2**: TFAP2A peaks at closed and inactive chromatin. (**2A**) TFAP2-pioneered-and-inhibited enhancers show increased nucleosome accessibility and increased levels active chromatin marks in *TFAP2*-KO cells compared to in WT cells. At these sites we infer that TFAP2 paralogs recruit or stabilize the binding of enzymes that condense chromatin and that inhibit the binding of transcriptional activators that are otherwise inclined to bind at them. These elements are significantly enriched near genes whose expression is elevated in *TFAP2*-KO cells relative to in WT cells (i.e., TFAP2-inhibited genes). Such genes were associated with the GO terms cell-cell adhesion and cell migration. (**2B**) At TFAP2-independent elements, neither the nucleosome accessibility nor active histone marks are altered in *TFAP2*-KO cells relative to in WT cells. These elements were associated with neither TFAP2-activated nor TFAP2-inhibited genes.

**Figure 3:**
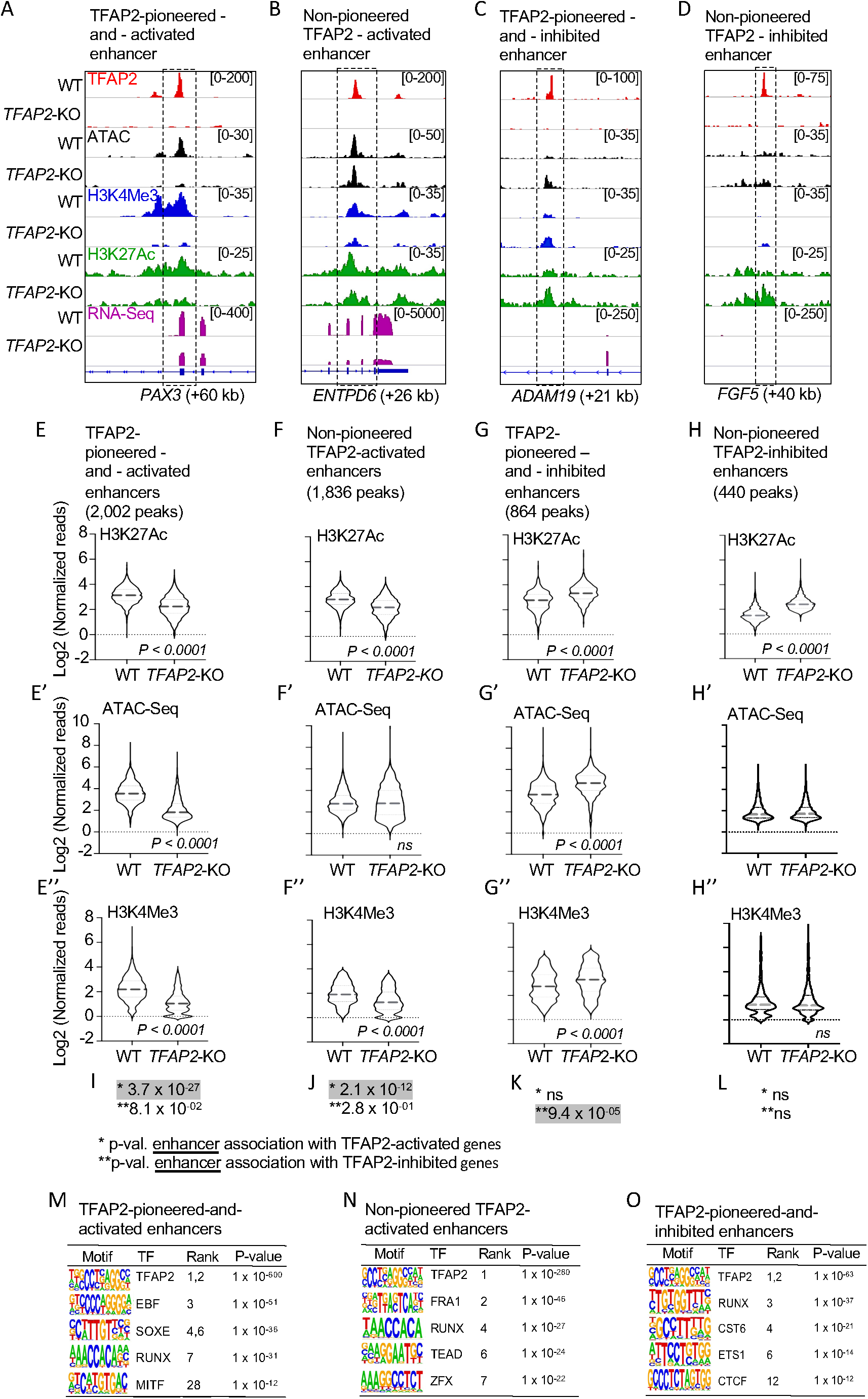
TFAP2 paralogs facilitate gene expression by opening and condensing chromatin. (**A-D**) Screenshot of IGV genome browser (GRCH37/hg19), visualizing anti-TFAP2A CUT&RUN-seq (red), ATAC-seq (black), anti-H3K4Me3 CUT&RUN-seq (blue), anti-H3K27Ac CUT&RUN-seq (green) and RNA-seq (magenta) datasets at: (**A**) an TFAP2-pioneered-and-activated enhancer at the *PAX3* (+60 kb) locus; (**B**) a non-pioneered TFAP2A-activated enhancer at the *ENTPD6* (+26kb) locus; (**C**) an TFAP2-pioneered-and-inhibited enhancer at the *ADAM19* (+21 kb) locus; and (**D**) a non-pioneered TFAP2A-inhibited enhancer at the *FGF5* (+40kb) locus. Genotypes as labeled; y-axis in E applies to E-H, etc. (**E-H’’**) Violin plots conveying normalized reads of (**E-H**) anti-H3K27Ac (two independent replicates), (**E’-H’**) ATAC-seq (four independent replicates) and **(E’’-H”)** anti- H3K4Me3 (two independent replicates) CUT&RUN-seq at **E-E”**) TFAP2-pioneered- and-activated enhancers, (**F-F’’**) non-pioneered TFAP2-activated enhancers, (**G-G’’**) TFAP2-pioneered-and-inhibited enhancers, and (**H-H’’**) non-pioneered TFAP2- inhibited enhancers. The number of peaks in each group is indicated. P-values shown were determined by Student’s t-test. (**I-L**) Hypergeometric analysis of TFAP2 regulated enhancers at TFAP2-activated (*) and TFAP2-inhibited (**) genes in WT cells (FDR < 0.05, |log2FC| > 1). ns; not significant. (**M-O**) Enrichment of transcription factor binding motifs at (**M**) TFAP2-pioneered-and-activated enhancers, at (**N**) non-pioneered TFAP2-activated enhancers and at (**O**) TFAP2-pioneered-and- inhibited enhancers as determined using HOMER motif analysis. P values were calculated using ZOOPS scoring (zero or one occurrence per sequence) coupled with hypergeometric enrichment analysis. TF; transcription factor.

Both TFAP2-pioneered-and-activated enhancers and non-pioneered TFAP2A- activated enhancers were associated with TFAP2-activated genes. Interestingly, the association was stronger for the former subset (**Fig. 3 I, J**; **Table 1,** compare rows 4 and 5). Moreover, at both subsets, the H3K4me3 signal, which is associated with enhancer activity [68], was reduced in *TFAP2*-KO cells relative to WT cells (**Fig. 3E’’, F’’**). While both subsets were strongly enriched for the TFAP2 binding site and certain other binding sites (e.g., RUNX), the subset pioneered by TFAP2 was more strongly enriched for the SOXE and MITF binding sites, while the non-pioneered subset was more strongly enriched for the FRA1, TEAD and the ZFX binding sites (**Fig. 3M, N**). Of note, FRA1 is a pioneer factor [71] which could explain why these elements do not depend on TFAP2 to be free of nucleosomes.

**Table 1:** TFAP2-regulated enhancers and their association with TFAP2- regulated genes (Fisher’s Exact Test, hypergeometric analysis):

| Effect of TFAP2 on: |  |  |  | Association with TFAP2-activated genes | Association with TFAP2-inhibited genes |
| --- | --- | --- | --- | --- | --- |
| Enhancer | H3K27Ac | Chromatin | N (# of elements) | p-value \|Log2FC\| >1 | p-value \|Log2FC\| >1 |
| All TFAP2-activated | Increases | Any effect or none | 3,838 | $1.12 \times 10^{-24}$ | ns |
| TFAP2-pioneered-and-activated | Increases | Opens | 2,002 | $3.7 \times 10^{-27}$ | ns |
| Non-pioneered TFAP2-activated | Increases | None | 1,836 | $2.1 \times 10^{-12}$ | ns |
| All TFAP2-inhibited | Decreases | Any effect or none | 1,304 | ns | $4.8 \times 10^{-06}$ |
| TFAP2-pioneered-and-inhibited | Decreases | Condenses | 864 | ns | $9.44 \times 10^{-05}$ |
| Non-pioneered TFAP2-Inhibited | Decreases | None | 440 | ns | ns |
| All TFAP2A peaks | Any effect or none | Any effect or none | 36,948 | $2.4 \times 10^{-09}$ | ns |
| TFAP2A peaks | None | None | 23,735 | ns | ns |

Column 1: groups of enhancers directly bound by TFAP2A, determined by CUT&RUN. Pioneered, refers to the set of enhancers where the chromatin is opened or condensed by TFAP2. Non-pioneered, refers to the set of enhancers where only the H3K27Ac signal is activated or repressed by TFAP2 (i.e. where the ATAC-Seq signal is independent of TFAP2 binding). Columns 2-3: TFAP2-dependent H3K27Ac signal (increased or decreased) and/or TFAP2-dependent nucleosome depleted regions (ATAC-Seq; opened or condensed) at TFAP2 bound enhancers. Column 4: the number of target elements regulated by TFAP2. Columns 5-6: the effect of TFAP2 on gene expression (two independent clones, 4 replicates each) and WT (4 replicates). Differential gene expression (adjusted p-value < 0.05) in *TFAP2*-KO cells and WT cells was determined by RNA-Seq. The p-value from the Fisher's Exact Test in which the null hypothesis is that no association exists. DEG: differentially expressed genes. Attention should be focused on columns 5-6 which indicate the strength of association between the regulation status of a gene and the TFAP2-regulated enhancer of interest; significant associations are highlighted yellow. ns; not statically significant.

### TFAP2A inhibits enhancers by blocking the opening of chromatin

Because of evidence that TFAP2A directly represses gene expression [72–74] we next sought to identify enhancers directly inhibited by TFAP2 paralogs. To this end we filtered promoter-distal TFAP2A peaks for those where the local H3K27Ac signal was higher in *TFAP2*-KO cells than in WT cells (adj. p <0.05, log2FC>1). Analogously to TFAP2-activated enhancers, TFAP2-inhibited enhancers were split between a subset where the ATAC-seq signal was higher in *TFAP2*-KO cells than in WT cells (as illustrated in **Fig. 2 Box 2A; Fig. 3C, G-G’**), implying TFAP2 paralogs maintain condensed chromatin at these sites, and a subset where it was unchanged (**Fig. 2 Box 2B; Fig. 3D, H-H’**). The first but not the second subset was significantly associated with TFAP2-inhibited genes (**Fig. 3K**, **Table 1**), and the average H3K4me3 signal at the first but not the second subset was higher in *TFAP2*-KO cells than in WT cells (**Fig. 3G’’, H”**). We define these enhancers as TFAP2-pioneered- and-inhibited. The binding site for TFAP2 was strongly enriched for in these enhancers, as were those for ETS1 and CTCF (**Fig. 3O**), both transcriptional repressors [75, 76]. By contrast, the subset of candidate TFAP2-inhibited enhancers where the ATAC- and average H3K4me3 signals were unchanged between TFAP2- KO and WT cells was not associated with TFAP2-inhibited genes. We infer these elements are not, in fact, TFAP2-inhibited enhancers. In conclusion, at TFAP2- inhibited enhancers TFAP2 recruits and/or retains a machinery that condenses chromatin and inhibits enhancer activity; the canonical pioneer factor FOXA1 also has this potential [77, 78]. Alternatively, TFAP2 itself may instead play a distinct role in keeping enhancers closed rather than actively closing them from a presumed prior open state.

We analyzed whether TFAP2 paralogs directly activate or directly inhibit promoters, and how they do so. Although TFAP2A peaks are frequently found at promoters (8277 genes have a TFAP2A peak within 3 kb of the TSS), it was uncommon for the underlying H3K27Ac and H3K4Me3 signals to be elevated (or reduced) in *TFAP2*- KO cells relative to WT cells. Based on this quality, we found 119 candidates for directly TFAP2-activated promoters, and 31 candidates for directly TFAP2-inhibited promoters. Nonetheless, similar to the trends for TFAP2 regulated enhancers, the pioneered subset of directly TFAP2-activated promoters was more strongly associated with TFAP2-activated genes than the non-pioneered subset, and only the pioneered subset of TFAP2-inhibited promoters was associated with TFAP2- inhibited genes (**Supplemental Fig. S11A-L; additional examples in Supplemental Fig. S12A-B**).

Interestingly, at the majority of TFAP2A peaks, whether at open or closed chromatin, neither the ATAC-seq nor H3K27Ac signals were TFAP2-dependent. This large subset of quiescent TFAP2A peaks was associated with neither TFAP2-activated nor TFAP2-inhibited genes (**Table 1, row 10**). In contrast to the quiescent subset of peaks, the set of all TFAP2A peaks was modestly associated with TFAP2-activated genes (**Table 1, row 9**). Nonetheless, we infer that the common practice of using ChIP-seq or CUT&RUN-seq data to identify genes directly regulated by a transcription factor is subject to false positives.

### At a subset of MITF/TFAP2 co-bound peaks, TFAP2 paralogs facilitate MITF binding

A prediction of the TFAP2-as-pioneer-factor model is that binding of transcription factors, like MITF, will depend on TFAP2. Among 36,621 MITF peaks in WT SK-MEL-28 cells that we previously identified by CUT&RUN [17], we found that 15,752 (43%) overlap a TFAP2A peak. Of these, 9,413 (60%) were within open and active (i.e., ATAC+ and H3K27Ac+) chromatin (**Supplemental Fig. S12A**). To assess MITF binding in the absence of TFAP2, we carried out anti-MITF CUT&RUN in *TFAP2*-KO cell lines. Of note, as *MITF* RNA levels in *TFAP2*-KO cells are only about 40-50% of those in WT cells, an across-the-board decrease in the average height (read depth) of MITF peaks was possible. Instead, we observed that the average height of MITF peaks not overlapping TFAP2A peaks was equivalent in *TFAP2*-KO cells and in WT (**Supplemental Fig. S12B** and **Supplemental Fig. S12D** for example at *MLANA*). By contrast, the height of about 35% (5,443 /15,752) of MITF peaks overlapping TFAP2A peaks was significantly lower in *TFAP2*-KO cells than in WT cells (adj. p <0.05, log2FC < -1). (**Fig. 4A-D; Supplemental Fig. S12C-D** and all replicates shown in **Supplemental Fig. S13**). We refer to these as directly TFAP2-activated MITF peaks.

**Figure 4:**
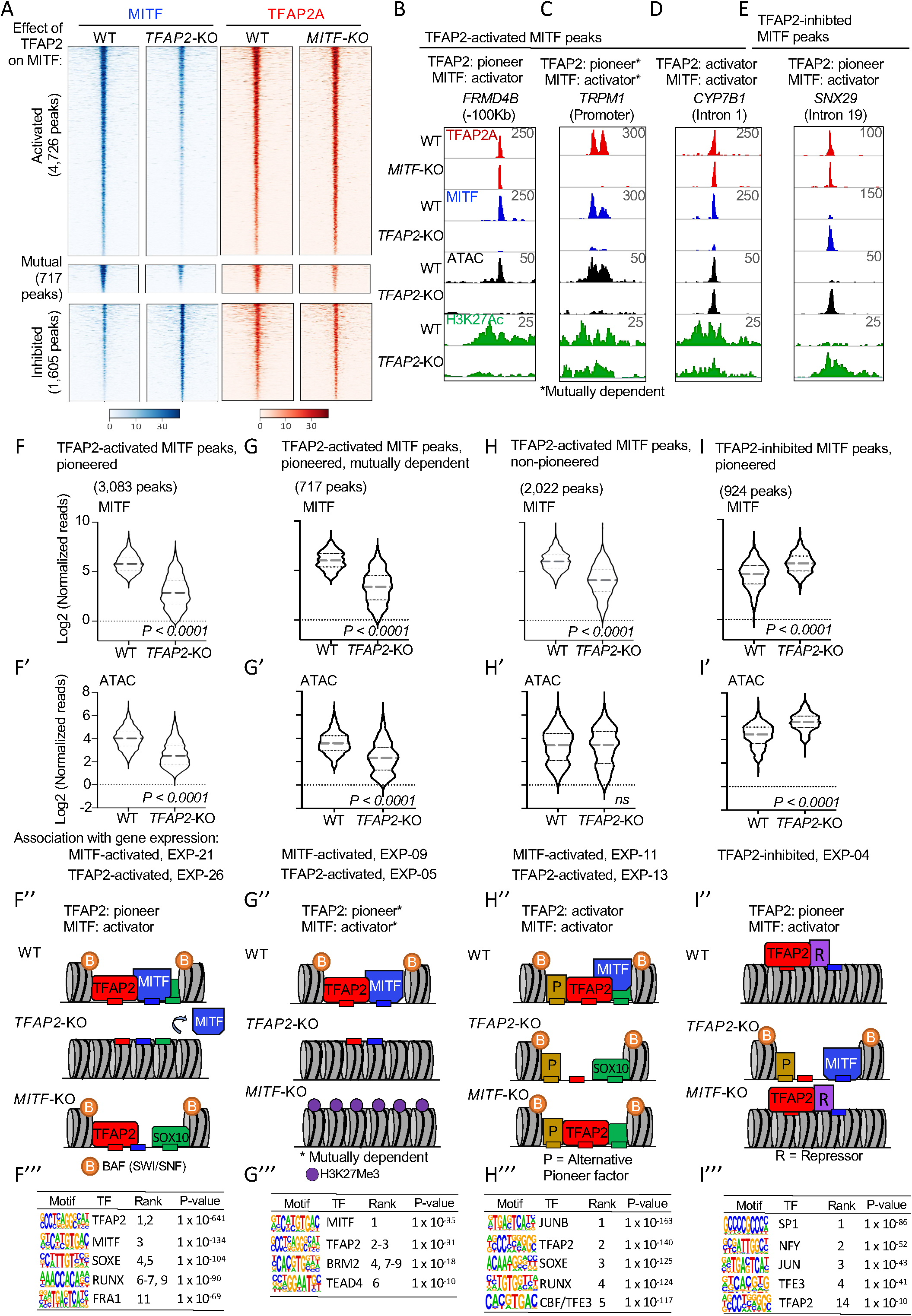
TFAP2 paralogs facilitate chromatin access by MITF. (**A**) Density heatmaps of anti-MITF CUT&RUN-seq in WT and *TFAP2*-KO SK-MEL-28 cells, and anti-TFAP2 CUT&RUN-seq in WT and *MITF*-KO SK-MEL-28 cells at TFAP2- dependent MITF peaks (first cluster), mutually dependent peaks (second cluster) and TFAP2-inhibited MITF peaks (third cluster). Number of peaks in each group as labelled. Regions shown are +/- 3 kb from peak center, normalized reads (RPKM). (**B-E**) Screenshot of IGV genome browser (GRCH37/hg19), showing anti-TFAP2A (red) CUT&RUN-seq in WT and *MITF*-KO cells, and anti-MITF (blue) CUT&RUN-seq, ATAC-seq (black) and anti-H3K27Ac (green) CUT&RUN-seq profiles in WT and *TFAP2*-KO cells. Examples of MITF binding at (**B**) a TFAP2-pioneered-and-activated enhancer (**C**) a TFAP2-pioneered-and-activated promoter (**D**) a non-pioneered TFAP2-activated enhancer and (**E**) a TFAP2-pioneered-and-inhibited enhancer. Genotypes as labeled; y-axis in F applies to F-I, etc. (**F-I’**) Violin plots representing (**F,G,H,I**) anti-MITF CUT&RUN-seq (two independent replicates), and (**F’, G’,H’, I’**) ATAC-seq (four independent replicates) at the indicated number of peaks. P-value according to Student’s t-test, ns; not statistically significant, normalized reads RPKM. Association with gene expression; hypergeometric analysis of TFAP2-depednent and TFAP2-inhibited MITF peaks are shown at TFAP2-activated and MITF-activated genes (FDR < 0.05, log2FC > |1|). **(F’’- I”)** Schematic representation of TFAP2- pioneered-and-activated, TFAP2-activated and TFAP2-inhibited MITF peaks as labelled; B; BAF complex (SWI/SNF), P; alternative pioneer factor. R; repressor protein. Transcription factor binding sites indicated by small rectangles, TFAP2 (red), MITF (blue) and alternative pioneer factor (yellow), example activator SOX10 (green). **(F’’’ – I’’’)** Enrichment of transcription factor motifs using HOMER at (**F’’’**) TFAP2- pioneered -and-activated MITF peaks, (**G’’’**) non-pioneered TFAP2-activated MITF peaks, (**H’’’**) mutually dependent peaks and (**I’’’**) TFAP2-inhibited MITF peaks. TF; transcription factor.

We reasoned that TFAP2 paralogs could facilitate MITF binding by displacing nucleosomes (i.e., in pioneer factor mode) or alternatively by elevating MITF’s affinity for open DNA. Consistent with both models, we observed that directly TFAP2- activated MITF peaks fall in three subsets with respect to the TFAP2-dependence of the underlying ATAC-seq signal. Thus, at about 57% (3,083/5,443) of directly TFAP2-activated MITF peaks the ATAC-seq signal was significantly lower (**Fig. 4B-C, 4F-F’**), at 37% (2,022/5,443) it was unchanged (**Fig. 4D, 4H-H’**), and at 6% it was higher (**Supplemental Fig. S14A-A’**) in *TFAP2*-KO cells compared to WT cells. The first two subsets were strongly associated with TFAP2-activated genes (Hypergeometric test; p-value *=* 8.4 x 10^-26^ and p-value *=* 1.07 x 10^-13^ respectively) and with MITF-activated genes (Hypergeometric test; p-value *=* 1.16 x 10^-21^ and p- value *=* 4.3 x 10^-11^ respectively) (**Fig. 4F’, H’, Table 2**). We infer that at the first subset of TFAP2-dependent MITF peaks, TFAP2 is a pioneer factor (or participates in pioneering function), facilitating access to chromatin for MITF and other transcription factors (illustrated in **Fig. 4F’’**). Supporting this prediction, the transcription factor binding sites for MITF, SOX10, RUNX and FRA1 were strongly enriched at such elements (**Fig. 4F’’’**). At the second subset, TFAP2 is a transcriptional activator that recruits MITF, also functioning as a transcriptional activator; we presume another protein serves as a pioneer factor at this subset (illustrated in **Fig. 4H’’**). Consistent with this notion, the binding site for JUN, a widely deployed pioneer factor [79], site is strongly enriched in these elements (**Fig. 4H’’’**). Examples are shown of TFAP2-activated MITF peaks near *FRMD4B, CYP7B1, TRPM1, SOX9, EDNRB, MREG, GPR143, SNAI2, MEF2C, MYO5A, PAX3, EN1* and *FOXI3* genes (**Fig. 4B-D and Supplemental Fig. S12D**). At the third subset of TFAP2-dependent MITF peaks, where ATAC-seq signal was higher in *TFAP2*-KO cells than in WT cells (**Supplemental Fig. S14A-A’**), TFAP2 may serve as a pioneer factor for MITF in MITF’s proposed role as transcriptional repressor [17] (illustrated in **Supplemental Fig. S14A’’**). However, this category of element was not enriched near genes inhibited by either TFAP2 or MITF (Hypergeometric test; p-value *=* 6.02 x 10^-02^ and p-value *=* 9.12 x 10^-02^ respectively). In summary, these results are consistent with TFAP2 facilitating access for MITF as a transcriptional activator to enhancers in both pioneer-factor and non-pioneer factor modes.

**Table 2:** TFAP2-dependent MITF peaks and gene expression (Fisher’s Exact Test; hypergeometric analysis):

| Effect of TFAP2 on |  |  | Association with<br>activated genes | Association with<br>inhibited genes |
| --- | --- | --- | --- | --- |
| MITF binding | Chromatin | N (# of<br>elements) | p-value<br>\|Log2FC\| >1 | p-value<br>\|Log2FC\| >1 |
| All TFAP2 - activated | Any effect<br>or none | 5,443 | $8.4 \times 10^{-26}$ | ns |
| Pioneered | Opens | 3,083 | $1.6 \times 10^{-26}$ | ns |
| Non-pioneered | None | 2,358 | $1.07 \times 10^{-13}$ | ns |
| Mutually - activated | Opens | 717 | $3.47 \times 10^{-05}$ | ns |
| All TFAP2 - inhibited | Any effect<br>or none | 1,605 | ns | $5.0 \times 10^{-03}$ |
| Pioneered | Condenses | 924 | ns | $3.5 \times 10^{-04}$ |
| Non-pioneered | None | 681 | ns | ns |
| None | Any effect<br>or none | 10,418 | ns | ns |

Column 1: groups of directly TFAP2-regulated MITF peaks (activated or inhibited), determined by anti-MITF CUT&RUN in *TFAP2*-KO and WT SK-MEL-28 cell lines. Column 2: TFAP2-dependent MITF peaks at TFAP2-regulated nucleosome depleted regions (ATAC-Seq; opened or condensed by TFAP2). Column 3: the number of directly TFAP2-regulated MITF peaks. Column 4-7: The effect of TFAP2-regulated MITF peaks on gene expression (two independent clones, 4 replicates each) and WT (4 replicates). Differentially expressed genes (adjusted p-value <0.05) between *TFAP2*-KO cells and WT cells was determined by RNA-Seq. The p-value from the Fisher's Exact Test in which the null hypothesis is that no association exists. DEG: differentially expressed genes. Attention should be focused on columns 4-5, which indicate the strength of association between the regulation status of a gene and the TFAP2-regulated MITF peaks of interest; significant associations are highlighted yellow. ns; not statically significant.

To test whether depended on MITF, we carried out anti-TFAP2A CUT&RUN in *MITF*-KO cells. *TFAP2A* mRNA and protein levels were equivalent in *MITF*-KO and WT cells (**Supplemental Fig. S9E**), and the average TFAP2A peak height was globally equivalent as determined by CUT&RUN. At 13% (717/ 5334) of TFAP2- dependent MITF peaks, the TFAP2A peak was, reciprocally, significantly reduced in *MITF*-KO cells (**Fig. 4A and 4C**). At such loci, the average ATAC-seq signal was reduced in *TFAP2*-KO cells as compared to WT cells (**Fig. 4C, 4G-G’**). We termed these peaks mutually-dependent (i.e. mutually-activated peaks) (**illustrated in Fig. 4G’’**). Interestingly, mutually-dependent MITF/ TFAP2 peaks were enriched in binding motifs for TFAP2, MITF, BRM2 and TEAD4 but, unlike the other subsets of TFAP2-activated MITF peaks, not for SOXE (**Fig. 4H’’’**). SOX10 co-binds many loci together with MITF [15] and if SOX10 is absent from mutually-dependent peaks this may explain the dependence of TFAP2A binding on MITF at these sites. At ∼40% (288/ 717) of the mutually dependent peaks, and in the gene body surrounding these peaks, the repressive histone mark H3K27Me3, known to be applied by the polycomb complex [69, 80], was significantly higher in *MITF*-KO cells relative to WT cells but, unexpectedly, not in *TFAP2*-KO cells relative to WT cells, even though MITF binding was lower in *TFAP2*-KO cells (illustrated in **Fig 4G’’, Supplemental Fig. 12E-G**).

In summary, the binding of MITF depends on TFAP2 at about one third of MITF peaks that overlap TFAP2A peaks. Such TFAP2-activated MITF binding occurs both at loci where nucleosome packing depends on TFAP2 (pioneer factor mode) and where it does not (non-pioneer factor mode). At a subset of the former, but none of the latter, TFAP2A binding is, reciprocally, MITF-dependent.

### At a subset of MITF/TFAP2A co-bound peaks, TFAP2 paralogs inhibit chromatin access for MITF

In Figure 3 we established that at some TFAP2A peaks, TFAP2 paralogs maintain closed chromatin, and presumably inhibit binding of transcription factors. Consistent with this prediction, among MITF peaks overlapping TFAP2A peaks, 10% (1,605) of the MITF peaks were higher in *TFAP2*-KO cells than in WT cells (third cluster **Fig. 4A, E; Supplemental Fig. S13**), implying TFAP2 paralogs inhibit binding of MITF at this subset. At most peaks within this subset (58%, 924/1,605), the ATAC-seq signal was also significantly higher in *TFAP2*-KO cells versus WT cells (violin plots, **Fig. 4I, I’,** illustrated in **Fig. 4I’’**), implying TFAP2 maintains condensed chromatin at this subset. DNA elements underlying this subset were enriched for transcription factor binding sites including those for SP1, NFY, JUN and TFE3 (**Fig. 4I’’’**). Moreover, these elements were modestly associated with TFAP2-inhibited genes (**Table 2;** row 8).

Of note, at the majority of MITF peaks that overlap TFAP2A peaks (65%, 10,418/ 15,752), the height of the MITF peak was equivalent in *TFAP2*-KO and WT cells (TFAP2-independent MITF peaks) (**Supplemental Fig. S13, Supplemental Fig. S15A;**), implying TFAP2 paralogs neither recruit nor repel MITF at these sites. Interestingly, such TFAP2-independent MITF peaks were not strongly enriched for the TFAP2 binding site (**Supplemental Fig. S15A**), implying that TFAP2 is attracted to many of these sites via other proteins rather than binding directly to the DNA. Such indirect binding may be less avid, as the average height of TFAP2-independent MITF peaks was smaller than that of TFAP2-dependent MITF peaks (**Supplemental Fig. S13,** compare MITF signal in first and fourth cluster WT cells). As expected, TFAP2-independent MITF peaks were associated neither with TFAP2-activated nor TFAP2-inhibited genes (**Table 2**; row 10).

### TFAP2 and MITF co-regulate genes in the melanocyte gene regulatory network

The delayed pigmentation in zebrafish *tfap2a/tfap2e* double mutants, and the reduced expression of many pigmentation genes in *TFAP2*-KO cells, is consistent with two scenarios which are not exclusive of one another. In the first, TFAP2 paralogs directly activate *MITF* expression and thereby indirectly activate expression of pigmentation genes. In the second, TFAP2 paralogs directly activate expression of pigmentation genes. Supporting the first scenario, there is a TFAP2-pioneered-and-activated enhancer in intron 2 of the *MITF* gene (**Supplemental Fig. S16**), and *MITF* mRNA levels are about 40-50% lower in *TFAP2*-KO cells than in WT cells. However, arguing against a strong role for this mechanism, many of the genes whose expression was most strongly reduced in *MITF*-KO cells compared to WT cells were completely TFAP2-independent, or were TFAP2-inhibited (**Supplemental Table 4**). To assess the second scenario, we identified the set of genes activated directly by MITF, defined as MITF-activated genes associated with an MITF peak, and the set of genes directly activated by TFAP2, defined as TFAP2-activated genes associated with an TFAP2-activated enhancer (i.e., of pioneered or non-pioneered variety). Supporting the second mechanism, genes activated directly both by TFAP2 and by MITF were enriched for GO terms related to pigmentation and pigment cell differentiation (**Fig. 5A-B**) [81]. Of note, genes related to pigmentation were also among those apparently directly regulated solely by MITF or TFAP2 paralogs (**Fig. 5C**). We took a similar approach to identify genes directly inhibited by TFAP2 and/or by MITF (**Fig. 5D**). Genes directly inhibited by both were strongly enriched for GO terms related to cell motility and cell migration (**Fig. 5D-E**). Additional categories of genes strongly activated or inhibited by TFAP2 included genes associated with cell proliferation and cell differentiation (**Fig. 6A; Supplemental Fig. S17A-B**) and with cell-cell adhesion, respectively (**Fig. 6B; Supplemental Fig. S17C)**.

**Figure 5:**
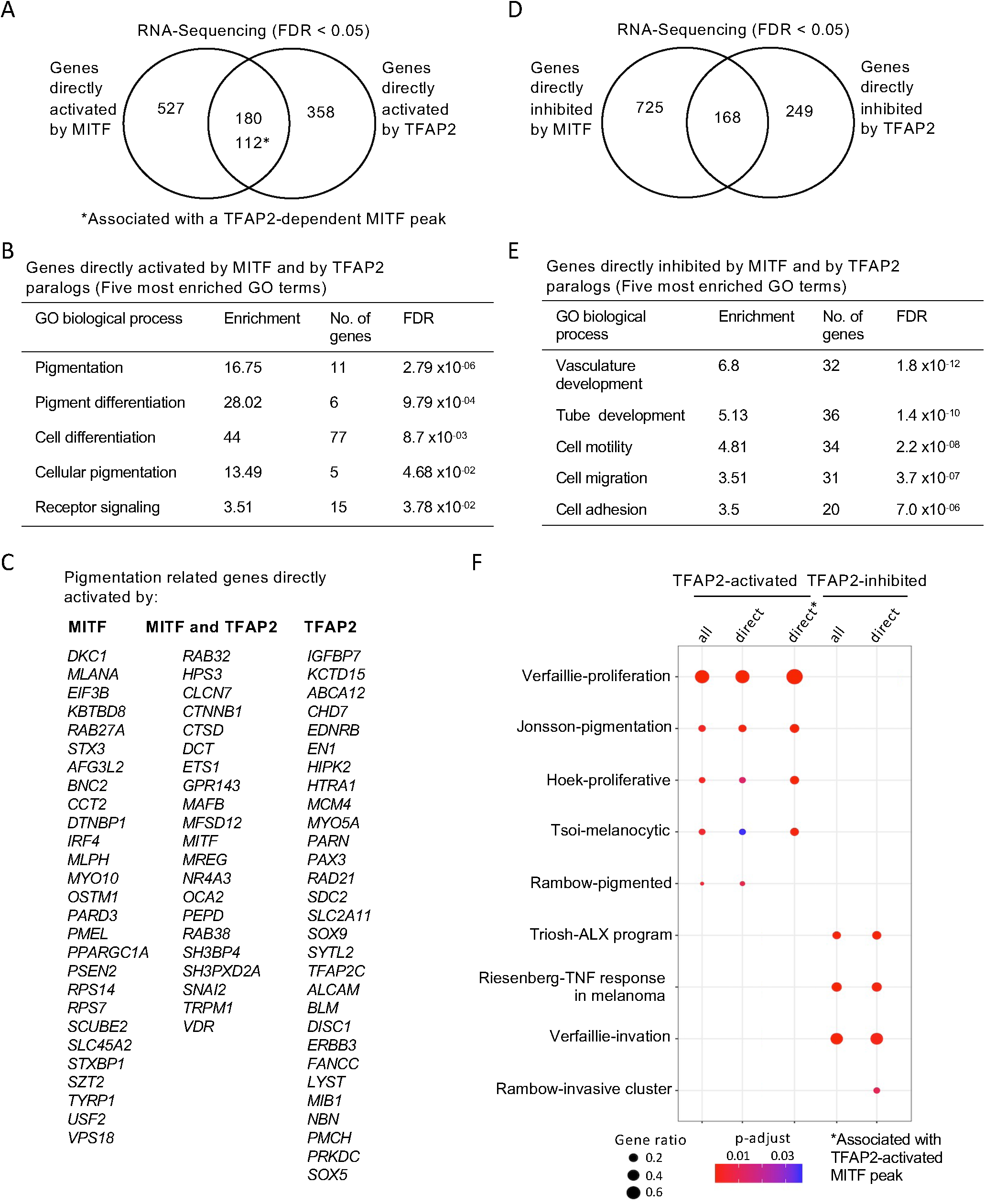
TFAP2 and MITF co-regulate pigmentation and cell differentiation genes in SK-MEL-28 cell lines. (**A**) Venn diagram representing the overlap of genes apparently directly activated by MITF (i.e., expression lower in *MITF*-KO than in WT cells, FDR < 0.05, and an MITF peak within 100kb of the TSS) and genes directly activated by TFAP2 (i.e., expression lower in *TFAP2*-KO than in WT cells, FDR < 0.05, and an TFAP2-activated enhancer of any category within 100kb of the TSS). The number of overlapping genes with TFAP2-dependent MITF peaks are also shown (*). (**B**) Gene ontology (GO) biological process analysis enriched among MITF- and TFAP2-activated genes are shown (Top 5 hits). (**C**) A curated list of pigment-associated genes [118] was intersected with directly MITF-activated, overlapping genes of directly MITF- and TFAP2-activated genes, and directly TFAP2-acitvated genes and represented by gene list. (**D**) Venn diagram representing directly MITF inhibited genes (MITF peak within 100kb of a TSS), based on RNA-seq, in *MITF*-KO versus WT cells (FDR < 0.05) and directly TFAP2 inhibited genes (TFAP2-inhibited enhancers, of any category, within 100kb of a TSS), based on RNA-seq, in *TFAP2*-KO versus WT cells (FDR < 0.05). (**E**) Gene ontology (GO) biological process analysis enriched among MITF- and TFAP2- inhibited genes are shown (Top 5 most enriched GO terms). GO analysis was performed using PANTHER. (**F**) Dot plot of enrichment analysis showing the enrichment of melanoma gene signatures from the literature in directly TFAP2- activated and TFAP2-inhibted genes. P value is red lowest to blue highest; gene ratio is the fraction of all genes in the gene signature category that are included in the set identified here. TFAP2-activated genes associated with TFAP2-dependent MITF peaks are shown (*).

**Figure 6:**
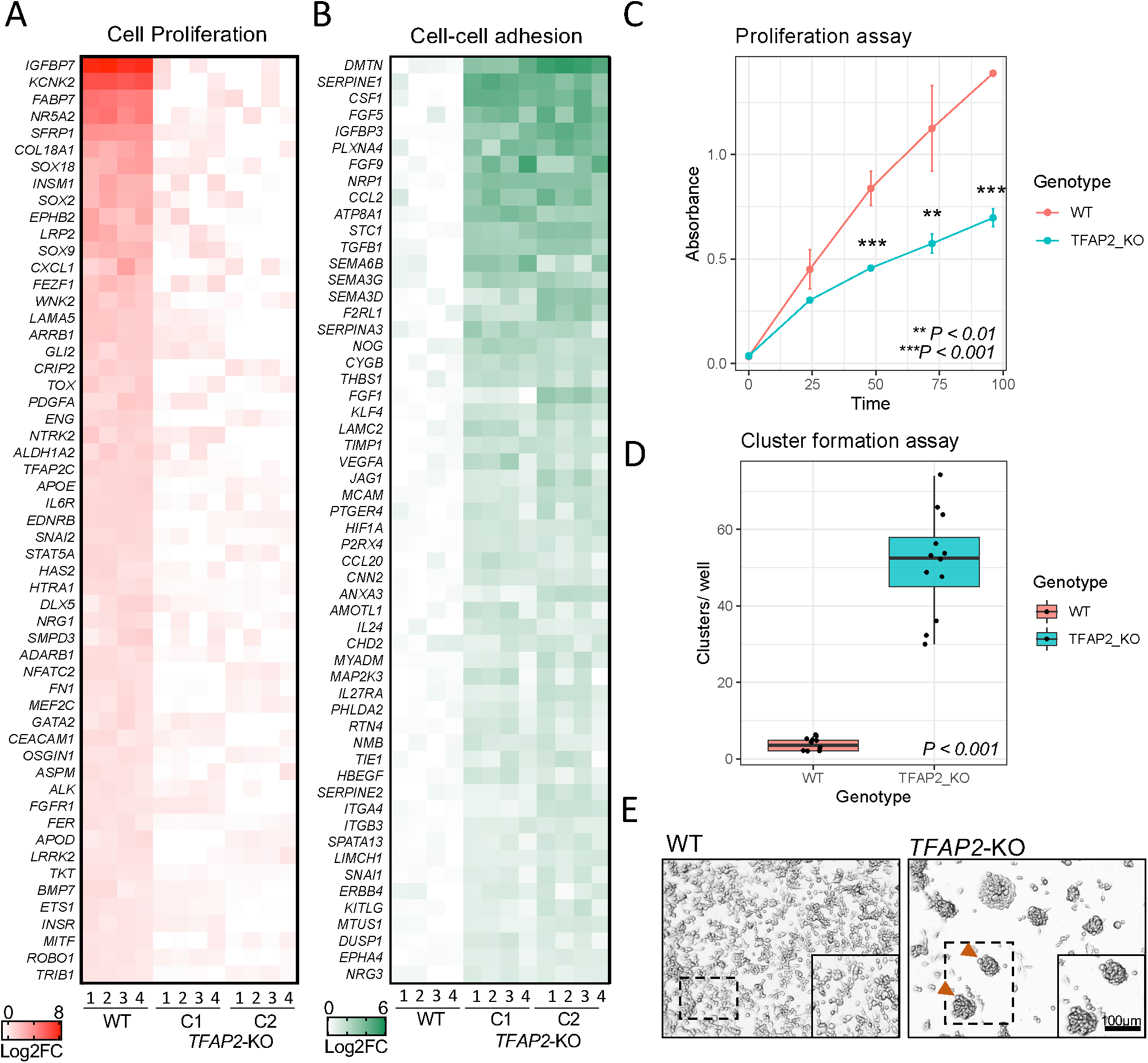
TFAP2 paralogs promote cell proliferation and inhibit cell-cell adhesion a melanoma cell line. (**A**) Heatmap representing the top 55 directly TFAP2-activated genes and (**B**) the top 55 directly TFAP2-inhibited genes that are associated with the GO terms cell pigmentation and cell-cell adhesion, respectively. Four replicate RNA-Seq experiments are shown for WT cells and two clones of *TFAP2*-KO cells (Clone 2.12 and Clone 4.3) (FDR < 0.05). (**C**) Growth curves (mean ± SE of mean) over 100 hours of cultivation for WT and *TFAP2*-KO SK-MEL-28 cells. x-axis is time and y-axis is absorbance at 450nm which is directly proportional to number of living. (**D**) Box plots representing the quantification of cluster formation on low-bind plates after 72 hours of culture (n = 12 independent experiments, p < 0.001 by Student’s t-test, plot shows mean ± SD). (**E**) Representative images of clusters formed in WT and *TFAP2*-KO cells after 72 hours.

We next examined the association of TFAP2-activated and TFAP2-inhibited genes with gene expression profiles from melanoma tumors and cell lines with distinct phenotypes [82–88]. Enrichment analysis showed that melanoma profiles previously found to be associated with high levels of MITF activity [89] were enriched for genes directly activated by TFAP2, including the subset associated with TFAP2-dependent MITF peaks (**Fig. 5F**). Moreover, melanoma profiles associated with low levels of MITF activity were enriched for genes directly inhibited by TFAP2 (**Fig. 5F**). These findings support other observations that the level of TFAP2 expression has a profound effect on the melanoma phenotype [32, 90].

Finally, we performed cell proliferation, cell adhesion and cell migration assays on wild-type and TFAP2-depleted SK-MEL-28 cells. Consistent with effects on gene expression, *TFAP2*-KO cells showed reduced proliferation over 24, 48, 72 and 96 hours compared to WT cells (**Fig. 6E**), and in cluster formation assays, *TFAP2*-KO cells showed increased cell-cell adhesion compared to WT cells (**Fig. 6E**). However, in contrast to the effects on gene expression, while WT cells migrated to fill in a scratch within 24 hours, neither *TFAP2*-KO clone filled it within that time (**Supplemental Fig. S18A**). This finding also contrasts with the observation that the expression of *tfap2e* correlates negatively with the migratory capacity of zebrafish models of melanoma [32], but it is consistent with the accumulation of melanocytes in the dorsum of zebrafish *tfap2a* knockout embryos [23, 63, 64] and *tfap2a/ tfap2e* double mutant embryos. It is noteworthy that in *MITF*-KO cells, similar to in *TFAP2*- KO cells, cell migration genes are upregulated, but cell migration *in-vitro* is inhibited [17]. In summary, reduced expression of pigmentation, differentiation and cell growth genes and elevated expression of cell adhesion and cell invasion genes in *TFAP2*- KO cells as compared to WT cells is largely explained by the direct effects of TFAP2 paralogs on these genes.

## Discussion

Here, single-cell seq analysis of zebrafish embryos indicated that expression of MITF-target genes encoding regulators of pigmentation occurs only in the subset of *mitfa*-expressing cells which also express *tfap2* paralogs. In embryos with loss-of-function mutations in *tfap2a* and *tfap2e*, the two *tfap2* paralogs with highest expression in melanophore precursors, there are fewer embryonic melanophores than in wild-type embryo’s and pigmentation is delayed. The phenotype was similar, but less penetrant than the phenotype in *tfap2a* mutants injected with *tfap2e* MO [10]. We speculate that homeostatic upregulation of *tfap2c* in *tfap2a/tfap2e* double mutants, possibly mediated by transcriptional adaptation [91], suppresses the phenotype. Simultaneous deletion of *tfap2a* and *tfap2c* eliminates neural crest precluding the possibility of examining melanocyte development in such embryos [23, 29, 64]. It has been thus far impossible to evaluate the phenotypic consequence of loss, rather than just reduction, of Tfap2 activity in the melanophore lineage in zebrafish. Although RNA-less loss-of-function mutants of *tfap2a* and *tfap2e* should prevent transcriptional adaptation [91, 92], it is possible the homeostatic upregulation of *tfap2c* in the existing mutants documented here occurs by another mechanism. Of note, in mice with simultaneous tissue specific deletion of Tfap2a and Tfap2b in the neural crest lineage the number of embryonic melanocytes is reduced relative to wild-type [33]. Nonetheless, because a) MITF is known to directly activate expression of pigmentation genes and cell cycle genes, b) TFAP2A and MITF bind many of the same loci in melanocytes [33], and c) there is prior evidence that TFAP2 paralogs can serve as pioneer factors [41, 42, 45], we hypothesized that TFAP2 is a pioneer factor that facilitates chromatin access for MITF.

To test the TFAP2-pioneer-factor model, the two paralogs with highest expression, TFAP2A and TFAP2C, were deleted from the SK-MEL-28 cells. Creation and integration of several genomic datasets permitted us to identify genomic elements that were bound by TFAP2A in WT cells and that either lost or gained H3K27Ac signal in *TFAP2*-KO cells; we inferred that these elements were enhancers directly activated or inhibited by TFAP2. As expected by the TFAP2-as-pioneer-factor model, at a subset of TFAP2A-activated enhancers, TFAP2 paralogs were required to keep chromatin open, and at all TFAP2-inhibited enhancers it was required to keep it closed. We also showed that TFAP2 paralogs facilitated binding of another transcription factor (MITF), that this occurred at loci where TFAP2 facilitates chromatin access, and that this dependency was not reciprocal (at most loci bound by both transcription factors). These findings imply that TFAP2 paralogs recruit distinct transcription factors to distinct categories of enhancer, that these in turn recruit enzymes that open chromatin or that condense it (TFAP2 paralogs also may directly recruit such enzymes) [93], and that the state of chromatin accessibility determines access of other transcription factors. Therefore, TFAP2 paralogs facilitate chromatin access by other transcription factors, either as a pioneer factor or as a linker that connects pioneer factors to chromatin-remodeling machinery. Notably, the role of MITF at the co-bound enhancers is still largely unexplored.

Of note, there are loci where the pioneer factor FOXA1 appears to recruit proteins like GRG3 (groucho-related gene 3) that condense chromatin [77, 78]. At TFAP2- inhibited enhancers, our data do not distinguish between whether TFAP2 binds open loci and closes them or instead binds to closed loci and helps stabilize the closed state. Of note, the majority of TFAP2A peaks did not coincide with enhancers either activated or inhibited by TFAP2 paralogs and were thus quiescent. In some cases, such apparently quiescent TFAP2A peaks may represent loci that TFAP2 paralogs have stably pioneered; a precedent for this scenario is that at a subset of elements pioneered by PAX7, chromatin remains open after the removal of PAX7 [94]. To explore this possibility, TFAP2A and ATAC-seq will need to be monitored in a developmental time series, or in TFAP2-KO cells before and after TFAP2A has been re-expressed.

Like other pioneer factors, TFAP2 can activate enhancers in a non-pioneer factor mode. At a subset of TFAP2A-activated enhancers the H3K27Ac signal, but not the ATAC-seq signal, was lower in TFAP2-KO cells than in WT cells. Evidence that such elements are indeed TFAP2-activated enhancers is that their average H3K4Me3 signal was also lower in TFAP2-KO cells than in WT cells. We infer that TFAP2 activates these enhancers, but not as a pioneer factor. At such enhancers the continued presence of TFAP2 is necessary for continued acetylation of histone H3 lysine 27 (H3K27Ac), which fits with evidence that TFAP2 binds the histone acetyl transferase p300/CBP [95] and inhibits the NURD histone-deacetylase complex [93]. TFAP2 may attract other transcription factors without affecting nucleosome positioning; indeed, some TFAP2-dependent MITF peaks were found at non- pioneered TFAP2-activated enhancers. The fact that the TFAP2 binding site is not strongly enriched in non-pioneered TFAP2-activated enhancers implies TFAP2 binds these elements indirectly. Although we refer to these elements as non-pioneered TFAP2-activated enhancers, our experimental design could not rule out the possibility that some such enhancers were stably pioneered by TFAP2 as discussed above. However, the observation that these sites are less enriched for the TFAP2 binding site than the TFAP2-pioneered-and-activated enhancers, and that the binding sites of pioneer factors FOS and JUN are enriched [96], supports the alternative model that such elements are simply pioneered by different transcription factors.

Interestingly, at a subset of MITF/TFAP2A overlapping peaks where MITF was lost in *TFAP2*-KO cells, TFAP2A binding was lost in *MITF*-KO cells (MITF/TFAP2A mutually dependent peaks). There is precedent for reciprocal dependence of the binding of pioneer factors, in the cases of both FOXA1 and steroid hormone receptors, at subsets of sites where they are co-bound [97]. Notably, at many MITF/TFAP2A mutually dependent peaks, the repressive mark H3K27Me3 accumulated in MITF-KO cells. This is consistent with evidence that the SWI/SNF complex, which MITF probably recruits to such loci, competes for access to chromatin against the H3K27Me3-depositing Polycomb repressor complex and that the binding of pioneer factors is impeded by condensed H3K27Me3-positive chromatin [80, 98]. We propose that at the majority of TFAP2-dependent MITF peaks, but not at MITF/TFAP2A mutually dependent peaks, a measure of BRG1 binding is retained in MITF-KO cells, possibly recruited by another activator like SOX10, but this is not the case for MITF/TFAP2A mutually dependent peaks.

Finally, by identifying enhancers regulated by TFAP2, whether in pioneer mode or not, we were able to identify with high-confidence genes directly activated and directly inhibited by TFAP2. The genes directly activated by TFAP2, including by the specific mechanism of facilitating MITF access at a nearby enhancer, were enriched in GO terms related to pigmentation and proliferation, and overlapped significantly with expression profiles previously associated with high levels of MITF activity [82–88]. The MITF “rheostat model” predicts that high MITF activity promotes cell proliferation [2, 12]. Proliferation of both *MITF*-KO [17] and *TFAP2*-KO cells (shown here) is slower than of the wild-type counterparts. We suggest there are two mechanisms to explain the shared expression profiles and phenotypes between *TFAP2*-KO cells and *MITF*-KO cells. First, the expression of *MITF* mRNA is about 40-50% lower in *TFAP2*-KO than WT cells. It is unclear how important this effect is because the average read depth of MITF peaks was equivalent in WT and TFAP2- KO cells, in particular at MITF-activated genes that are not also TFAP2-activated (e.g., *MLANA*). Second, TFAP2 facilitates binding of MITF to enhancers and promoters, leading to higher expression of associated MITF-target genes. The “rheostat model” also predicts that low MITF activity promotes cell invasion [12], and there was a large overlap of TFAP2-inhibited and MITF-inhibited genes and these genes were enriched for those involved in cell-cell adhesion. We found evidence that TFAP2 directly inhibits enhancers, but this did not involve TFAP2 and MITF cooperativity. Directly-TFAP2-inhibited genes were associated with cell-cell adhesion and *TFAP2*-KO cells formed large clusters when cultured on low adhesive plates. Interestingly, in zebrafish melanoma, low *tfap2e* expression correlates with increased metastases [32]. We have previously shown that MITF binds to enhancers near MITF-inhibited genes [17]. However, mechanisms by which MITF inhibits gene expression in melanoma cells and *in-vivo* remain to be elucidates. Given that an MITF-low is a deadly status for melanoma, an interesting possibility for therapy would be to effectively covert them to MITF-high by manipulating pioneer factors or other MITF-cofactors. In summary, MITF activity in melanoma cells – and thus the phenotypes of these cells – depends in part on the presence of transcription factors that give MITF access to specific regulatory elements.

## Materials and Methods

### Zebrafish lines and maintenance

D. rerio were maintained in the University of Iowa Animal Care Facility according to a standard protocol (protocol no. 6011616). All zebrafish experiments were performed in compliance with the ethical regulations of the Institutional Animal Care and Use Committee at the University of Iowa and in compliance with NIH guidelines. Zebrafish embryos were maintained at 28.5 C and staged by hours or days post-fertilization (hpf or dpf).

### Single cell suspension, fluorescence activated cell sorting and library preparation

50 Tg(*mitfa*:GFP) transgeneic zebrafish embryos [99] were collected at 28 hours post fertilization (hpf) and washed in PBS without Ca2+ or Mg2+ (Life technology). Embryos were manually dechorionated under a dissecting scope (Leica KL300 LED), washed twice in 1X PBS and then cell dissociated cells using a pestle and incubated in trypsin (0.25%)-EDTA (Life technology) at 33°C for 30 minutes (ensuring to pipette mixed every 5 minutes). Reactions were quenched by adding PBS supplemented with 5% fetal bovine serum (FBS, Life technologies). Dissociated cells were centrifuged at 500 g for 5 minutes and cell pellets were washed with PBS- 5%FBS before passing thought a Bel Art™ SP Scienceware™ Flowmi™ 40 μm cell strainers (ThermoFisher Scientific). Dissociated cells were re-suspended into single-cell solution and analyzed at the University of Iowa Flow Cytometry Facility, using an Aria Fusion instrument (Becton Dickinson, Franklin Lakes, NJ).

### Single-cell RNA-sequencing (10x genomics) and data processing

The sorted cellular suspensions were loaded on a 10x Genomics Chromium instrument to generate single-cell gel beads in emulsion (GEMs). Approximately 10,000 cells were loaded per channel. Single-cell RNA-Seq libraries were prepared using Single Cell 3′ Reagent Kits v2: Chromium™ Single Cell 3′ Library & Gel Bead Kit v2, PN-120237; Single Cell 3′ Chip Kit v2 PN-120236 and i7 Multiplex Kit PN-120262 (10x Genomics) following the Single Cell 3′ Reagent Kits v2 User Guide (Manual Part # CG00052 Rev A). Libraries were run on an Illumina HiSeq 4000 as 2 × 150 paired-end reads. Sequencing results were demultiplexed and converted to FASTQ format using Illumina bcl2fastq software.

A custom reference genome for the Zebrafish was constructed with GRCz11 primary assembly (Ensembl) using Cell Ranger (Cell Ranger mkref function). 10X Genomics scRNA-seq reads were then processed and aligned to this reference with Cell Ranger (Cell Ranger Count function). Further analysis and visualization was performed using Seurat (v4.1.0) [51], cells with fewer than 200 RNA feature counts and greater than 5% mitochondrial contamination were removed with filtering. RNA counts were normalized, and FindVariableFeatures was run with the following parameters: selection.method = “vst”, nfeatures = 2,000. The cells were originally clustered in a UMAP by RunPCA then RunUMAP with dims 1:30. FindNeighbors was run with dims 1:30 followed by FindClusters using a resolution of 1.2. Pigment clusters were further analyzed by sub-setting out and re-clustering using dims 1:30, which identified 10 clusters. Pseudotime was performed using Monocle3 (v0.2.3.0) [100].

### Generation of a zebrafish *tfap2e* loss-of-function allele

To generate the tfap2e loss-of-function allele, we designed paired (e.g., left and right) zinc finger nucleases (ZFN) targeting exon 2 of the tfap2e locus resulting in non-homologous end-joining and disruption of the open reading frame for Tfap2e. Briefly, the online tool, ZiFiT [101], was used to identify an optimal ZFN target site [utilizing the CoDa approach [102]. Once identified, a custom DNA fragment encoding the entire left or right zinc finger array (ZFA) along with flanking XbaI and BamHI restriction sites was synthesized (Integrated DNA Technologies, Coralville, IA). Subsequently, the ZFA fragment was subcloned into pMLM290 (Addgene, plasmid 21872), which includes a modified FokI nuclease domain [103]. Next, the fully assembled ZFN was PCR amplified, directionally cloned into pENTR-D/TOPO (ThermoFisher Scientific), and finally subcloned into pCS2+DEST using Gateway LR Clonase II enzyme mix (ThermoFisher Scientific). Once assembled, the final pCS2+ plasmids were sequence verified, linearized, mRNA synthesized in vitro (mMessage mMachine SP6 Kit, Ambion/ThermoFisher Scientific). Synthesized RNA was cleaned using the Qiagen RNeasy Kit (Qiagen) and both left and right ZFN components were co-injected into 1-cell stage zebrafish embryos. Following injections, embryos were initially screened via PCR and restriction enzyme digest to confirm editing at the target site. Upon confirmation, additional embryos from a similar clutch (F0’s) were allowed to develop into adulthood, ‘mosaics’ identified and out-crossed, and a stable F1 generation isolated.

### Cell lines, reagents, and antibodies

The cells referred to as WT throughout the document are the parent SK-MEL-28 (HTB-72) line. They and the derivative line, delta6-MITF knockout cells (referred to as MITF-KO cells in this work), were obtained from the laboratory of Dr. Eirikur Steingrimsson. The cells were grown in RPMI 1640 medium (Gibco #5240025) supplemented with 10% FBS (Gibco #10270106) at 5% CO_2_ and 37°C. Cells were tested for, and determined to be free of, mycoplasma. SK-MEL-28 cells harbor the BRAF^V600E^ and p53^L145R^ mutations [104]. The following primary antibodies and their respective dilutions were used in western blotting (WB) and CUT&RUN experiments: anti-Tubulin (Sigma, #T6199), 1:5000 (WB); anti-MITF (Sigma, #HPA003259), 1:2000 (WB), 1:100 (CUT&RUN); anti-TFAP2A (Abcam, ab108311), 1:5000 (WB), 1:200 (CUT&RUN); anti-TFAP2C (Santa-Cruz #SC-12762 X), 1:1000 (WB); anti- H3K27Ac (EMD Millipore, #07-360), 1:100 (CUT&RUN); anti-H3K4Me3 (EMD Millipore, #05-745R), 1:100 (CUT&RUN); H3K27Me3 (EMD Millipore, #07-449), 1:100 (CUT&RUN); Rabbit IgG (EMD Millipore,#12-370), 1:100 (CUT&RUN); Mouse IgG (EMD Millipore, #12-371), 1:100 (CUT&RUN). The following secondary antibodies and their respective dilutions were used: Anti-mouse IgG(H+L) DyLight 800 conjugate (Cell Signaling Technologies, #5257), 1:20000; and anti-rabbit IgG(H+L) DyLight 680 conjugate Cell Signaling Technologies, #5366), 1:100. Images were captured using an Odyssey CLx Imager (LICOR Biosciences).

### Purification of pA/G-MNase

*E.coli* strain BL21-DE3 was transformed with plasmid DNA pAG-MNase-6xHis (Addgene, plasmid #123461). Recombinant pAG-MNase was purified from cells grown in LB medium to OD600 0.6 at 37°C. Cells were induced with 0.5 mM IPTG and cultured for 16 hours at 20°C. Cell pellets were homogenized in lysis buffer (10 mM Tris, pH 7.5, 300 mM NaCl, 10 mM imidazole) containing lysozyme and protease inhibitors, then sonicated and the slurry was cleared by centrifugation (35K RPM, Ti70 rotor). The supernatant was subjected to IMAC chromatography (NI-NTA column) and to size-exclusion fractionation (Superdex 75) using a BioLogic DuoFlow QuadTec FPLC system (Bio-Rad). The purified pAG-MNase was concentrated by buffer exchange with ultrafiltration (Amicon Ultra-15, 10K). Finally, the purified pAG-MNase was diluted in dilution buffer (10 mM Tris pH7.5, 120 mM NaCl, 0.01mM EDTA, and 50% glycerol), and stored at -80°C.

### Generation of *TFAP2A*; *TFAP2C* knockout cell lines (*TFAP2*-KO)

*TFAP2*-KO clones were generated using the Alt-R CRISPR-Cas9 technology from Integrated DNA Technologies (IDT). Briefly, crRNAs targeting exon 2 of *TFAP2A* and *TFAP2C* were designed using the Cas9 guide RNA design checker tool (crRNA sequences below). Equimolar concentrations of crRNA and tracrRNA (IDT, #1072532) were annealed to form gRNA complexes. The ribonucleoprotein (RNP) complex was prepared by mixing gRNAs and Cas9 protein (IDT #1081058). SK-MEL-28 cells were transfected with constructs encoding components of RNP complexes using the Lipofectamine CRISPRmax Cas9 transfection reagent (ThermoFisher #CMAX00015) following the manufacturer’s protocol. Single-cell colonies were screened by PCR and Sanger sequencing using primers flanking the cut sites (primer sequences below). Mutant clones (clone 2.12 and clone 4.3) were selected and further screened by western blotting, using anti-TFAP2A and anti- TFAP2C antibodies. The control cell lines used in this study were generated following this protocol but without adding gRNA duplexes.

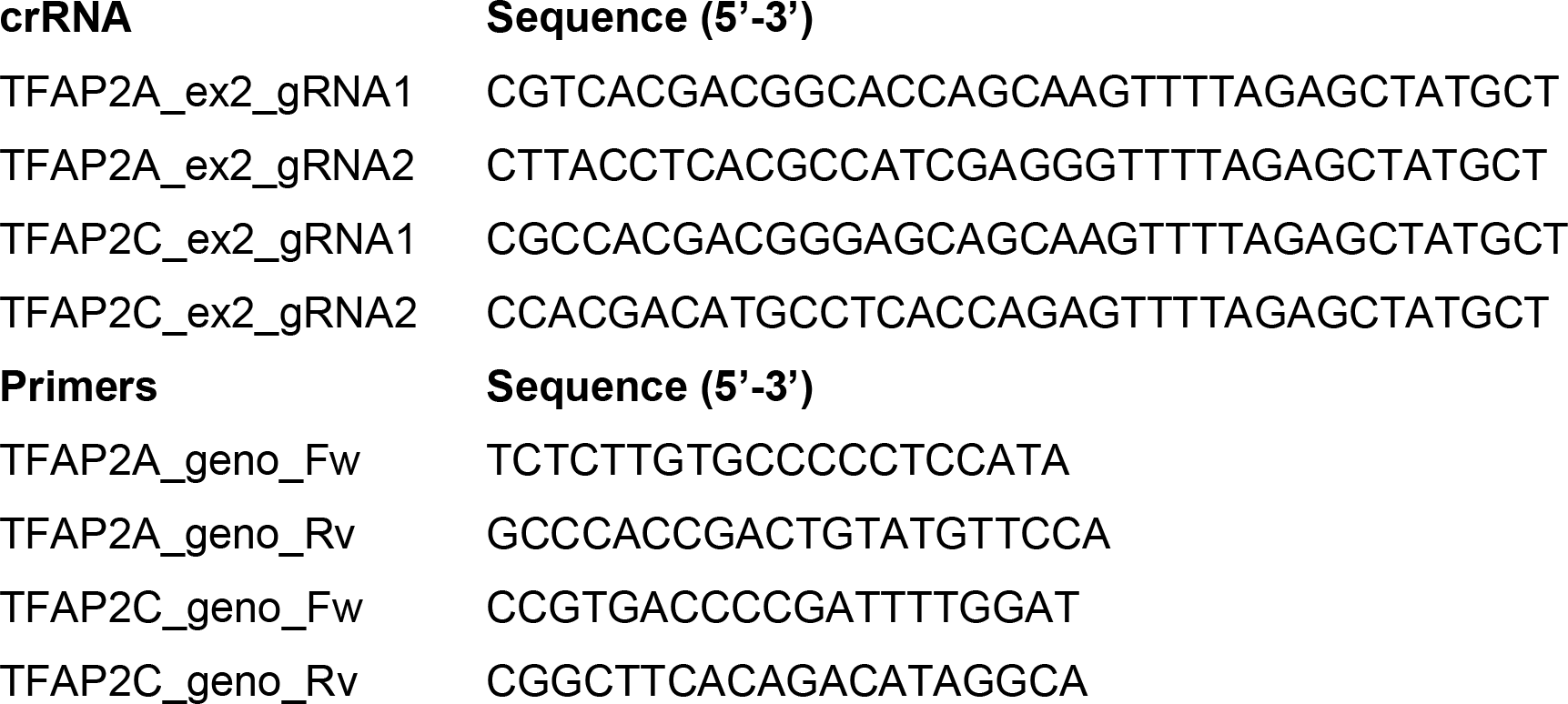

### SDS-PAGE and Western blotting

*TFAP2*-KO and WT cells were washed in ice-cold PBS. RIPA buffer containing protease inhibitors (Roche, cOmplete Mini) was added and cells were lysed on ice for 20 minutes. Cell lysates were centrifuged at 14,000 g for 20 minutes and the quantity of protein in the supernatants was quantified using Bradford assays (Bio-Rad #5000002). Laemmli sample buffer (Bio-Rad #1610747, 5% 2-mercaptoethanol) was added to 20 µg protein and samples were boiled at 95°C for 5 minutes before being loaded onto a 10% SDS-polyacrylamide gel (Bio-Rad #4568034). Protein was transferred to polyvinylidene fluoride (PVDF) membranes (Thermo Scientific #88520), which were incubated overnight with primary antibody. Membranes were washed 3 times with TBS-T and incubated with horseradish peroxidase-conjugated anti-rabbit or anti-mouse for 1 hour at room temperature, washed, and imaged using an Amersham Imager 600.

### ATAC-seq

ATAC-seq was performed according to [105, 106] with minor alterations. Briefly, 70,000 *TFAP2*-KO cells (clone 2.12 and clone 4.3, four replicates each) and WT cells (four replicates) were lysed in ice-cold lysis buffer (10 mM Tris-HCl, pH 7.4, 10 mM NaCl, 3 mM MgCl2, 0.1% NP-40: Sigma). Transposition was performed directly on nuclei using 25 µl tagmentation reaction mix (Tagment DNA Buffer #15027866, Tagment DNA Enzyme #15027865 from Illumina Tagment DNA kit #20034210). Tagged DNA was subjected to PCR amplification and library indexing, using the NEBNext High-Fidelity 2x PCR Master Mix (New England Biolabs #M0451S) with Nextera DNA CD Indexes (Illumina #20015882), according to the following program: 72 C for 5 minutes; 98 C for 30 seconds; 12 cycles of 98 C for 10 seconds, 63 C for 1 minute. The PCR product was purified with 1.8 C for 30 seconds, and 72 times the volume of Ampure XP beads (Beckman Coulter #A63881). Library quality was assessed using a BioAnalyzer 2100 High Sensitivity DNA Chip (Agilent Technologies). All DNA libraries that exhibited a nucleosome pattern were pooled and processed for 150bp paired-end sequencing.

### ATAC-seq peak calling and differential analysis

ATAC-seq was performed using 150 bp paired-end sequencing reads. Raw ATAC- seq reads were trimmed using Trim Galore Version 0.6.3 (Developed by Felix Krueger at the Babraham Institute) and aligned to human genome assembly hg19 (GRCh37) using Bowtie 2 [107, 108] with default parameters. Sorting, removal of PCR duplicates, and identification of fragments shorter than 100 bp as the nucleosome-depleted-regions (NDRs), was performed using BAM filter version 0.5.9. DeepTools version 3.3.0 [109] was used to check the reproducibility of the biological replicates and generate bigWig coverage files for visualization. Peaks were called using model-based analysis of ChIP-seq 2 (MACS2, version 2.1.1.20160309.6) [110]. NDRs for which accessibility differed between *TFAP2*-KO and WT cells were identified using DiffBind version 2.10 [111] with log2 fold-change threshold of >1 and a false discovery rate (FDR) < 0.05. NDRs that are directly regulated by TFAP2 were identified by overlapping differentially accessible NDRs with anti-TFAP2A CUT&RUN peaks; a 1-bp window was used to define overlap. Peaks were assigned to genes using GREAT with a peak-to-gene association rule of the nearest-gene-within-100 kb [112]. Both the raw ATAC-seq files and processed sequencing data presented in this manuscript have been deposited in the Gene Expression Omnibus (GEO) repository (GSE number pending).

### CUT&RUN

CUT&RUN sequencing was performed in *TFAP2*-KO cells (clone 2.12 and clone 4.3, two replicates each) and WT cells (two replicates) as previously described [113, 114], but with minor modifications. Cells in log-phase culture (approximately 80% confluent) were harvested by cell scraping, centrifuged at 600 g (Eppendorf, centrifuge 5424) and washed twice in calcium-free wash-buffer (20 mM HEPES, pH7.5, 150 mM NaCl, 0.5 mM spermidine, and protease inhibitor cocktail cOmplete Mini, EDTA-free from Roche). Pre-activated concanavalin A-coated magnetic beads (Bangs Laboratories, Inc) were added to cell suspensions (2×10^5^ cells) and incubated for 15 minutes at 4°C. Antibody buffer (wash-buffer with 2mM EDTA and 0.05% digitonin) containing anti-TFAP2A, anti-MITF, anti-H3K4Me3, anti-H3K27Me3, anti-H3K27Ac or Rabbit IgG was added and cells were incubated overnight at 4°C. Cells were washed in dig-wash buffer (wash buffer containing 0.03% digitonin), and pAG-MNase was added at a concentration of 500 μg/mL. The pAG-MNase reactions were quenched with 2X Stop Buffer (340mM NaCl, 20mM EDTA, 4mM EGTA, 0.05% digitonin, 100 μg/mL RNAse A (10 mg/mL, Thermo Fisher Scientific #EN0531), 50 μg/mL glycogen (20mg/mL, Thermo Fisher Scientific #R0561) and 2 pg/mL sonicated yeast spike-in control). Released DNA fragments were treated with 1 μL/mL phosphatase K (20mg/mL, Thermo Fisher Scientific #25530049) for 1 hour at 50°C and purified by phenol/chloroform-extraction and ethanol-precipitation. Fragment sizes were analyzed using a 2100 Bioanalyzer (Agilent).

### CUT&RUN library preparation and data analysis

CUT&RUN libraries were prepared using the KAPA Hyper Prep Kit (Roche). Quality control post-library amplification consisted of fragment analysis using the 2100 Bioanalyzer (Agilent). Libraries were pooled, brought to equimolar concentrations, and sequenced with 150 bp paired-end reads on an Illumina HiSeq X platform (Novogene, Sacramento, CA). For quality control, paired-end FASTQ files were processed using FastQC (Babraham Bioinformatics). Reads were trimmed using Trim Galore Version 0.6.3 (Developed by Felix Krueger at the Babraham Institute) and then mapped against the hg19 genome assembly using Bowtie2 version 2.1.0 [107, 108]. The mapping parameters and peak calling of MACS2 peaks [110] were performed as previously described [113, 114] against their matching control IgG samples. Differential analysis of H3K27Ac and of H3K27Me3 signal in WT and *TFAP2*-KO cells was preformed using MACS2 with a Log2 fold-change threshold of 1, and p-value < 1 x 10^-5^. Differential H3K4Me3, MITF and TFAP2A signal in WT, *TFAP2*-KO and when mentioned *MITF*-KO cells was determined using DiffBind version 2.10.0 [111] with a Log2 fold-change threshold of 1, and FDR < 0.05. DeepTools version 3.3.0 was used to check the reproducibility of the biological replicates, to generate bigwig normalized (RPKM) coverage files for visualization and to plot average CUT&RUN-seq and ATAC-seq profiles (-plotProfile) and generate heatmaps (-plotHeatmap) of normalized reads [109]. MultiBigwigSummary was used to extract read counts (-outRawCounts) [109] and Prism was used to generate Violin and Box plots. Peaks were assigned to genes using GREAT with a peak-to-gene association rule of the nearest-gene-within-100 kb [112]

### RNA-seq

Four replicate RNA-seq experiments were performed on *TFAP2*-KO cells (clone 2.12 and clone 4.3) and WT cells. Total RNA was extracted by direct cell lysis using the RNeasy Plus Mini Kit with QiaShredder (Qiagen #47134). RNA samples with an RNA integrity number (RIN) above nine were used for library generation and 150 bp paired-end sequencing on the Illumina HiSeq2500 platform (Novogene, Sacramento, CA). FASTQ sequence files were processed using FastQC (Babraham Bioinformatics) for quality control, and reads were trimmed using Trim Galore Version 0.6.3 (Developed by Felix Krueger at the Babraham Institute) and subsequently aligned to human genome assembly hg19 (GRCh37) using STAR [115]. The output of the --quantMode GeneCounts function of STAR was used for the calculation of differential transcript expression using DESeq2 [116].The rlog function was used to generate log2-transformed normalized counts. Adjusted p-value < 0.05 was used as the threshold for statistically significant differences. Functional enrichment analyses was performed using PANTHER [117]. A full list of genes differentially expressed between *TFAP2*-KO and WT cells is provided in Supplemental Table 1-3.

### Motif analyses

Both de novo and known motifs were identified within 200 bp of TFAP2-activated and TFAP2-inhibited enhancer and promoter peak summits using HOMER (- findMotifsGenome).

### Statistical analysis

Fisher’s Exact Test was used to assess association between TFAP2-regulated elements (enhancers and promoters) with TFAP2-regulated gene expression. Fisher’s Exact Test is a statistical significance test based on the hypergeometric distribution and is used to analyze contingency tables to determine if there is an association between the classes in the columns and rows of the contingency table. Use of this test is relevant for the analysis of differential expression because we classify enhancers as either TFAP2-activated (downregulated in TFAP2-KO cells) or TFAP2-inhbited (upregulated in TFAP2-KO cells) as well as near a target of interest (i.e., TFAP2-activated or TFAP2-inhibited) or not. This test determines if there is an association between genes that are near an TFAP2-regulated enhancer and genes whose expression varies between TFAP2-KO cells and WT cells.

Data processing and analysis was performed in R and the code can be found at https://GitHub.com/ahelv/Differential_Expression. GraphPad-Prism was used to perform Student’s t-test as indicated in the figure legends.

### Proliferation assay

Cell proliferation was monitored at 24, 48, 72 and 96 hours using the CellTiter 96 Aqueous One Solution reagent kit according to the manufacturer’s protocol. Cells were serum depleted for 24 hours before transferring 5,000 cells per well of cell culture treated 96-well plate (corning) containing RPMI media supplemented with 10% FBS and 1% PS. One the day of the experiment, CellTiter 96 Aqueous One Solution was added to cells at a ratio of 6:1 (Media: solution). After 1 hour at 37°C in a humidified, 5% CO_2_ atmosphere, the absorbance at 450nm was recorded using an ELISA plate reader. Background absorbance was measured from cell free wells containing media and CellTiter 96 Aqueous One Solution. The experiment was performed in triplicate and repeated 3 times.

### Cluster formation assay

Cluster formation assays were performed as previously described [32]. In brief, cells were trypsinized and resuspended in RPMI media supplemented with 10% FBS and 1% PS. A flat bottom Low Attachment Surface-well plate (Corning) was seeded with 5,000 cells/well in a final volume of 100uL. Clusters were allowed to form over the course of 3 days in a humidified 37°C incubator at which time the number of clusters were counted per well using Image J software. The experiment was performed in triplicate and repeated twelve times.

### Wound scratch assay

A total of 500K cells were seeded per well of 6-well plate (Thermo Scientific, # 1483210) to reach confluent monolayer. Cells were incubated in serum free media for 6 hours before wounding with a 200 µL pipette tip. Scratches were manually imaged on an inverted light microscope (Leica #10445930) every 6 hours over a 24- hour time period. The distance of scratch closure between WT and *TFAP2*-KO cells were analyzed with Image J software.

### Data availability

All raw sequencing files and processed data presented in this manuscript have been deposited in the Gene Expression Omnibus (GEO) repositor. For ATAC-Seq and CUT&RUN-seq data (GSE190610 and GSE153020) and for single cell RNA-Seq (GSE198791).

## Acknowledgements

This work was supported by grants from the National Institutes of Health (NIH) to RAC (R01-AR062457), and a postdoctoral fellowship from the American Association for Anatomy (CK). The ES laboratory is supported by grants 207067 and 217768 from the Research Fund of Iceland.

## Author Contributions

Conceptualization: CK, RAC.

Formal analysis: CK, RD.

Funding acquisition: CK, ES, RAC.

Investigation: CK, RAC.

Methodology: CK, RD, HS, EVO, AH, GB.

Statistical analysis: CK, CMF, AH.

Supervision: ES, RAC.

Writing – original draft: CK, RAC.

## Figure legends

**Figure S1: UMAPs showing expression of select genes in *foxd3+* neural crest, *twist+* cranial neural crest and basal cell clusters.** (**A-C**) Uniform Manifold Approximation and Projection (UMAP) obtained after clustering (dimensions, dims = 30, resolution = 1.2) GFP+ cells (n = 11,217 cells) sorted from *Tg(mitfa:GFP)* zebrafish embryos at 28 hours post fertilization (hpf). Feature plots showing expression of select marker genes (**A**) *foxd3*+ neural crest clusters (**B**) *twist*+ cranial neural crest clusters and (**C**) basal cell clusters. (**D**) Feature plots showing expression of *mitfa* and *sox10*. Note that *mitfa* positive clusters are a subset of all *sox10* positive clusters. Dotted lines indicate clusters of interest.

**Figure S2: Expression of select marker genes in *foxd3/her*+ neural crest, MIX, xanthophore and iridophore cell clusters.** (**A**) Violin plots showing expression of Notch pathway genes in *foxd3/her*+ NC cluster 6. Note the lack of notch signaling gene expression in *foxd3+* NC cluster 7. (**B**) Violin plots showing expression of select genes in MIX cell clusters, some of which are also expressed in MX, xanthoblast and iridoblast cell clusters. (**C**) Violin plots showing expression of select marker genes in xanthoblast and xanthophore cell clusters. (**D**) Violin plots showing expression of select marker genes in iridoblast and iridophore cell clusters.

**Figure S3: Expression of TFAP2 paralogs in the pigment cell lineage.** (**A**) Violin plots showing expression of *tfap2* paralogs in NC, MIX *tfap2*-low, MIX *tfap2-*high, MX, melanophore, xanthoblast, xanthophore, iridoblast and iridophore cell clusters. Numeric labels below each violin plot refer cluster numbers as described in Figure 1B.

**Figure S4: Expression of *mitfa* and *tfap2e* precede activation of Mitfa target gene in the pigment cell lineage.** (**A**) UMAP feature plots and **B**) violin plots showing expression of *mitfa, tfap2e,* and genes in the melanin synthesis pathway, whose human orthologs are known to be directly activated by MITF, in the pigment cell lineage. Although robust *mitfa* expression is present in the MIX-*tfap2*-low cluster (cluster 8a), expression of genes in the melanin synthesis pathway is absent in this cluster. Expression of the latter begins concurrent with expression of *tfap2e* in the MIX-*tfap2*-high cluster (cluster 8b).

**Figure S5: Zebrafish homozyg**ous for a frame-shift inducing mutation in *tfap2e* do not display a pigmentation phenotype, and melanophores in *tfap2a/tfap2e* double mutants are melanized by 48 hpf. (A) A 157 base pair mutation at the end of *tfap2e* exon 2 disrupts splicing and results in a premature stop codon. (B) PCR using primers in *tfap2e* exon 2 and intron 2 (e2-i2) amplifies a band of the expected 450 base pair size in *tfap2e* mutants but not wildtype (WT), whereas primers in exon 2 and exon 3 (e2-e3) amplify only in wild type. NTC: not template control. (C) qRT-PCR analysis of *tfap2e* expression shows that the transcript is strongly decreased in *tfap2e ^ui157/ ui157^* mutants, consistent with nonsense-mediated decay (Student’s t-test, **** p<0.0001). (D-E) *tfap2e* mutant zebrafish at 36 hpf, *tfap2e*^+/ *ui157*^ (D) and *tfap2e ^ui157/ ui157^* (E) are phenotypically indistinguishable. (F) Histogram illustrating the number of pigmented melanocytes in the dorsum of *tfap2e*^+/ *ui157*^ and *tfap2e ^ui157/ ui157^* mutant zebrafish embryos. G-J) Lateral views of 48 hpf embryos in the chorion. G) A wildtype embryo shows normal melanocyte patterning. (H) A *tfap2a^-/-^* mutant embryo, with fewer and paler embryonic melanocytes than wildtypes. (I) In *tfap2a^-/-^ ;tfap2e^+/-^* and (J) in *tfap2a^-/-^;tfap2e^-/-^* mutants individual melanocytes are grossly as melanized as they are in wild-types. These mutants are phenotypically indistinguishable from *tfap2a^-/-^* mutant embryos.

**Figure S6: qRT-PCR expression analysis of *tfap2* paralogs in *tfap2* zebrafish mutants.** (**A**) Expression of *tfap2* paralogs in pooled embryos. Wildtype (n=20), *tfap2a^low/low^* (n=20), and *tfap2a^low/low^*; *tfap2e ^ui157/ ui157^* (n=18). One-way ANOVA with Bonferroni correction, * signifies adj. p<0.05. RQ, relative quantification.

**Figure S7: TFAP2 binds to open and closed chromatin.** (**A**) Screenshot of IGV genome browser (GRCH37/hg19), visualizing anti-TFAP2A and IgG CUT&RUN-seq profiles. Peaks were called using MACS2 software (two independent replicates) and are illustrated by blue bars under the anti-TFAP2A track. (**B**) Density heatmap centred on the 36,867 TFAP2A peaks identified by anti-TFAP2A CUT&RUN in WT SK-MEL-28 cells. Regions shown are +/- 3 kb from peak center. Promoter peaks are < 3kb and enhancer peaks >3kb from a transcriptional start site (TSS). Anti-TFAP2A, anti-H3K4Me3 and anti-H3K27Ac CUT&RUN-seq, and ATAC-Seq profiles are shown. Color code reflects normalized Reads Per Kilobase, per Million mapped reads (RPKM). (**C**) Histogram representing H3K27Ac signal, binned from low to high read-depth (normalized RPKM) on the x-axis and percentage of TFAP2A promoter peaks (black) and TFAP2A enhancer peaks (red) on the y-axis. (**D**) Violin plots illustrating TFAP2A and IgG normalized reads (RPKM) at open (ATAC-peaks) and at closed chromatin (no ATAC-peak) (**E**) Density heatmap representing TFAP2A CUT&RUN and ATAC-seq profiles at TFAP2A peaks that overlap nucleosome depleted regions (ATAC-peaks) and at nucleosome bound DNA (no ATAC-peak), the number of TFAP2A peaks in each group are as labeled.

**Figure S8: Example loci of TFAP2A peaks at open and closed chromatin.** (**A-B**) IGV screenshots of TFAP2A peaks that (**A**) closed chromatin and (**B**) open chromatin, based on ATAC-seq, in SK-MEL-28 cells. Genes names and distance to that gene’s TSS are as labeled. Note that scale for ATAC-seq reads (RPKM) is constant but for TFAP2A CUT&RUN is variable. *Schematics*, renderings of TFAP2 in **A**) closed and **B**) open chromatin. (**C**) HOMER motif analysis at TFAP2A peaks at closed chromatin and (**D**) at open chromatin. TF, transcription factors; the top ranking transcription factor motif is shown, with P-values calculated with HOMER-based hypergeometric enrichment analysis.

**Figure S9: Generation of *TFAP2A*; *TFAP2C* double mutant SK-MEL-28 cell lines.** (**A**) RNA-seq showing transcript counts of TFAP2 paralogs in SK-MEL-28 cells. Transcript counts for WT cells (n=4) and two *TFAP2A;TFAP2C* double knockout clones (4 replicates each) are shown. The expression of WT and mutant alleles of *TFAP2A* is comparable between cell lines whereas mutant alleles of *TFAP2C* are strongly reduced. (**B**) Two CRISPR guide RNAs (gRNA) each were designed to target exon 2 of *TFAP2A* and *TFAP2C* (yellow boxes). (**C**) A 401 base pair inversion and a 452 base pair deletion at exon 2 of *TFAP2A* and *TFAP2C*, respectively, was identified in clone 4.3. *TFAP2A* and *TFAP2C* mutant alleles resulted in a frame-shift and premature stop codon in alleles of both genes. (**D**) A 405 base pair deletion at exon 2 of *TFAP2A* resulted in a frame-shift and premature stop codon in clone 2.12. A 455 base pair deletion, and a 70 base pair insertion, 1 base pair deletion (Indel) was identified in exon 2 of *TFAP2C*. Such mutations resulted in a frame-shift and premature stop codon. Additional permutations were not identified at exon 2 of *TFAP2A* or *TFAP2C* in clone 4.3 or clone 2.12 cells. Inv, inversion; Del, deletion. (**E**) Western blot analysis confirming loss of TFAP2A and TFAP2C immunoactivity in clone 4.3 and clone 2.12 cell lines.

**Figure S10: TFAP2 paralogs directly activate and inhibit promoters as pioneer factors.** (**A-D**) Screenshot of IGV genome browser (GRCH37/hg19), visualizing anti- TFAP2A CUT&RUN-seq (red), ATAC-seq (black), anti-H3K4Me3 CUT&RUN-seq (blue), anti-H3K27Ac CUT&RUN-seq (green) and RNA-seq (magenta) datasets at (**A**) TFAP2-pioneered-and-activated promoters, (**B**) non-pioneered TFAP2-activated promoters, (**C**) TFAP2-pioneered-and-inhibited promoters and (**D**) non-pioneered TFAP2-activated promoters . Genotypes as labeled; y-axes are grouped scaled per dataset. (**E-H**) Violin plots representing anti-H3K27Ac (two independent replicates) (**E’-H’**) ATAC-seq (four independent replicates), and (**E’’-H”**) anti-H3K4Me3 (two independent replicates) normalized reads at (**E-E’’**) TFAP2-pioneered-and-activated promoters, (**F’F”**) non-pioneered TFAP2-activated promoters, (**G-G’’**) TFAP2- pioneered-and-inhibited promoters, (**H-H’’**) non-pioneered TFAP2-inhibited promoters. P-values shown were determined by Student’s t-test. (**I-L**) Hypergeometric analysis of TFAP2 regulated enhancers at TFAP2-activated (*) and TFAP2-inhibited (**) genes in *TFAP2*-KO cells (FDR < 0.05, |log2FC| > 1). The number of promoters in each category of TFAP2-regulated promoters is shown.

**Figure S11: Examples of TFAP2-activated and -inhibited promoters.** (**A-B**) Screenshots of IGV genome browser (GRCH37/hg19) visualizing anti-TFAP2A, anti- H3K4Me3, anti-H3K27Ac CUT&RUN-seq, ATAC-seq and RNA-seq profiles at (**A**) the TFAP2-pioneered-and-activated *ZNF540* promoter and (**B**) the TFAP2- pioneered-and-inhibited *S100A16* promoter. Genotypes as labeled; y-axes are scaled per dataset.

**Figure S12: TFAP2 paralogs modulate the binding of MITF.** (**A**) Density heatmap centred on 9,413 peaks co-bound by TFAP2A and MITF, showing anti-TFAP2, anti-MITF, ATAC-seq, and anti-H3K27Ac CUT&RUN profiles in SK-MEL-28 cells. Regions shown are +/- 5 kb from peak center. (**B**) Violin plot showing anti-MITF CUT&RUN signal (RPKM) in *TFAP2*-KO and SK-MEL-28 cells at MITF peaks that are not bound by TFAP2A. There is no change in the average read-depth of such MITF peaks in WT vs TFAP2-KO cells. ns; non-significant by Student’s t-test. **(C)** Scatterplot of TFAP2-dependent (i.e., TFAP2-activated and TFAP2-inhibited) MITF peaks. X axis is log2 normalized MITF CUT&RUN read depth in peaks in WT cells. Y axis is log2FC of MITF CUT&RUN read depth in *TFAP2*-KO versus WT cells. **(D)** Screenshots of IGV genome browser (GRCH37/hg19); genotypes as labeled, visualizing 5kb windows centered on TFAP2A peaks and showing indicated genomic data at several examples of TFAP2-activated MITF peaks and mutually-dependent MITF/TFAP2 peaks, and a single example of a non-overlapping, TFAP2-independent MITF peak. **(E)** Violin plot showing increased H3K27Me3 CUT&RUN signal at mutually-dependent TFAP2A/MITF peaks in *MITF*-KO cells versus WT cells. P-values were determined by Student’s t-test. Y axis is normalized reads (RPKM). **(F-G)** Screenshots of IGV genome browser (GRCH37/hg19), visualizing anti- H3K27me3 CUT&RUN-seq profiles in *MIT*F-KO and WT cells. Peaks were called using MACS2 software (two independent replicates) and are illustrated by blue bars. Two examples loci of mutually-dependent TFAP2A/MITF peaks showing increased H3K27Me3 signals are shown **(F)** at the *TRPM1* promoter and **(G)** at two *FRMD4B* enhancers. Note that H3K27Me3 signal extends much farther than just the region of the TFAP2A/MITF peak. Mutually-dependent peaks are indicated by red arrows.

**Figure S13: Density heatmap of TFAP2 regulated MITF peaks.** Density heatmap centered on TFAP2-activated MITF peaks (first cluster), mutually dependent (mutually activated) peaks (second cluster), TFAP2-inhibited MITF peaks and (third cluster) and TFAP2-independent MITF peaks (forth cluster), showing two replicates of anti-MITF CUT&RUN in SK-MEL-28 and *TFAP2*-KO cells, and two replicates of anti-TFAP2A CUT&RUN in SK-MEL-28 and *MITF*-KO cells. Regions shown are +/- 5 kb from peak center.

**Figure S14: TFAP2 and MITF do not co-inhibit enhancers at TFAP2-inhibited or MITF-inhibited genes in SK-MEL-28 cells.** (**A-A’**) Violin plot of TFAP2-activated MITF peaks at TFAP2-pioneered-and-inhibited enhancers, which are potentially TFAP2/MITF co-inhibited enhancers, showing (**A**) anti-MITF CUT&RUN and (**A’**) ATAC-Seq profiles in *TFAP2*-KO and WT cells. Such loci were not significantly enriched at MITF-inhibited or TFAP2-inhibited genes. (**A’’**) Schematic of theoretical TFAP2/ MITF co-inhibited enhancers. In this example, TFAP2 is a pioneer factor recruiting MITF in its repressor form to condense chromatin. Loss of TFAP2 in *TFAP2*-KO cells results in loss of MITF-repressor binding and opening of chromatin by an alternative pioneer factor.

**Figure S15: MITF-activated genes associated with TFAP2-independent MITF peaks are enriched for cell cycle and DNA-repair.** (**A**) Plot-profile showing MITF CUT&RUN peak signal at TFAP2-independent MITF peaks in *TFAP2*-KO and WT SK-MEL-28 cell lines. (**B**) Genes associated with TFAP2-independent anti-MITF peaks were analyzed for enriched gene ontology biological process using GREAT (single nearest gene +/- 100kb).

**Figure S16: A possible TFAP2-pioneered-and-activated enhancer at intron 2 of MITF.** Screenshot of IGV genome browser (GRCH37/hg19) visualizing anti-TFAP2A, ATAC-seq, anti-H3K27Ac and RNA-seq profiles at intron 2 of MITF. Dashed rectangle indicates an TFAP2-activated ATAC-seq peak; like the ATAC-seq signal, the H3K27Ac-signal is significantly lower in TFAP2-KO cells relative to in WT cells implying this is a TFAP2-pioneered-and-activated enhancer, although the log2 fold change of the H3K27Ac signal does not meet the cut-off we used to define this category of enhancer elsewhere in the paper. MITF and downstream regions are shown, blue arrows indicate strand orientation and horizontal rectangles the exons. Genotypes are as labeled; y-axes are grouped scaled per dataset.

**Figure S17: GO analysis of TFAP2-activated and inhibited genes.** (**A**) Heatmap representing the top 55 directly TFAP2-activated genes that are included in the Gene Ontology (GO) biological process categories “cell pigmentation” and “cell differentiation.” Four replicate RNA-Seq experiments are shown for WT cells and two clones of *TFAP2*-KO cells (Clone 2.12 and Clone 4.3) (FDR < 0.05). (**B-C**) GO biological process terms enriched among (**B**) directly TFAP2-activated genes and (**C**) directly TFAP2-inhibited genes.

**Figure S18: Scratch assay.** Micrographs of (top) WT and (bottom) *TFAP2*-KO cells in petri dishes at the indicated time after a scratch through the cells was made with a pipette tip. After 24 hours, the WT but not the *TFAP2*-KO cells had filled in the scratch through migration and/or proliferation.

## References cited

1. Seberg HE, Van Otterloo E, Cornell RA. Beyond MITF: Multiple transcription factors directly regulate the cellular phenotype in melanocytes and melanoma. Pigment Cell Melanoma Res. 2017;30(5):454–66. doi: 10.1111/pcmr.12611. PubMed PMID: 28649789.

2. Goding CR. Mitf from neural crest to melanoma: signal transduction and transcription in the melanocyte lineage. Genes Dev. 2000;14(14):1712–28. Epub 2000/07/18. PubMed PMID: 10898786.

3. Hartman ML, Czyz M. MITF in melanoma: mechanisms behind its expression and activity. Cell Mol Life Sci. 2015;72(7):1249–60. doi: 10.1007/s00018-014-1791-0. PubMed PMID: 25433395; PubMed Central PMCID: PMCPMC4363485.

4. Mollaaghababa R, Pavan WJ. The importance of having your SOX on: role of SOX10 in the development of neural crest-derived melanocytes and glia. Oncogene. 2003;22(20):3024–34. PubMed PMID: 12789277.

5. Atchison ML. Function of YY1 in Long-Distance DNA Interactions. Front Immunol. 2014;5:45. Epub 2014/02/28. doi: 10.3389/fimmu.2014.00045. PubMed PMID: 24575094; PubMed Central PMCID: PMCPMC3918653.

6. Eckert D, Buhl S, Weber S, Jager R, Schorle H. The AP-2 family of transcription factors. Genome biology. 2005;6(13):246. doi: 10.1186/gb-2005-6-13-246. PubMed PMID: 16420676; PubMed Central PMCID: PMC1414101.

7. Hoek KS, Schlegel NC, Eichhoff OM, Widmer DS, Praetorius C, Einarsson SO, et al. Novel MITF targets identified using a two-step DNA microarray strategy. Pigment Cell Melanoma Res. 2008;21(6):665–76. Epub 2008/12/11. doi: 10.1111/j.1755-148X.2008.00505.x. PubMed PMID: 19067971.

8. Strub T, Giuliano S, Ye T, Bonet C, Keime C, Kobi D, et al. Essential role of microphthalmia transcription factor for DNA replication, mitosis and genomic stability in melanoma. Oncogene. 2011;30(20):2319–32. Epub 2011/01/25. doi: onc2010612 [pii] 10.1038/onc.2010.612. PubMed PMID: 21258399.

9. Betancur P, Bronner-Fraser M, Sauka-Spengler T. Assembling neural crest regulatory circuits into a gene regulatory network. Annu Rev Cell Dev Biol. 2010;26:581–603. Epub 2009/07/07. doi: 10.1146/annurev.cellbio.042308.113245. PubMed PMID: 19575671.

10. Van Otterloo E, Li W, Bonde G, Day KM, Hsu MY, Cornell RA. Differentiation of zebrafish melanophores depends on transcription factors AP2 alpha and AP2 epsilon. PLoS genetics. 2010;6(9). Epub 2010/09/24. doi: 10.1371/journal.pgen.1001122. PubMed PMID: 20862309; PubMed Central PMCID: PMC2940735.

11. Van Otterloo E, Li W, Garnett A, Cattell M, Medeiros DM, Cornell RA. Novel Tfap2-mediated control of soxE expression facilitated the evolutionary emergence of the neural crest. Development. 2012;139(4):720–30. Epub 2012/01/14. doi: 10.1242/dev.071308. PubMed PMID: 22241841; PubMed Central PMCID: PMC3265060.

12. Carreira S, Goodall J, Denat L, Rodriguez M, Nuciforo P, Hoek KS, et al. Mitf regulation of Dia1 controls melanoma proliferation and invasiveness. Genes Dev. 2006;20(24):3426–39. Epub 2006/12/22. doi: 10.1101/gad.406406. PubMed PMID: 17182868; PubMed Central PMCID: PMC1698449.

13. Rambow F, Marine JC, Goding CR. Melanoma plasticity and phenotypic diversity: therapeutic barriers and opportunities. Genes Dev. 2019;33(19-20):1295–318. Epub 2019/10/03. doi: 10.1101/gad.329771.119. PubMed PMID: 31575676; PubMed Central PMCID: PMCPMC6771388.

14. de la Serna IL, Ohkawa Y, Higashi C, Dutta C, Osias J, Kommajosyula N, et al. The microphthalmia-associated transcription factor requires SWI/SNF enzymes to activate melanocyte-specific genes. J Biol Chem. 2006;281(29):20233–41. Epub 2006/05/02. doi: M512052200 [pii] 10.1074/jbc.M512052200. PubMed PMID: 16648630.

15. Laurette P, Strub T, Koludrovic D, Keime C, Le Gras S, Seberg H, et al. Transcription factor MITF and remodeller BRG1 define chromatin organisation at regulatory elements in melanoma cells. eLife. 2015;4. doi: 10.7554/eLife.06857. PubMed PMID: 25803486; PubMed Central PMCID: PMC4407272.

16. Lang D, Lu MM, Huang L, Engleka KA, Zhang M, Chu EY, et al. Pax3 functions at a nodal point in melanocyte stem cell differentiation. Nature. 2005;433(7028):884–7. Epub 2005/02/25. doi: nature03292 [pii] 10.1038/nature03292. PubMed PMID: 15729346.

17. Dilshat R, Fock V, Kenny C, Gerritsen I, Lasseur RMJ, Travnickova J, et al. MITF reprograms the extracellular matrix and focal adhesion in melanoma. eLife. 2021;10. Epub 2021/01/14. doi: 10.7554/eLife.63093. PubMed PMID: 33438577; PubMed Central PMCID: PMCPMC7857731.

18. Du J, Li L, Ou Z, Kong C, Zhang Y, Dong Z, et al. FOXC1, a target of polycomb, inhibits metastasis of breast cancer cells. Breast Cancer Res Treat. 2012;131(1):65–73. Epub 2011/04/06. doi: 10.1007/s10549-011-1396-3. PubMed PMID: 21465172.

19. Mitchell PJ, Timmons PM, Hebert JM, Rigby PW, Tjian R. Transcription factor AP-2 is expressed in neural crest cell lineages during mouse embryogenesis. Genes Dev. 1991;5(1):105–19.

20. Moser M, Ruschoff J, Buettner R. Comparative analysis of AP-2 alpha and AP-2 beta gene expression during murine embryogenesis. Developmental Dynamics. 1997;208(1):115–24.

21. Bamforth SD, Braganca J, Eloranta JJ, Murdoch JN, Marques FI, Kranc KR, et al. Cardiac malformations, adrenal agenesis, neural crest defects and exencephaly in mice lacking Cited2, a new Tfap2 co-activator. Nat Genet. 2001;29(4):469–74. PubMed PMID: 11694877.

22. Brewer S, Jiang X, Donaldson S, Williams T, Sucov HM. Requirement for AP-2alpha in cardiac outflow tract morphogenesis. Mech Dev. 2002;110(1-2):139–49. PubMed PMID: 11744375.

23. Knight RD, Nair S, Nelson SS, Afshar A, Javidan Y, Geisler R, et al. lockjaw encodes a zebrafish tfap2a required for early neural crest development. Development. 2003;130(23):5755–68. PubMed PMID: 14534133.

24. Tan CC, Sindhu KV, Li S, Nishio H, Stoller JZ, Oishi K, et al. Transcription factor Ap2delta associates with Ash2l and ALR, a trithorax family histone methyltransferase, to activate Hoxc8 transcription. Proc Natl Acad Sci U S A. 2008;105(21):7472–7. Epub 2008/05/23. doi: 10.1073/pnas.0711896105. PubMed PMID: 18495928; PubMed Central PMCID: PMCPMC2396708.

25. Schorle H, Meier P, Buchert M, Jaenisch R, Mitchell PJ. Transcription factor AP-2 essential for cranial closure and craniofacial development. Nature. 1996;381(6579):235–8.

26. Luo T, Matsuo-Takasaki M, Thomas ML, Weeks DL, Sargent TD. Transcription factor AP-2 is an essential and direct regulator of epidermal development in Xenopus. Dev Biol. 2002;245(1):136–44. PubMed PMID: 11969261.

27. Wang X, Bolotin D, Chu DH, Polak L, Williams T, Fuchs E. AP-2alpha: a regulator of EGF receptor signaling and proliferation in skin epidermis. J Cell Biol. 2006;172(3):409–21. Epub 2006/02/02. doi: jcb.200510002 [pii] 10.1083/jcb.200510002. PubMed PMID: 16449191; PubMed Central PMCID: PMC2063650.

28. Kuckenberg P, Kubaczka C, Schorle H. The role of transcription factor Tcfap2c/TFAP2C in trophectoderm development. Reproductive biomedicine online. 2012;25(1):12–20. doi: 10.1016/j.rbmo.2012.02.015. PubMed PMID: 22560121.

29. Li W, Cornell RA. Redundant activities of Tfap2a and Tfap2c are required for neural crest induction and development of other non-neural ectoderm derivatives in zebrafish embryos. Dev Biol. 2007;304(1):338–54. Epub 2007/01/30. doi: S0012-1606(06)01501-6 [pii] 10.1016/j.ydbio.2006.12.042. PubMed PMID: 17258188; PubMed Central PMCID: PMC1904501.

30. Wang X, Pasolli HA, Williams T, Fuchs E. AP-2 factors act in concert with Notch to orchestrate terminal differentiation in skin epidermis. J Cell Biol. 2008;183(1):37–48. Epub 2008/10/01. doi: jcb.200804030 [pii] 10.1083/jcb.200804030. PubMed PMID: 18824566; PubMed Central PMCID: PMC2557040.

31. Kołat D, Kałuzińska Ż, Bednarek AK, Płuciennik E. WWOX Loses the Ability to Regulate Oncogenic AP-2γ and Synergizes with Tumor Suppressor AP-2α in High-Grade Bladder Cancer. Cancers (Basel). 2021;13(12). Epub 2021/07/03. doi: 10.3390/cancers13122957. PubMed PMID: 34204827; PubMed Central PMCID: PMCPMC8231628.

32. Campbell NR, Rao A, Zhang M, Baron M, Heilmann S, Deforet M, et al. Cell state diversity promotes metastasis through heterotypic cluster formation in melanoma. Developmental Cell. 2021:2020.08.24.265140. doi: 10.1101/2020.08.24.265140.

33. Seberg HE, Van Otterloo E, Loftus SK, Liu H, Bonde G, Sompallae R, et al. TFAP2 paralogs regulate melanocyte differentiation in parallel with MITF. PLoS genetics. 2017;13(3):e1006636. doi: 10.1371/journal.pgen.1006636. PubMed PMID: 28249010; PubMed Central PMCID: PMCPMC5352137.

34. Rothstein M, Simoes-Costa M. Heterodimerization of TFAP2 pioneer factors drives epigenomic remodeling during neural crest specification. Genome Res. 2020;30(1):35–48. Epub 20191217. doi: 10.1101/gr.249680.119. PubMed PMID: 31848212; PubMed Central PMCID: PMCPMC6961570.

35. Sennett R, Wang Z, Rezza A, Grisanti L, Roitershtein N, Sicchio C, et al. An Integrated Transcriptome Atlas of Embryonic Hair Follicle Progenitors, Their Niche, and the Developing Skin. Dev Cell. 2015;34(5):577–91. Epub 20150806. doi: 10.1016/j.devcel.2015.06.023. PubMed PMID: 26256211; PubMed Central PMCID: PMCPMC4573840.

36. Praetorius C, Grill C, Stacey SN, Metcalf AM, Gorkin DU, Robinson KC, et al. A polymorphism in IRF4 affects human pigmentation through a tyrosinase-dependent MITF/TFAP2A pathway. Cell. 2013;155(5):1022–33. doi: 10.1016/j.cell.2013.10.022. PubMed PMID: 24267888; PubMed Central PMCID: PMC3873608.

37. Rambow F, Job B, Petit V, Gesbert F, Delmas V, Seberg H, et al. New Functional Signatures for Understanding Melanoma Biology from Tumor Cell Lineage-Specific Analysis. Cell Rep. 2015;13(4):840–53. Epub 2015/10/23. doi: 10.1016/j.celrep.2015.09.037. PubMed PMID: 26489459; PubMed Central PMCID: PMCPMC5970542.

38. Voss TC, Hager GL. Dynamic regulation of transcriptional states by chromatin and transcription factors. Nat Rev Genet. 2014;15(2):69–81. Epub 2013/12/18. doi: 10.1038/nrg3623. PubMed PMID: 24342920; PubMed Central PMCID: PMCPMC6322398.

39. Zaret KS, Carroll JS. Pioneer transcription factors: establishing competence for gene expression. Genes Dev. 2011;25(21):2227–41. doi: 10.1101/gad.176826.111. PubMed PMID: 22056668; PubMed Central PMCID: PMCPMC3219227.

40. Zaret KS. Pioneer Transcription Factors Initiating Gene Network Changes. Annu Rev Genet. 2020;54:367–85. Epub 2020/09/05. doi: 10.1146/annurev-genet-030220-015007. PubMed PMID: 32886547; PubMed Central PMCID: PMCPMC7900943.

41. Sherwood RI, Hashimoto T, O’Donnell CW, Lewis S, Barkal AA, van Hoff JP, et al. Discovery of directional and nondirectional pioneer transcription factors by modeling DNase profile magnitude and shape. Nat Biotechnol. 2014;32(2):171–8. Epub 2014/01/21. doi: 10.1038/nbt.2798. PubMed PMID: 24441470; PubMed Central PMCID: PMCPMC3951735.

42. Pihlajamaa P, Sahu B, Lyly L, Aittomaki V, Hautaniemi S, Janne OA. Tissue-specific pioneer factors associate with androgen receptor cistromes and transcription programs. EMBO J. 2014;33(4):312–26. Epub 2014/01/24. doi: 10.1002/embj.201385895. PubMed PMID: 24451200; PubMed Central PMCID: PMCPMC3989639.

43. Tan SK, Lin ZH, Chang CW, Varang V, Chng KR, Pan YF, et al. AP-2gamma regulates oestrogen receptor-mediated long-range chromatin interaction and gene transcription. EMBO J. 2011;30(13):2569–81. doi: 10.1038/emboj.2011.151. PubMed PMID: 21572391; PubMed Central PMCID: PMC3155293.

44. Grossman SR, Engreitz J, Ray JP, Nguyen TH, Hacohen N, Lander ES. Positional specificity of different transcription factor classes within enhancers. Proc Natl Acad Sci U S A. 2018;115(30):E7222–E30. Epub 2018/07/11. doi: 10.1073/pnas.1804663115. PubMed PMID: 29987030; PubMed Central PMCID: PMCPMC6065035.

45. Pastor WA, Liu W, Chen D, Ho J, Kim R, Hunt TJ, et al. TFAP2C regulates transcription in human naive pluripotency by opening enhancers. Nat Cell Biol. 2018;20(5):553–64. Epub 2018/04/27. doi: 10.1038/s41556-018-0089-0. PubMed PMID: 29695788; PubMed Central PMCID: PMCPMC5926822.

46. Li L, Wang Y, Torkelson JL, Shankar G, Pattison JM, Zhen HH, et al. TFAP2C- and p63-Dependent Networks Sequentially Rearrange Chromatin Landscapes to Drive Human Epidermal Lineage Commitment. Cell Stem Cell. 2019;24(2):271–84 e8. Epub 2019/01/29. doi: 10.1016/j.stem.2018.12.012. PubMed PMID: 30686763; PubMed Central PMCID: PMCPMC7135956.

47. Fernandez Garcia M, Moore CD, Schulz KN, Alberto O, Donague G, Harrison MM, et al. Structural Features of Transcription Factors Associating with Nucleosome Binding. Mol Cell. 2019;75(5):921–32 e6. Epub 2019/07/16. doi: 10.1016/j.molcel.2019.06.009. PubMed PMID: 31303471; PubMed Central PMCID: PMCPMC6731145.

48. Swinstead EE, Paakinaho V, Presman DM, Hager GL. Pioneer factors and ATP- dependent chromatin remodeling factors interact dynamically: A new perspective: Multiple transcription factors can effect chromatin pioneer functions through dynamic interactions with ATP-dependent chromatin remodeling factors. Bioessays. 2016;38(11):1150–7. Epub 2016/10/27. doi: 10.1002/bies.201600137. PubMed PMID: 27633730; PubMed Central PMCID: PMCPMC6319265.

49. Aras S, Saladi SV, Basuroy T, Marathe HG, Lores P, de la Serna IL. BAF60A mediates interactions between the microphthalmia-associated transcription factor and the BRG1-containing SWI/SNF complex during melanocyte differentiation. J Cell Physiol. 2019;234(7):11780–91. Epub 2018/12/06. doi: 10.1002/jcp.27840. PubMed PMID: 30515787; PubMed Central PMCID: PMCPMC6426657.

50. Keenen B, Qi H, Saladi SV, Yeung M, de la Serna IL. Heterogeneous SWI/SNF chromatin remodeling complexes promote expression of microphthalmia-associated transcription factor target genes in melanoma. Oncogene. 2010;29(1):81–92. Epub 2009/09/29. doi: onc2009304 [pii] 10.1038/onc.2009.304. PubMed PMID: 19784067; PubMed Central PMCID: PMC2803337.

51. Butler A, Hoffman P, Smibert P, Papalexi E, Satija R. Integrating single-cell transcriptomic data across different conditions, technologies, and species. Nat Biotechnol. 2018;36(5):411–20. Epub 20180402. doi: 10.1038/nbt.4096. PubMed PMID: 29608179; PubMed Central PMCID: PMCPMC6700744.

52. McInnes LH, John; Saul, Nathaniel; and Grossberger, Lukas. UMAP: Uniform Manifold Approximation and Projection. The Journal of Open Source Software. 2018;3:861. doi: doi.org/10.48550/arXiv.1802.03426.

53. Farnsworth DR, Saunders LM, Miller AC. A single-cell transcriptome atlas for zebrafish development. Dev Biol. 2020;459(2):100–8. Epub 20191127. doi: 10.1016/j.ydbio.2019.11.008. PubMed PMID: 31782996; PubMed Central PMCID: PMCPMC7080588.

54. Saunders LM, Mishra AK, Aman AJ, Lewis VM, Toomey MB, Packer JS, et al. Thyroid hormone regulates distinct paths to maturation in pigment cell lineages. eLife. 2019;8. Epub 20190529. doi: 10.7554/eLife.45181. PubMed PMID: 31140974; PubMed Central PMCID: PMCPMC6588384.

55. Wagner DE, Weinreb C, Collins ZM, Briggs JA, Megason SG, Klein AM. Single-cell mapping of gene expression landscapes and lineage in the zebrafish embryo. Science. 2018;360(6392):981–7. Epub 20180426. doi: 10.1126/science.aar4362. PubMed PMID: 29700229; PubMed Central PMCID: PMCPMC6083445.

56. Brombin A, Simpson DJ, Travnickova J, Brunsdon H, Zeng Z, Lu Y, et al. Tfap2b specifies an embryonic melanocyte stem cell that retains adult multifate potential. Cell Rep. 2022;38(2):110234. doi: 10.1016/j.celrep.2021.110234. PubMed PMID: 35021087; PubMed Central PMCID: PMCPMC8764619.

57. Howe DG, Bradford YM, Conlin T, Eagle AE, Fashena D, Frazer K, et al. ZFIN, the Zebrafish Model Organism Database: increased support for mutants and transgenics. Nucleic Acids Res. 2013;41(Database issue):D854-60. Epub 20121015. doi: 10.1093/nar/gks938. PubMed PMID: 23074187; PubMed Central PMCID: PMCPMC3531097.

58. Mead TJ, Yutzey KE. Notch pathway regulation of neural crest cell development in vivo. Dev Dyn. 2012;241(2):376–89. Epub 20120103. doi: 10.1002/dvdy.23717. PubMed PMID: 22275227; PubMed Central PMCID: PMCPMC3266628.

59. McMenamin SK, Bain EJ, McCann AE, Patterson LB, Eom DS, Waller ZP, et al. Thyroid hormone-dependent adult pigment cell lineage and pattern in zebrafish. Science. 2014;345(6202):1358–61. Epub 20140828. doi: 10.1126/science.1256251. PubMed PMID: 25170046; PubMed Central PMCID: PMCPMC4211621.

60. Singh AP, Dinwiddie A, Mahalwar P, Schach U, Linker C, Irion U, et al. Pigment Cell Progenitors in Zebrafish Remain Multipotent through Metamorphosis. Dev Cell. 2016;38(3):316–30. Epub 20160721. doi: 10.1016/j.devcel.2016.06.020. PubMed PMID: 27453500.

61. Petratou K, Subkhankulova T, Lister JA, Rocco A, Schwetlick H, Kelsh RN. A systems biology approach uncovers the core gene regulatory network governing iridophore fate choice from the neural crest. PLoS genetics. 2018;14(10):e1007402. Epub 20181004. doi: 10.1371/journal.pgen.1007402. PubMed PMID: 30286071; PubMed Central PMCID: PMCPMC6191144.

62. Bagnara JT, Matsumoto J, Ferris W, Frost SK, Turner WA, Jr., Tchen TT, et al. Common origin of pigment cells. Science. 1979;203(4379):410–5. doi: 10.1126/science.760198. PubMed PMID: 760198.

63. Knight RD, Javidan Y, Nelson S, Zhang T, Schilling T. Skeletal and pigment cell defects in the lockjaw mutant reveal multiple roles for zebrafish tfap2a in neural crest development. Dev Dyn. 2004;229(1):87–98. Epub 2003/12/31. doi: 10.1002/dvdy.10494. PubMed PMID: 14699580.

64. Barrallo-Gimeno A, Holzschuh J, Driever W, Knapik EW. Neural crest survival and differentiation in zebrafish depends on mont blanc/tfap2a gene function. Development. 2004;131(7):1463–77. PubMed PMID: 14985255.

65. O’Brien EK, d’Alençon C, Bonde G, Li W, Schoenebeck J, Allende ML, et al. Transcription factor Ap-2alpha is necessary for development of embryonic melanophores, autonomic neurons and pharyngeal skeleton in zebrafish. Dev Biol. 2004;265(1):246–61. doi: 10.1016/j.ydbio.2003.09.029. PubMed PMID: 14697367.

66. Dooley CM, Wali N, Sealy IM, White RJ, Stemple DL, Collins JE, et al. The gene regulatory basis of genetic compensation during neural crest induction. PLoS genetics. 2019;15(6):e1008213. Epub 20190614. doi: 10.1371/journal.pgen.1008213. PubMed PMID: 31199790; PubMed Central PMCID: PMCPMC6594659.

67. Creyghton MP, Cheng AW, Welstead GG, Kooistra T, Carey BW, Steine EJ, et al. Histone H3K27ac separates active from poised enhancers and predicts developmental state. Proc Natl Acad Sci U S A. 2010;107(50):21931–6. doi: 10.1073/pnas.1016071107. PubMed PMID: 21106759; PubMed Central PMCID: PMCPMC3003124.

68. Pekowska A, Benoukraf T, Zacarias-Cabeza J, Belhocine M, Koch F, Holota H, et al. H3K4 tri-methylation provides an epigenetic signature of active enhancers. EMBO J. 2011;30(20):4198–210. Epub 2011/08/19. doi: 10.1038/emboj.2011.295. PubMed PMID: 21847099; PubMed Central PMCID: PMCPMC3199384.

69. Ringrose L, Paro R. Epigenetic regulation of cellular memory by the Polycomb and Trithorax group proteins. Annu Rev Genet. 2004;38:413–43. Epub 2004/12/01. doi: 10.1146/annurev.genet.38.072902.091907. PubMed PMID: 15568982.

70. Buenrostro JD, Giresi PG, Zaba LC, Chang HY, Greenleaf WJ. Transposition of native chromatin for fast and sensitive epigenomic profiling of open chromatin, DNA- binding proteins and nucleosome position. Nat Methods. 2013;10(12):1213–8. doi: 10.1038/nmeth.2688. PubMed PMID: 24097267; PubMed Central PMCID: PMCPMC3959825.

71. Lee BK, Uprety N, Jang YJ, Tucker SK, Rhee C, LeBlanc L, et al. Fosl1 overexpression directly activates trophoblast-specific gene expression programs in embryonic stem cells. Stem Cell Res. 2018;26:95–102. Epub 2017/12/23. doi: 10.1016/j.scr.2017.12.004. PubMed PMID: 29272857; PubMed Central PMCID: PMCPMC5899959.

72. Liu H, Tan BC, Tseng KH, Chuang CP, Yeh CW, Chen KD, et al. Nucleophosmin acts as a novel AP2alpha-binding transcriptional corepressor during cell differentiation. EMBO Rep. 2007;8(4):394–400. Epub 2007/02/24. doi: 10.1038/sj.embor.7400909. PubMed PMID: 17318229; PubMed Central PMCID: PMCPMC1852768.

73. Lin CY, Chao A, Wang TH, Lee LY, Yang LY, Tsai CL, et al. Nucleophosmin/B23 is a negative regulator of estrogen receptor alpha expression via AP2gamma in endometrial cancer cells. Oncotarget. 2016;7(37):60038–52. Epub 2016/08/17. doi: 10.18632/oncotarget.11048. PubMed PMID: 27527851; PubMed Central PMCID: PMCPMC5312367.

74. Wong PP, Miranda F, Chan KV, Berlato C, Hurst HC, Scibetta AG. Histone demethylase KDM5B collaborates with TFAP2C and Myc to repress the cell cycle inhibitor p21(cip) (CDKN1A). Mol Cell Biol. 2012;32(9):1633–44. Epub 2012/03/01. doi: 10.1128/MCB.06373-11. PubMed PMID: 22371483; PubMed Central PMCID: PMCPMC3347242.

75. Kim S, Yu NK, Kaang BK. CTCF as a multifunctional protein in genome regulation and gene expression. Exp Mol Med. 2015;47:e166. Epub 2015/06/06. doi: 10.1038/emm.2015.33. PubMed PMID: 26045254; PubMed Central PMCID: PMCPMC4491725.

76. Mavrothalassitis G, Ghysdael J. Proteins of the ETS family with transcriptional repressor activity. Oncogene. 2000;19(55):6524–32. Epub 2001/02/15. doi: 10.1038/sj.onc.1204045. PubMed PMID: 11175368.

77. Sekiya T, Zaret KS. Repression by Groucho/TLE/Grg proteins: genomic site recruitment generates compacted chromatin in vitro and impairs activator binding in vivo. Mol Cell. 2007;28(2):291–303. Epub 2007/10/30. doi: 10.1016/j.molcel.2007.10.002. PubMed PMID: 17964267; PubMed Central PMCID: PMCPMC2083644.

78. Watts JA, Zhang C, Klein-Szanto AJ, Kormish JD, Fu J, Zhang MQ, et al. Study of FoxA pioneer factor at silent genes reveals Rfx-repressed enhancer at Cdx2 and a potential indicator of esophageal adenocarcinoma development. PLoS genetics. 2011;7(9):e1002277. Epub 2011/09/22. doi: 10.1371/journal.pgen.1002277. PubMed PMID: 21935353; PubMed Central PMCID: PMCPMC3174211.

79. Vierbuchen T, Ling E, Cowley CJ, Couch CH, Wang X, Harmin DA, et al. AP-1 Transcription Factors and the BAF Complex Mediate Signal-Dependent Enhancer Selection. Mol Cell. 2017;68(6):1067–82.e12. Epub 2017/12/23. doi: 10.1016/j.molcel.2017.11.026. PubMed PMID: 29272704; PubMed Central PMCID: PMCPMC5744881.

80. Wilson BG, Wang X, Shen X, McKenna ES, Lemieux ME, Cho YJ, et al. Epigenetic antagonism between polycomb and SWI/SNF complexes during oncogenic transformation. Cancer Cell. 2010;18(4):316–28. Epub 2010/10/19. doi: 10.1016/j.ccr.2010.09.006. PubMed PMID: 20951942; PubMed Central PMCID: PMCPMC2957473.

81. Baxter LL, Watkins-Chow DE, Pavan WJ, Loftus SK. A curated gene list for expanding the horizons of pigmentation biology. Pigment Cell Melanoma Res. 2019;32(3):348–58. doi: 10.1111/pcmr.12743. PubMed PMID: 30339321.

82. Hoek KS, Eichhoff OM, Schlegel NC, Döbbeling U, Kobert N, Schaerer L, et al. In vivo switching of human melanoma cells between proliferative and invasive states. Cancer Res. 2008;68(3):650–6. Epub 2008/02/05. doi: 10.1158/0008-5472.Can-07-2491. PubMed PMID: 18245463.

83. Verfaillie A, Imrichova H, Atak ZK, Dewaele M, Rambow F, Hulselmans G, et al. Decoding the regulatory landscape of melanoma reveals TEADS as regulators of the invasive cell state. Nat Commun. 2015;6:6683. Epub 2015/04/14. doi: 10.1038/ncomms7683. PubMed PMID: 25865119; PubMed Central PMCID: PMCPMC4403341.

84. Hoek KS, Schlegel NC, Brafford P, Sucker A, Ugurel S, Kumar R, et al. Metastatic potential of melanomas defined by specific gene expression profiles with no BRAF signature. Pigment Cell Res. 2006;19(4):290–302. Epub 2006/07/11. doi: 10.1111/j.1600-0749.2006.00322.x. PubMed PMID: 16827748.

85. Tsoi J, Robert L, Paraiso K, Galvan C, Sheu KM, Lay J, et al. Multi-stage Differentiation Defines Melanoma Subtypes with Differential Vulnerability to Drug-Induced Iron-Dependent Oxidative Stress. Cancer Cell. 2018;33(5):890–904 e5. Epub 2018/04/17. doi: 10.1016/j.ccell.2018.03.017. PubMed PMID: 29657129; PubMed Central PMCID: PMCPMC5953834.

86. Jonsson G, Busch C, Knappskog S, Geisler J, Miletic H, Ringner M, et al. Gene expression profiling-based identification of molecular subtypes in stage IV melanomas with different clinical outcome. Clin Cancer Res. 2010;16(13):3356–67. Epub 2010/05/13. doi: 10.1158/1078-0432.CCR-09-2509. PubMed PMID: 20460471.

87. Tirosh I, Izar B, Prakadan SM, Wadsworth MH, 2nd, Treacy D, Trombetta JJ, et al. Dissecting the multicellular ecosystem of metastatic melanoma by single-cell RNA-seq. Science. 2016;352(6282):189-96. Epub 2016/04/29. doi: 10.1126/science.aad0501. PubMed PMID: 27124452; PubMed Central PMCID: PMCPMC4944528.

88. Rambow F, Rogiers A, Marin-Bejar O, Aibar S, Femel J, Dewaele M, et al. Toward Minimal Residual Disease-Directed Therapy in Melanoma. Cell. 2018;174(4):843–55 e19. Epub 2018/07/19. doi: 10.1016/j.cell.2018.06.025. PubMed PMID: 30017245.

89. Yu G, Wang LG, Han Y, He QY. clusterProfiler: an R package for comparing biological themes among gene clusters. Omics. 2012;16(5):284–7. Epub 2012/03/30. doi: 10.1089/omi.2011.0118. PubMed PMID: 22455463; PubMed Central PMCID: PMCPMC3339379.

90. Huang S, Jean D, Luca M, Tainsky MA, Bar-Eli M. Loss of AP-2 results in downregulation of c-KIT and enhancement of melanoma tumorigenicity and metastasis. EMBO J. 1998;17(15):4358–69. doi: 10.1093/emboj/17.15.4358. PubMed PMID: 9687504; PubMed Central PMCID: PMC1170769.

91. El-Brolosy MA, Kontarakis Z, Rossi A, Kuenne C, Günther S, Fukuda N, et al. Genetic compensation triggered by mutant mRNA degradation. Nature. 2019;568(7751):193–7. Epub 20190403. doi: 10.1038/s41586-019-1064-z. PubMed PMID: 30944477; PubMed Central PMCID: PMCPMC6707827.

92. Kumari P, Sturgeon M, Bonde G, Cornell RA. Generating Zebrafish RNA-Less Mutant Alleles by Deleting Gene Promoters with CRISPR/Cas9. Methods Mol Biol. 2022;2403:91–106. doi: 10.1007/978-1-0716-1847-9_8. PubMed PMID: 34913119.

93. White JR, Thompson DT, Koch KE, Kiriazov BS, Beck AC, van der Heide DM, et al. AP-2alpha-mediated activation of E2F and EZH2 drives melanoma metastasis. Cancer Res. 2021. Epub 2021/07/03. doi: 10.1158/0008-5472.CAN-21-0772. PubMed PMID: 34210752.

94. Mayran A, Khetchoumian K, Hariri F, Pastinen T, Gauthier Y, Balsalobre A, et al. Pioneer factor Pax7 deploys a stable enhancer repertoire for specification of cell fate. Nat Genet. 2018;50(2):259–69. Epub 2018/01/24. doi: 10.1038/s41588-017-0035-2. PubMed PMID: 29358650.

95. Braganca J, Eloranta JJ, Bamforth SD, Ibbitt JC, Hurst HC, Bhattacharya S. Physical and functional interactions among AP-2 transcription factors, p300/CREB-binding protein, and CITED2. The Journal of biological chemistry. 2003;278(18):16021–9. Epub 2003/02/15. doi: 10.1074/jbc.M208144200. PubMed PMID: 12586840.

96. Bejjani F, Evanno E, Zibara K, Piechaczyk M, Jariel-Encontre I. The AP-1 transcriptional complex: Local switch or remote command? Biochim Biophys Acta Rev Cancer. 2019;1872(1):11–23. Epub 2019/04/30. doi: 10.1016/j.bbcan.2019.04.003. PubMed PMID: 31034924.

97. Swinstead EE, Miranda TB, Paakinaho V, Baek S, Goldstein I, Hawkins M, et al. Steroid Receptors Reprogram FoxA1 Occupancy through Dynamic Chromatin Transitions. Cell. 2016;165(3):593–605. Epub 2016/04/12. doi: 10.1016/j.cell.2016.02.067. PubMed PMID: 27062924; PubMed Central PMCID: PMCPMC4842147.

98. Petruk S, Cai J, Sussman R, Sun G, Kovermann SK, Mariani SA, et al. Delayed Accumulation of H3K27me3 on Nascent DNA Is Essential for Recruitment of Transcription Factors at Early Stages of Stem Cell Differentiation. Mol Cell. 2017;66(2):247–57 e5. Epub 2017/04/16. doi: 10.1016/j.molcel.2017.03.006. PubMed PMID: 28410996; PubMed Central PMCID: PMCPMC5412717.

99. Curran K, Raible DW, Lister JA. Foxd3 controls melanophore specification in the zebrafish neural crest by regulation of Mitf. Dev Biol. 2009;332(2):408–17. Epub 20090613. doi: 10.1016/j.ydbio.2009.06.010. PubMed PMID: 19527705; PubMed Central PMCID: PMCPMC2716409.

100. Cao J, Spielmann M, Qiu X, Huang X, Ibrahim DM, Hill AJ, et al. The single-cell transcriptional landscape of mammalian organogenesis. Nature. 2019;566(7745):496–502. Epub 20190220. doi: 10.1038/s41586-019-0969-x. PubMed PMID: 30787437; PubMed Central PMCID: PMCPMC6434952.

101. Sander JD, Maeder ML, Reyon D, Voytas DF, Joung JK, Dobbs D. ZiFiT (Zinc Finger Targeter): an updated zinc finger engineering tool. Nucleic Acids Res. 2010;38(Web Server issue):W462–8. Epub 2010/05/04. doi: 10.1093/nar/gkq319. PubMed PMID: 20435679; PubMed Central PMCID: PMCPMC2896148.

102. Sander JD, Dahlborg EJ, Goodwin MJ, Cade L, Zhang F, Cifuentes D, et al. Selection-free zinc-finger-nuclease engineering by context-dependent assembly (CoDA). Nat Methods. 2011;8(1):67–9. Epub 2010/12/15. doi: nmeth.1542 [pii] 10.1038/nmeth.1542. PubMed PMID: 21151135; PubMed Central PMCID: PMC3018472.

103. Miller JC, Holmes MC, Wang J, Guschin DY, Lee YL, Rupniewski I, et al. An improved zinc-finger nuclease architecture for highly specific genome editing. Nat Biotechnol. 2007;25(7):778–85. Epub 2007/07/03. doi: 10.1038/nbt1319. PubMed PMID: 17603475.

104. Leroy B, Girard L, Hollestelle A, Minna JD, Gazdar AF, Soussi T. Analysis of TP53 mutation status in human cancer cell lines: a reassessment. Hum Mutat. 2014;35(6):756–65. Epub 2014/04/05. doi: 10.1002/humu.22556. PubMed PMID: 24700732; PubMed Central PMCID: PMCPMC4451114.

105. Liu H, Duncan K, Helverson A, Kumari P, Mumm C, Xiao Y, et al. Analysis of zebrafish periderm enhancers facilitates identification of a regulatory variant near human KRT8/18. eLife. 2020;9. Epub 2020/02/08. doi: 10.7554/eLife.51325. PubMed PMID: 32031521; PubMed Central PMCID: PMCPMC7039683.

106. Buenrostro JD, Wu B, Chang HY, Greenleaf WJ. ATAC-seq: A Method for Assaying Chromatin Accessibility Genome-Wide. Curr Protoc Mol Biol. 2015;109:21.9.1-.9.9. Epub 2015/01/07. doi: 10.1002/0471142727.mb2129s109. PubMed PMID: 25559105; PubMed Central PMCID: PMCPMC4374986.

107. Langmead B, Trapnell C, Pop M, Salzberg SL. Ultrafast and memory-efficient alignment of short DNA sequences to the human genome. Genome biology. 2009;10(3):R25. Epub 2009/03/06. doi: 10.1186/gb-2009-10-3-r25. PubMed PMID: 19261174; PubMed Central PMCID: PMCPMC2690996.

108. Langmead B, Salzberg SL. Fast gapped-read alignment with Bowtie 2. Nat Methods. 2012;9(4):357–9. Epub 2012/03/06. doi: 10.1038/nmeth.1923. PubMed PMID: 22388286; PubMed Central PMCID: PMCPMC3322381.

109. Ramírez F, Ryan DP, Grüning B, Bhardwaj V, Kilpert F, Richter AS, et al. deepTools2: a next generation web server for deep-sequencing data analysis. Nucleic Acids Res. 2016;44(W1):W160–5. Epub 2016/04/16. doi: 10.1093/nar/gkw257. PubMed PMID: 27079975; PubMed Central PMCID: PMCPMC4987876.

110. Zhang Y, Liu T, Meyer CA, Eeckhoute J, Johnson DS, Bernstein BE, et al. Model-based analysis of ChIP-Seq (MACS). Genome biology. 2008;9(9):R137. Epub 2008/09/19. doi: 10.1186/gb-2008-9-9-r137. PubMed PMID: 18798982; PubMed Central PMCID: PMCPMC2592715.

111. Ross-Innes CS, Stark R, Teschendorff AE, Holmes KA, Ali HR, Dunning MJ, et al. Differential oestrogen receptor binding is associated with clinical outcome in breast cancer. Nature. 2012;481(7381):389–93. Epub 2012/01/06. doi: 10.1038/nature10730. PubMed PMID: 22217937; PubMed Central PMCID: PMCPMC3272464.

112. McLean CY, Bristor D, Hiller M, Clarke SL, Schaar BT, Lowe CB, et al. GREAT improves functional interpretation of cis-regulatory regions. Nat Biotechnol. 2010;28(5):495–501. Epub 2010/05/04. doi:nbt.1630 [pii] 10.1038/nbt.1630. PubMed PMID: 20436461.

113. Skene PJ, Henikoff S. An efficient targeted nuclease strategy for high-resolution mapping of DNA binding sites. eLife. 2017;6. Epub 2017/01/13. doi: 10.7554/eLife.21856. PubMed PMID: 28079019; PubMed Central PMCID: PMCPMC5310842.

114. Meers MP, Bryson TD, Henikoff JG, Henikoff S. Improved CUT&RUN chromatin profiling tools. eLife. 2019;8. Epub 2019/06/25. doi: 10.7554/eLife.46314. PubMed PMID: 31232687; PubMed Central PMCID: PMCPMC6598765.

115. Dobin A, Davis CA, Schlesinger F, Drenkow J, Zaleski C, Jha S, et al. STAR: ultrafast universal RNA-seq aligner. Bioinformatics. 2013;29(1):15–21. Epub 2012/10/30. doi: 10.1093/bioinformatics/bts635. PubMed PMID: 23104886; PubMed Central PMCID: PMCPMC3530905.

116. Love MI, Huber W, Anders S. Moderated estimation of fold change and dispersion for RNA-seq data with DESeq2. Genome biology. 2014;15(12):550. Epub 2014/12/18. doi: 10.1186/s13059-014-0550-8. PubMed PMID: 25516281; PubMed Central PMCID: PMCPMC4302049.

117. Mi H, Ebert D, Muruganujan A, Mills C, Albou LP, Mushayamaha T, et al. PANTHER version 16: a revised family classification, tree-based classification tool, enhancer regions and extensive API. Nucleic Acids Res. 2021;49(D1):D394–d403. Epub 2020/12/09. doi: 10.1093/nar/gkaa1106. PubMed PMID: 33290554; PubMed Central PMCID: PMCPMC7778891.

118. Baxter LL, Loftus SK, Pavan WJ. Networks and pathways in pigmentation, health, and disease. Wiley interdisciplinary reviews Systems biology and medicine. 2009;1(3):359–71. Epub 2010/02/18. doi: 10.1002/wsbm.20. PubMed PMID: 20161540; PubMed Central PMCID: PMC2804986.

