## Supplemental Figures for "TFAP2 paralogs facilitate chromatin access for MITF at pigmentation genes but inhibit expression of cell-cell adhesion genes independently of MITF"

Fig. S1

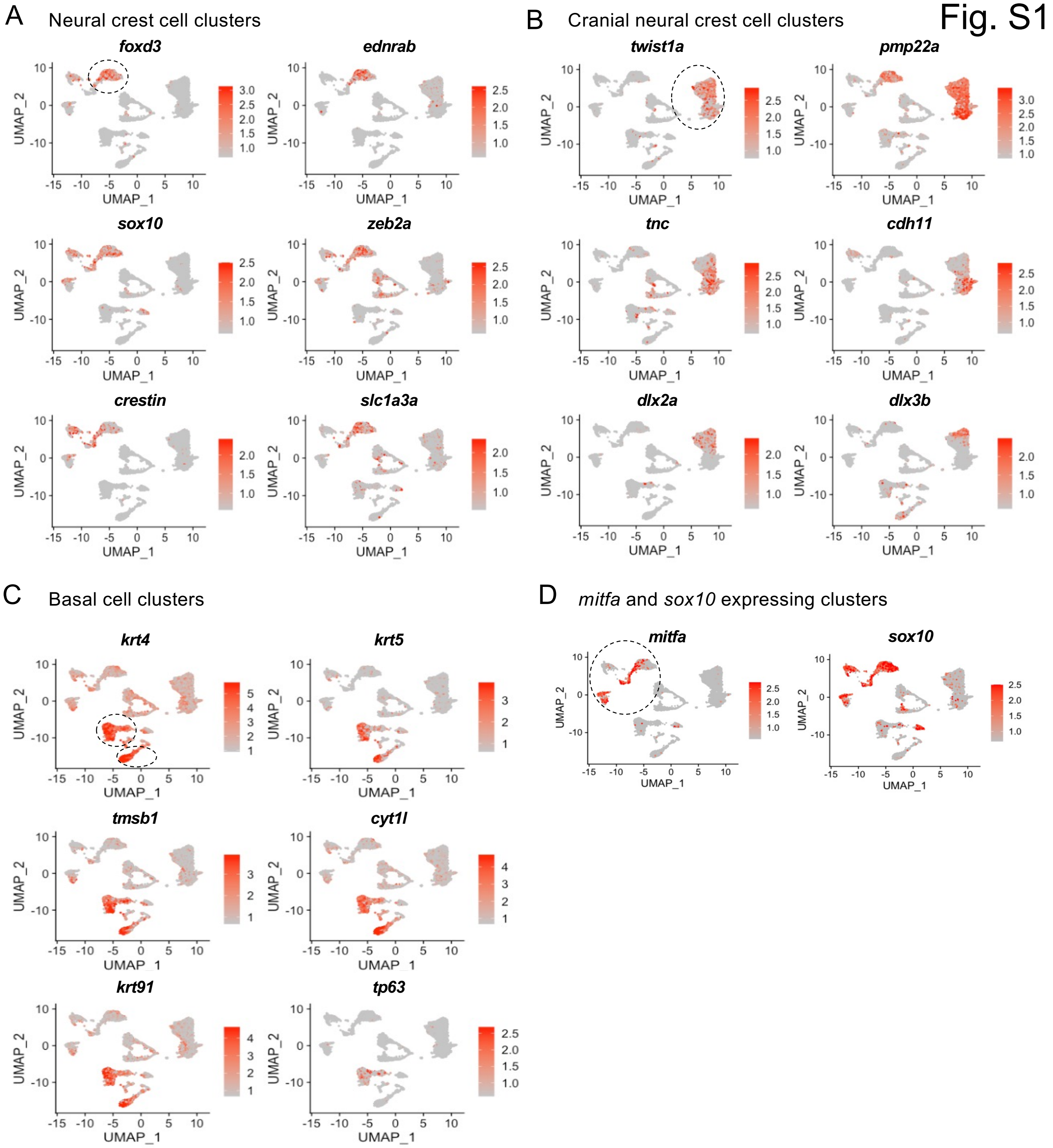

A
Neural crest cell clusters
foxd3
ednrab
sox10
zeb2a
crestin
slc1a3a
B
Cranial neural crest cell clusters
twist1a
pmp22a
tnc
cdh11
dlx2a
dlx3b
C
Basal cell clusters
krt4
krt5
tmsb1
cyt1l
krt91
tp63
D
mitfa and sox10 expressing clusters
mitfa
sox10
UMAP_1
UMAP_2

Fig. S2

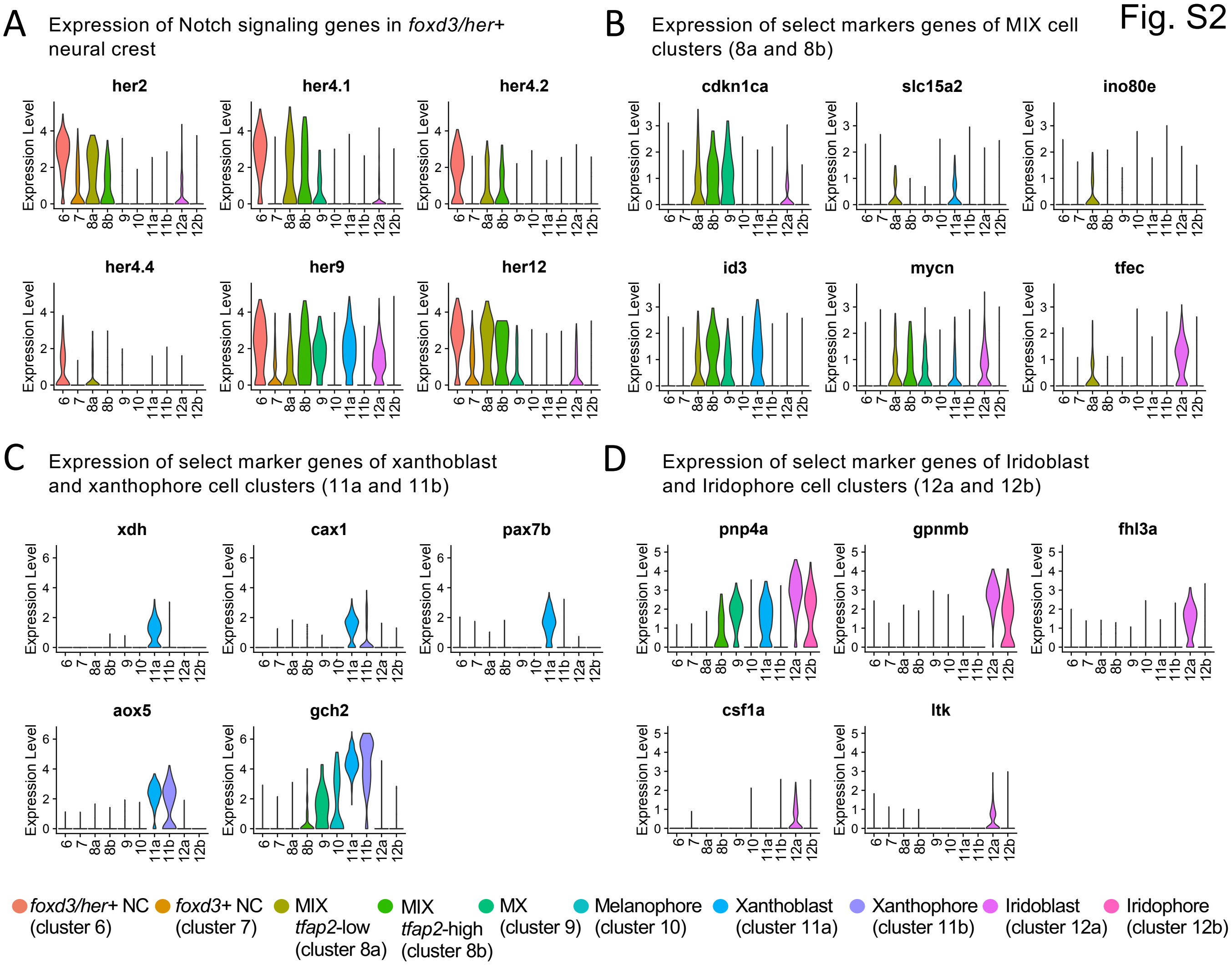
A
Expression of Notch signaling genes in *foxd3/her*+ neural crest
her2
her4.1
her4.2
her4.4
her9
her12
B
Expression of select markers genes of MIX cell clusters (8a and 8b)
cdkn1ca
slc15a2
ino80e
id3
mycn
tfec
C
Expression of select marker genes of xanthoblast and xanthophore cell clusters (11a and 11b)
xdh
cax1
pax7b
aox5
gch2
D
Expression of select marker genes of Iridoblast and Iridophore cell clusters (12a and 12b)
pnp4a
gpnmb
fhl3a
csf1a
ltk
Expression Level
6 7 8a 8b 9 10 11a 11b 12a 12b
*foxd3/her*+ NC (cluster 6)
*foxd3*+ NC (cluster 7)
MIX *tfap2*-low (cluster 8a)
MIX *tfap2*-high (cluster 8b)
MX (cluster 9)
Melanophore (cluster 10)
Xanthoblast (cluster 11a)
Xanthophore (cluster 11b)
Iridoblast (cluster 12a)
Iridophore (cluster 12b)

Fig. S3

A Expression of tfap2 paralog in pigment cell clusters (8a, 8b, 9 and 10)

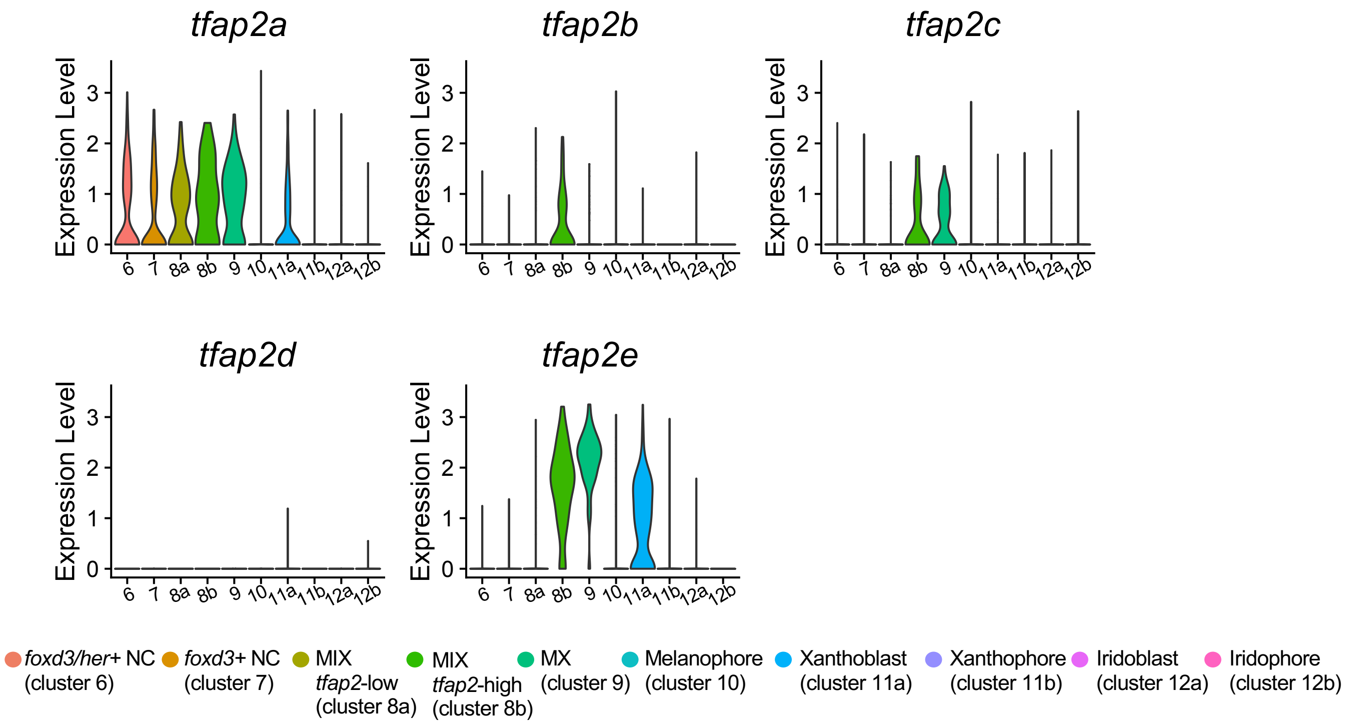

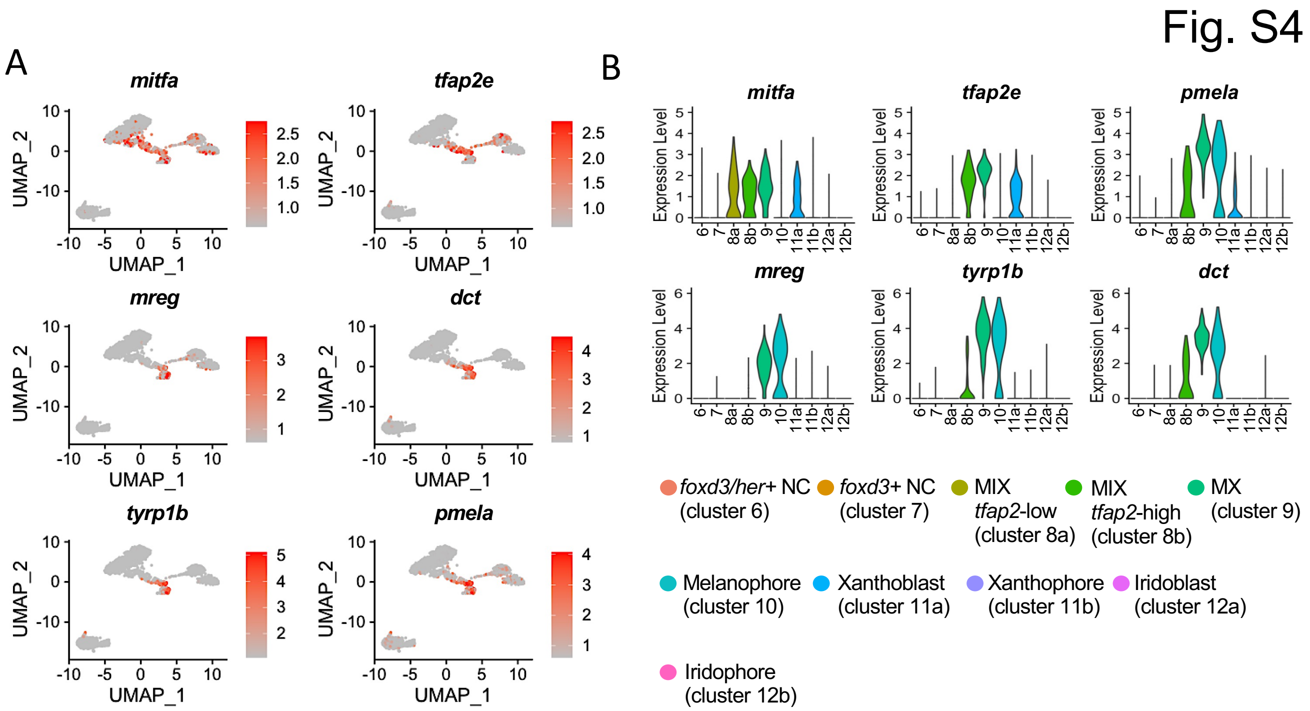

Fig. S4
A
mitfa
tfap2e
mreg
dct
tyrp1b
pmela
UMAP_1
UMAP_2
B
mitfa
tfap2e
pmela
mreg
tyrp1b
dct
Expression Level
6 7 8a 8b 9 10 11a 11b 12a 12b
foxd3/her+ NC (cluster 6)
foxd3+ NC (cluster 7)
MIX tfap2-low (cluster 8a)
MIX tfap2-high (cluster 8b)
MX (cluster 9)
Melanophore (cluster 10)
Xanthoblast (cluster 11a)
Xanthophore (cluster 11b)
Iridoblast (cluster 12a)
Iridophore (cluster 12b)

Fig. S5

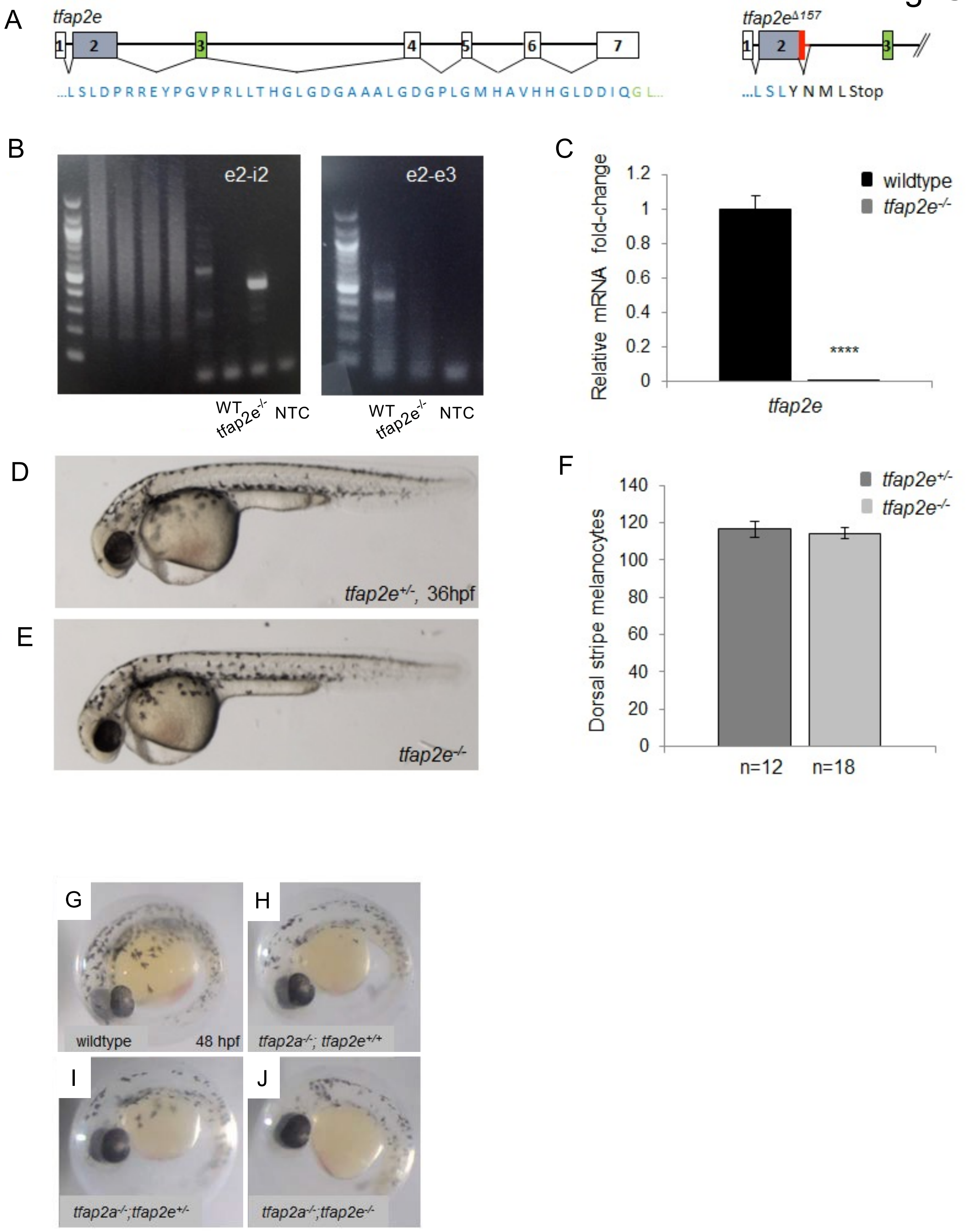
A
tfap2e
...LSLDPRREYPGVPRLLTHGLGDGAAALGDGPLGMHAVHHGLDDIQGL...
tfap2eΔ157
...LSLYNMLStop
B
e2-i2
e2-e3
WT
tfap2e-/-
NTC
C
Relative mRNA fold-change
wildtype
tfap2e-/-
****
tfap2e
D
tfap2e+/-, 36hpf
E
tfap2e-/-
F
Dorsal stripe melanocytes
tfap2e+/-
tfap2e-/-
n=12
n=18
G
wildtype
48 hpf
H
tfap2a-/-; tfap2e+/+
I
tfap2a-/-;tfap2e+/-
J
tfap2a-/-;tfap2e-/-

Fig. S6

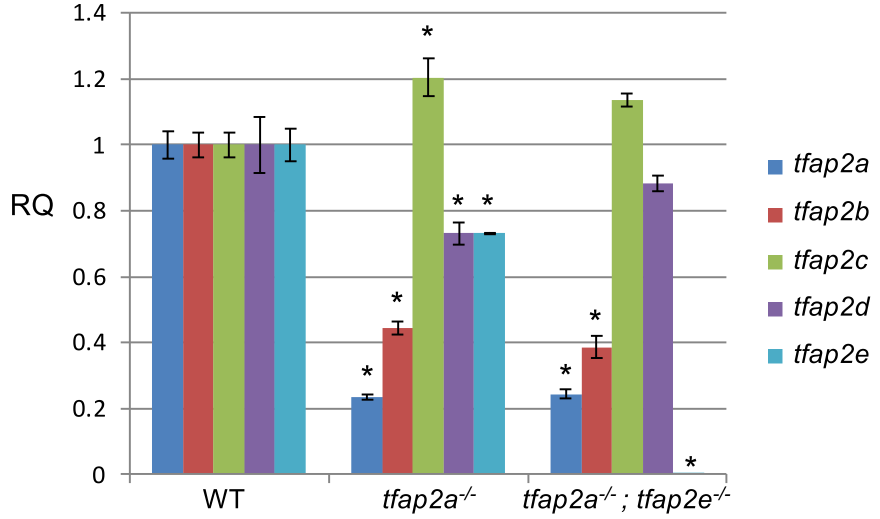

RQ
1.4
1.2
1
0.8
0.6
0.4
0.2
0
WT
tfap2a-/-
tfap2a-/- ; tfap2e-/-
*
tfap2a
tfap2b
tfap2c
tfap2d
tfap2e

Fig. S7

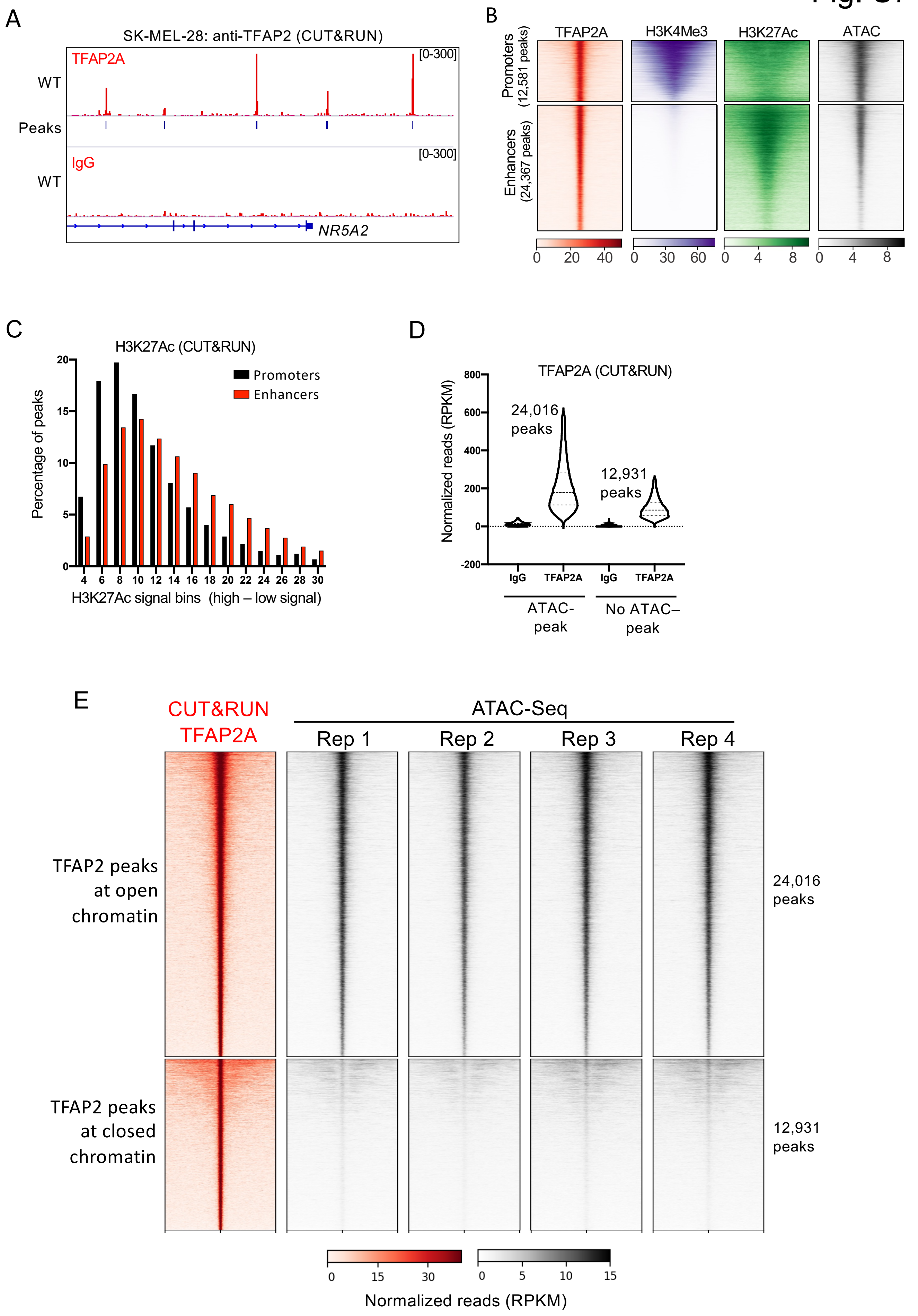

A
SK-MEL-28: anti-TFAP2 (CUT&RUN)
TFAP2A
WT
[0-300]
Peaks
IgG
WT
[0-300]
NR5A2
B
TFAP2A
H3K4Me3
H3K27Ac
ATAC
Promoters (12,581 peaks)
Enhancers (24,367 peaks)
0 20 40
0 30 60
0 4 8
0 4 8
C
H3K27Ac (CUT&RUN)
Promoters
Enhancers
Percentage of peaks
0 5 10 15 20
4 6 8 10 12 14 16 18 20 22 24 26 28 30
H3K27Ac signal bins (high – low signal)
D
TFAP2A (CUT&RUN)
Normalized reads (RPKM)
800 600 400 200 0 -200
24,016 peaks
12,931 peaks
IgG
TFAP2A
IgG
TFAP2A
ATAC-peak
No ATAC–peak
E
CUT&RUN TFAP2A
ATAC-Seq
Rep 1
Rep 2
Rep 3
Rep 4
TFAP2 peaks at open chromatin
TFAP2 peaks at closed chromatin
24,016 peaks
12,931 peaks
0 15 30
0 5 10 15
Normalized reads (RPKM)

### Fig. S8

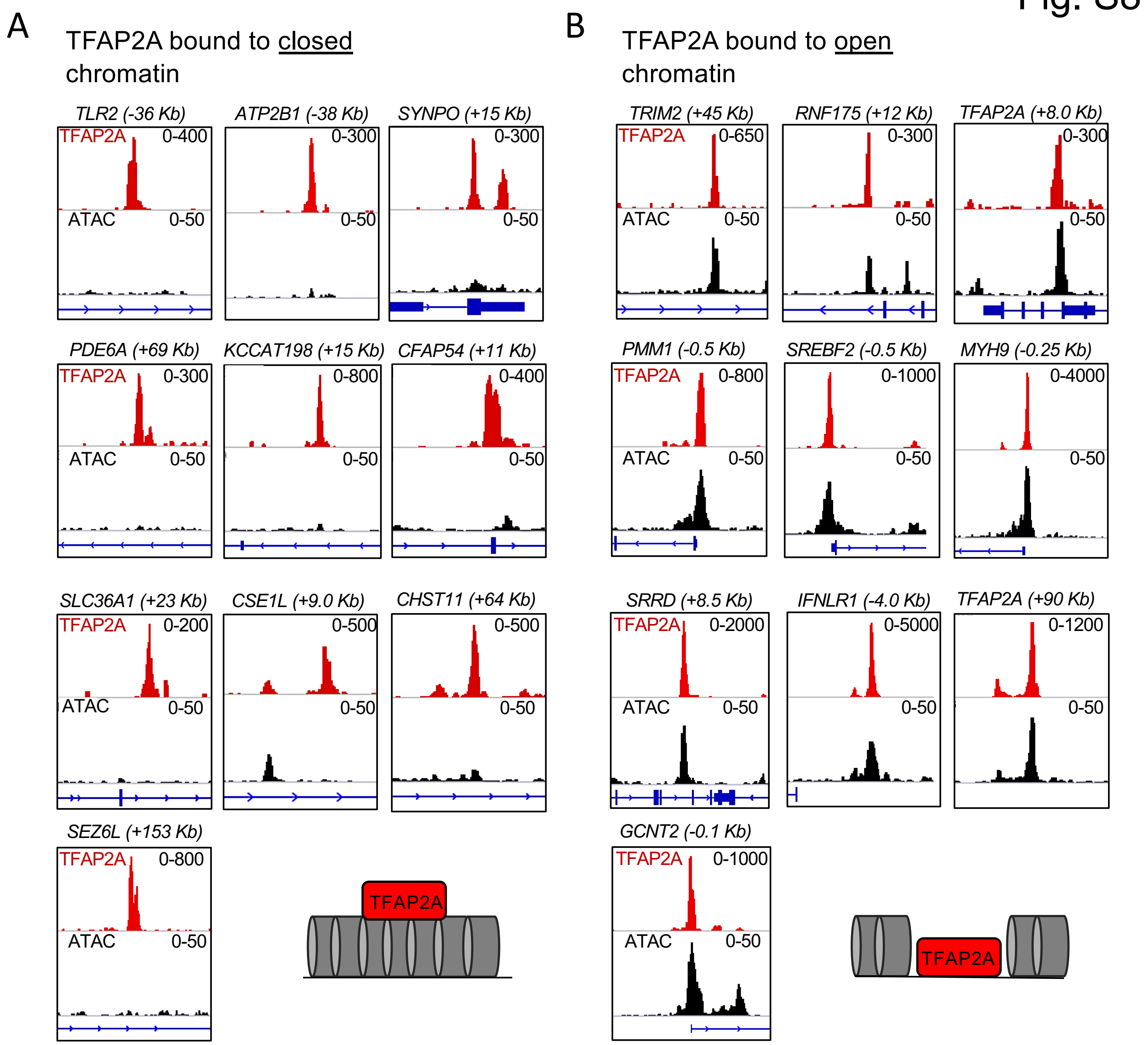


C

TFAP2A peaks not at ATAC peaks

| Motif | TF | Rank | Target | P-value |
|---|---|---|---|---|
| 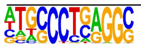 | TFAP2 | 1 | 59.8% | 1e-4375 |

D

TFAP2A peaks at ATAC peaks

| Motif | TF | Rank | Target | P-value |
|---|---|---|---|---|
| 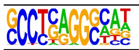 | TFAP2 | 1 | 61% | 1e-1785 |

Fig. S9

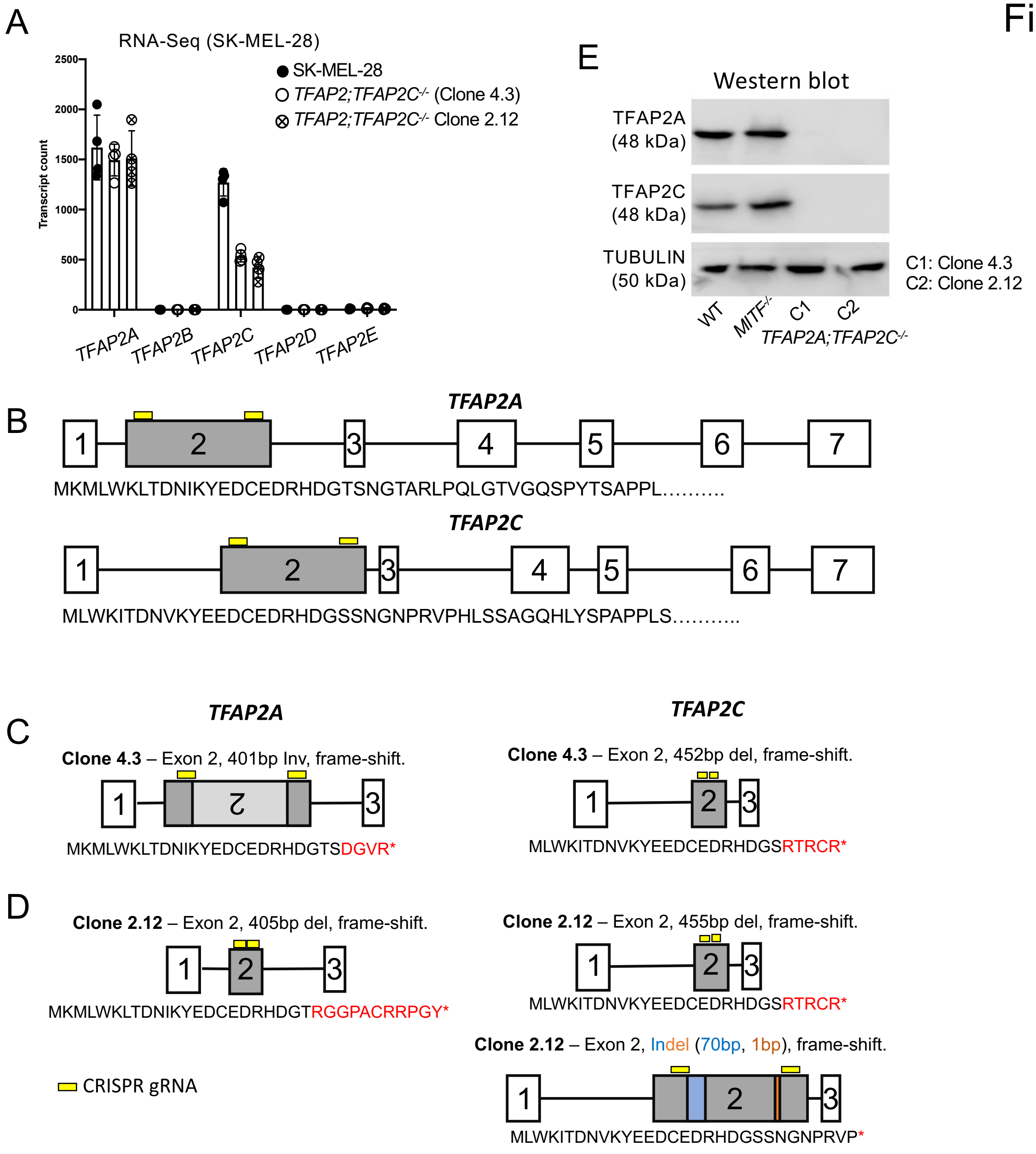

A
RNA-Seq (SK-MEL-28)
SK-MEL-28
TFAP2;TFAP2C-/- (Clone 4.3)
TFAP2;TFAP2C-/- Clone 2.12
Transcript count
2500
2000
1500
1000
500
0
TFAP2A
TFAP2B
TFAP2C
TFAP2D
TFAP2E
E
Western blot
TFAP2A (48 kDa)
TFAP2C (48 kDa)
TUBULIN (50 kDa)
C1: Clone 4.3
C2: Clone 2.12
WT
MITF-/-
C1
C2
TFAP2A;TFAP2C-/-
B
TFAP2A
MKMLWKLTDNIKYEDCEDRHDGTSNGTARLPQLGTVGQSPYTSAPPL..........
TFAP2C
MLWKITDNVKYEEDCEDRHDGSSNGNPRVPHLSSAGQHLYSPAPPLS...........
C
TFAP2A
Clone 4.3 – Exon 2, 401bp Inv, frame-shift.
MKMLWKLTDNIKYEDCEDRHDGTSDGVR*
TFAP2C
Clone 4.3 – Exon 2, 452bp del, frame-shift.
MLWKITDNVKYEEDCEDRHDGSRTRCR*
D
Clone 2.12 – Exon 2, 405bp del, frame-shift.
MKMLWKLTDNIKYEDCEDRHDGTRGGPACRRPGY*
Clone 2.12 – Exon 2, 455bp del, frame-shift.
MLWKITDNVKYEEDCEDRHDGSRTRCR*
Clone 2.12 – Exon 2, Indel (70bp, 1bp), frame-shift.
MLWKITDNVKYEEDCEDRHDGSSNGNPRVP*
CRISPR gRNA

Fig. S10

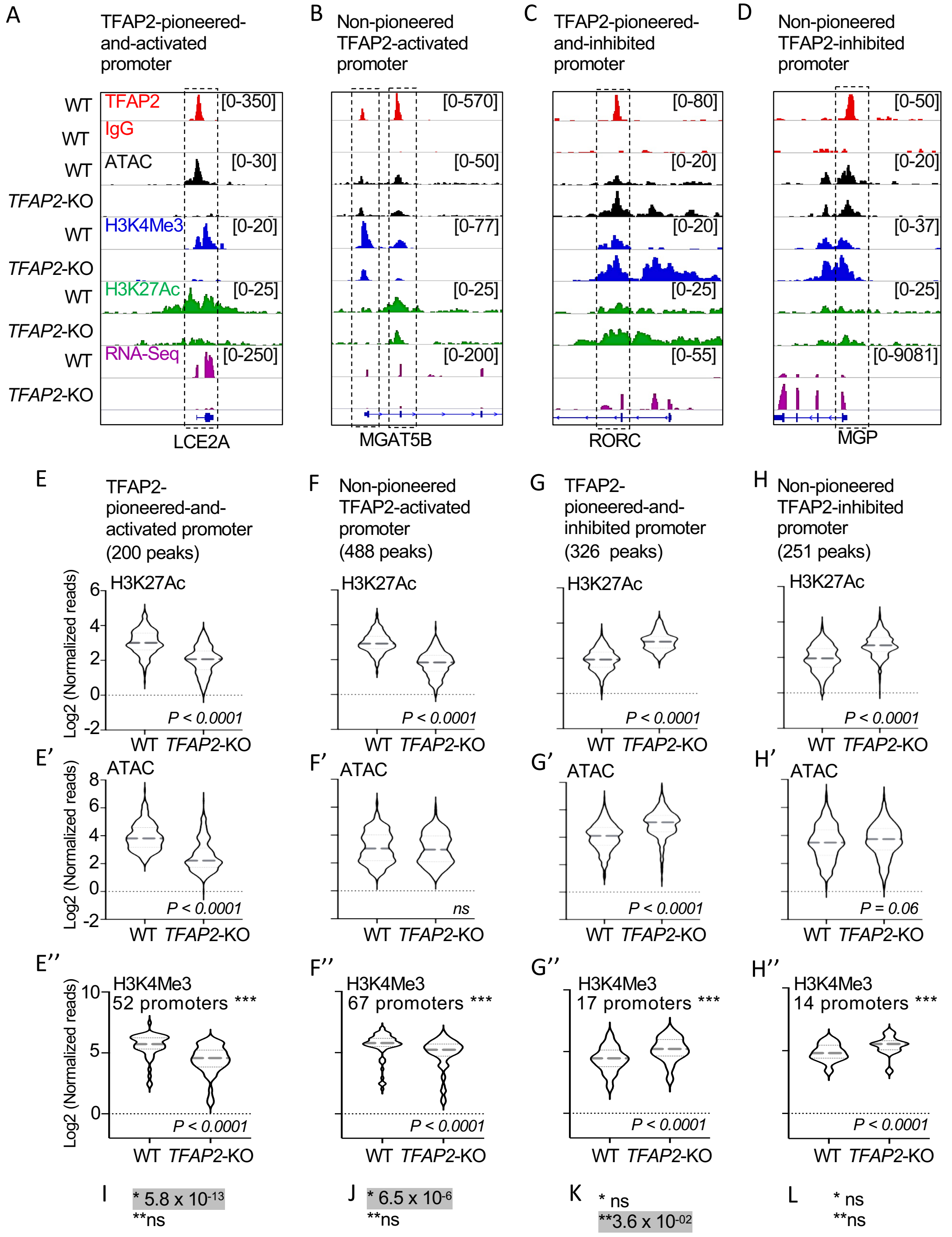


* p-val. Promoter (H3K27Ac/ATAC) with TFAP2-activated genes
**p-val. Promoter (H3K27Ac/ATAC) association with TFAP2-inhibited genes
*** Number of promoters with H3K27Ac, H3K4Me3 and ATAC activated or inhibited

Fig. S11

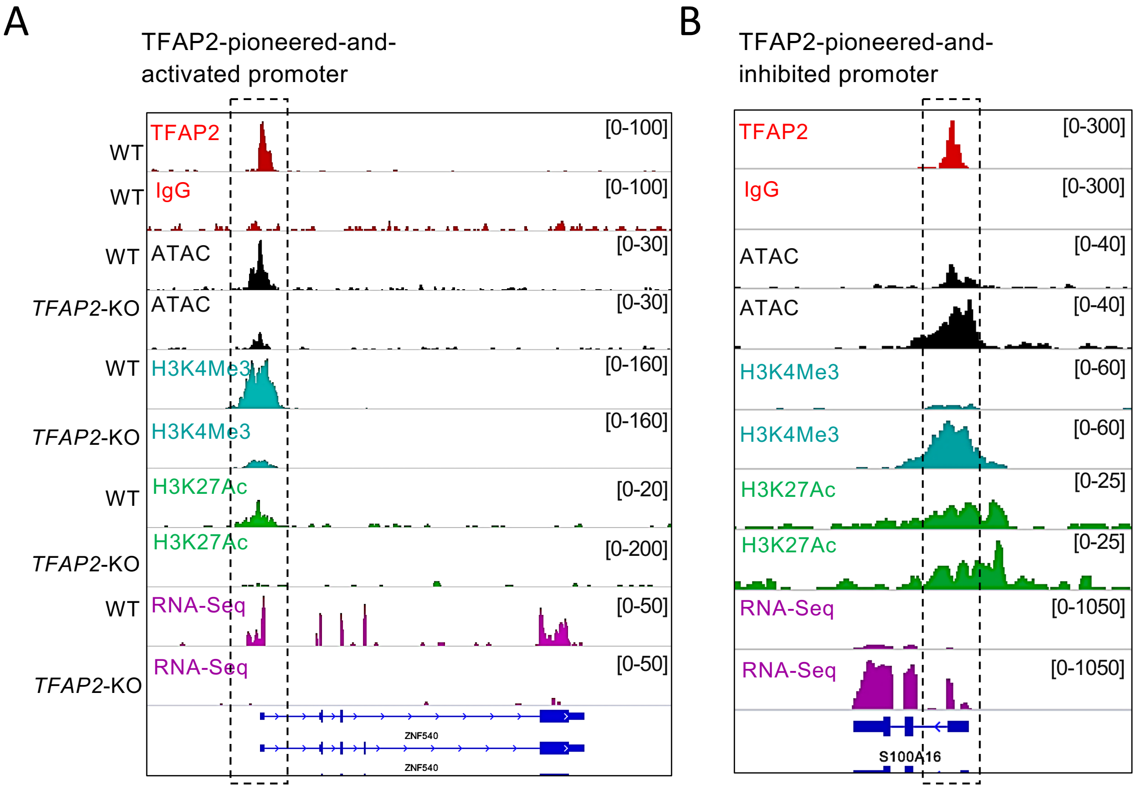
A
TFAP2-pioneered-and-
activated promoter
WT TFAP2 [0-100]
WT IgG [0-100]
WT ATAC [0-30]
*TFAP2*-KO ATAC [0-30]
WT H3K4Me3 [0-160]
*TFAP2*-KO H3K4Me3 [0-160]
WT H3K27Ac [0-20]
*TFAP2*-KO H3K27Ac [0-200]
WT RNA-Seq [0-50]
*TFAP2*-KO RNA-Seq [0-50]
ZNF540
ZNF540
B
TFAP2-pioneered-and-
inhibited promoter
TFAP2 [0-300]
IgG [0-300]
ATAC [0-40]
ATAC [0-40]
H3K4Me3 [0-60]
H3K4Me3 [0-60]
H3K27Ac [0-25]
H3K27Ac [0-25]
RNA-Seq [0-1050]
RNA-Seq [0-1050]
S100A16

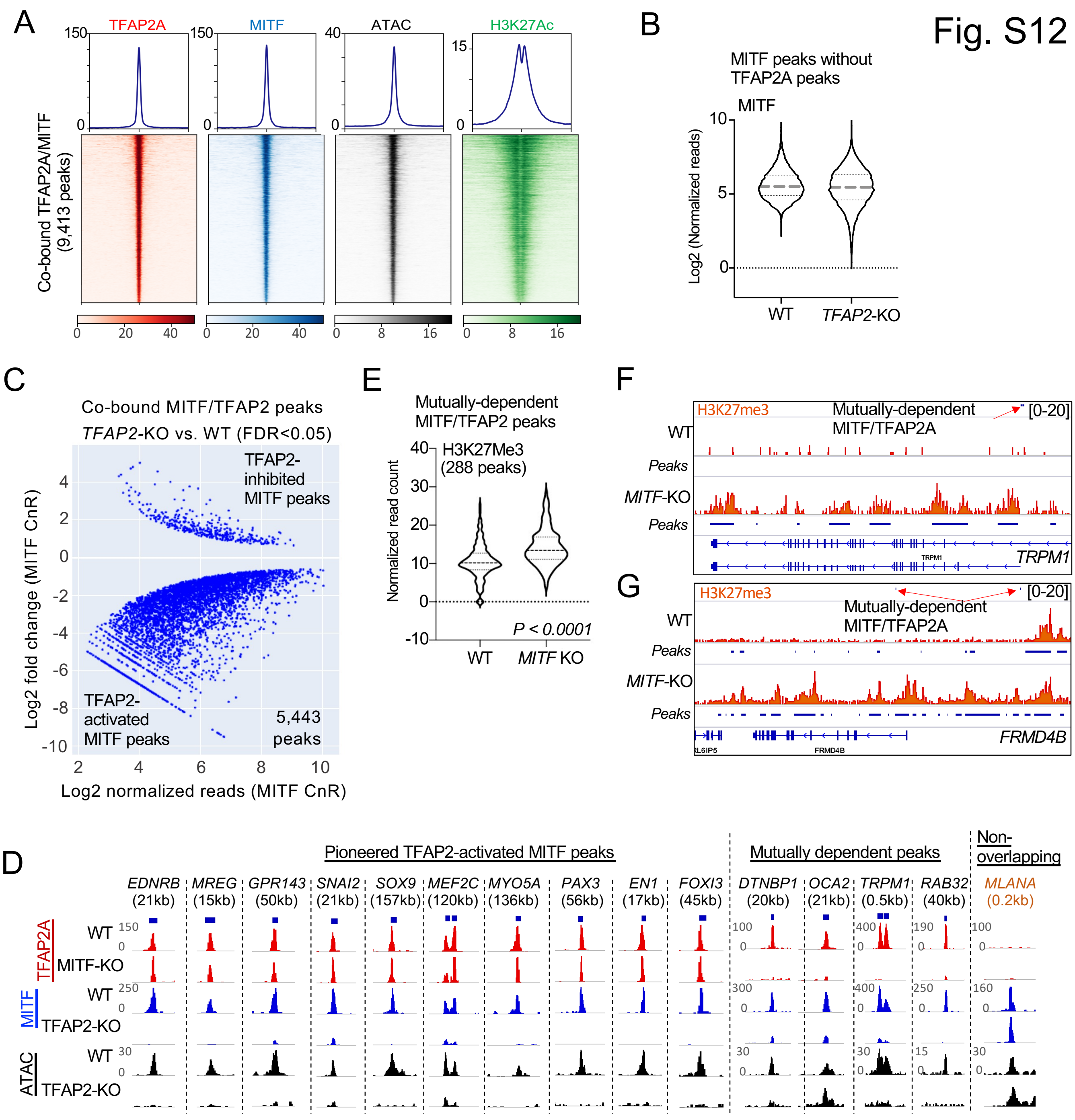
Fig. S12
A
TFAP2A
MITF
ATAC
H3K27Ac
Co-bound TFAP2A/MITF (9,413 peaks)
B
MITF peaks without TFAP2A peaks
MITF
Log2 (Normalized reads)
WT
TFAP2-KO
C
Co-bound MITF/TFAP2 peaks
TFAP2-KO vs. WT (FDR<0.05)
TFAP2-inhibited MITF peaks
TFAP2-activated MITF peaks
5,443 peaks
Log2 fold change (MITF CnR)
Log2 normalized reads (MITF CnR)
E
Mutually-dependent MITF/TFAP2 peaks
H3K27Me3 (288 peaks)
Normalized read count
P < 0.0001
WT
MITF KO
F
H3K27me3
Mutually-dependent MITF/TFAP2A
[0-20]
WT
Peaks
MITF-KO
TRPM1
G
H3K27me3
Mutually-dependent MITF/TFAP2A
[0-20]
WT
Peaks
MITF-KO
FRMD4B
D
Pioneered TFAP2-activated MITF peaks
Mutually dependent peaks
Non-overlapping
EDNRB (21kb)
MREG (15kb)
GPR143 (50kb)
SNAI2 (21kb)
SOX9 (157kb)
MEF2C (120kb)
MYO5A (136kb)
PAX3 (56kb)
EN1 (17kb)
FOXI3 (45kb)
DTNBP1 (20kb)
OCA2 (21kb)
TRPM1 (0.5kb)
RAB32 (40kb)
MLANA (0.2kb)
TFAP2A
WT
MITF-KO
MITF
WT
TFAP2-KO
ATAC
WT
TFAP2-KO

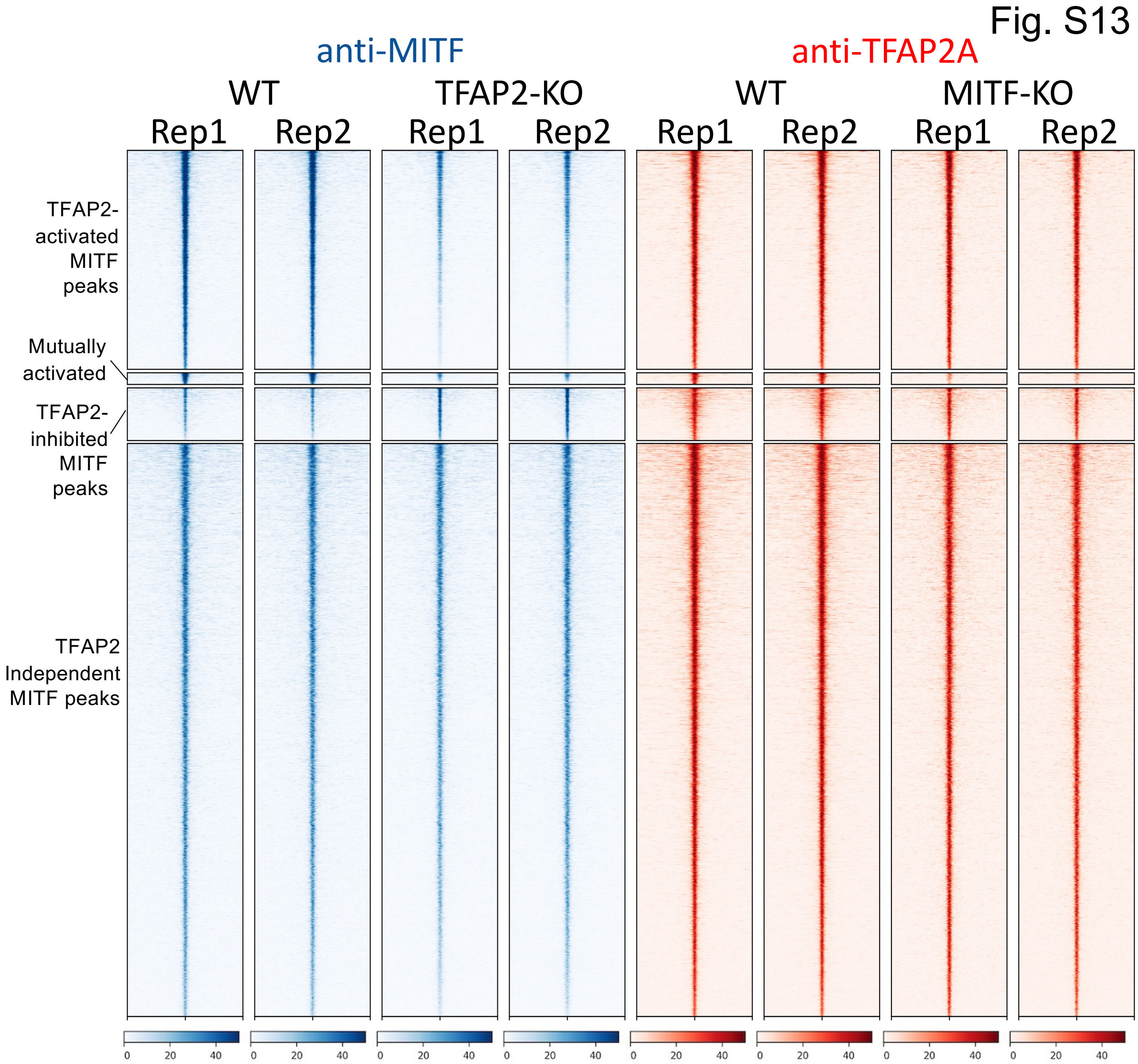
Fig. S13
anti-MITF
anti-TFAP2A
WT
TFAP2-KO
WT
MITF-KO
Rep1
Rep2
Rep1
Rep2
Rep1
Rep2
Rep1
Rep2
TFAP2-activated MITF peaks
Mutually activated
TFAP2-inhibited MITF peaks
TFAP2 Independent MITF peaks
0
20
40

### Fig. S14

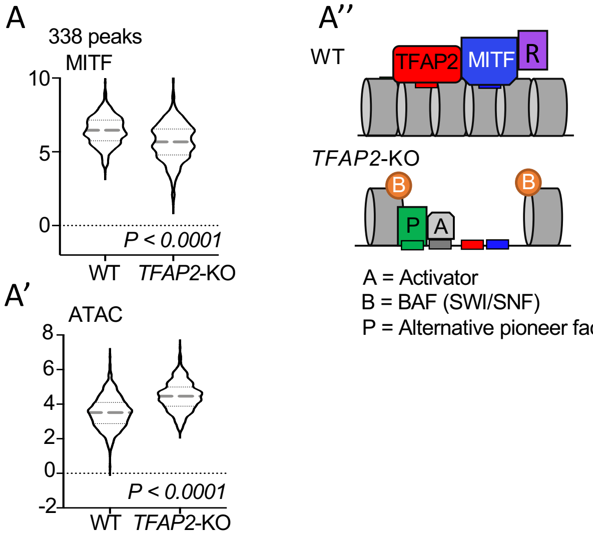


A = Activator
B = BAF (SWI/SNF)
P = Alternative pioneer factor

Association with gene expression:
MITF-inhibited, ns
TFAP2-inhibited, ns

### Fig. S15

A

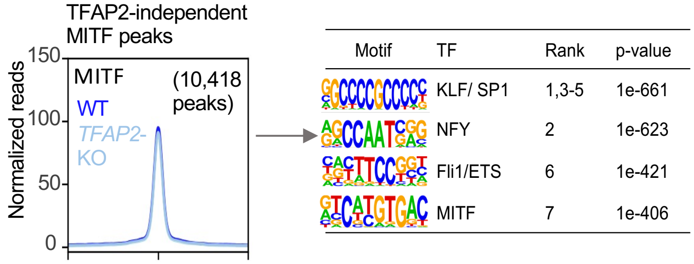
TFAP2-independent
MITF peaks
Normalized reads
150
100
50
0
MITF
WT
TFAP2-
KO
(10,418
peaks)
Motif
TF
Rank
p-value
KLF/ SP1
1,3-5
1e-661
NFY
2
1e-623
Fli1/ETS
6
1e-421
MITF
7
1e-406

B

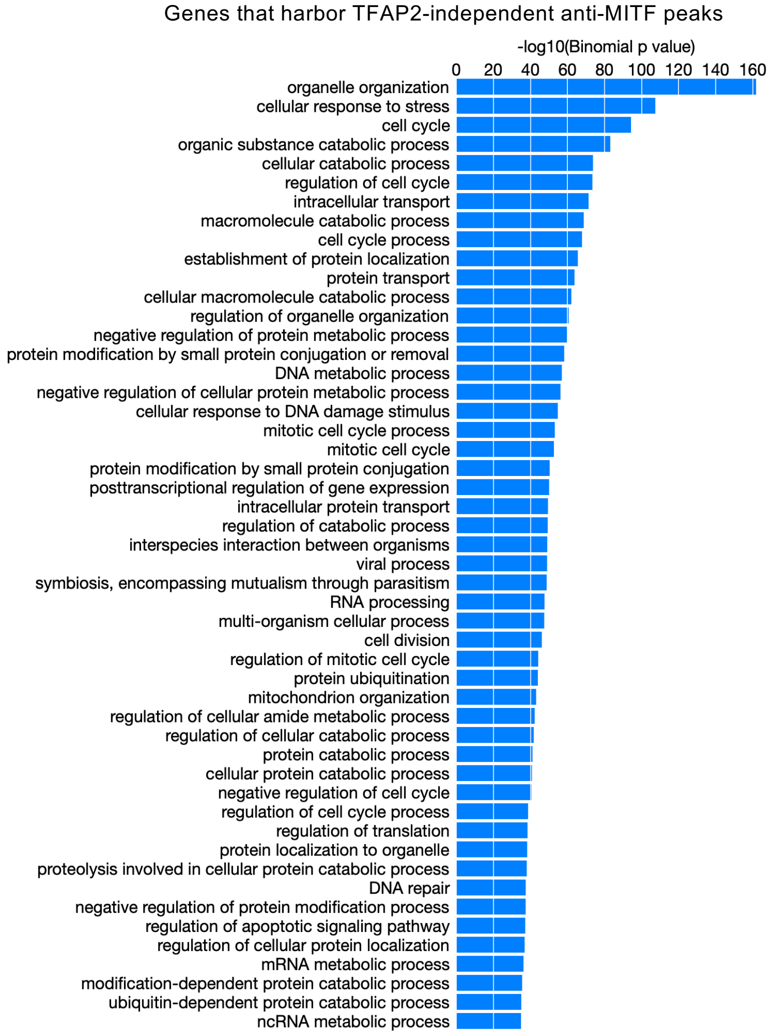
Genes that harbor TFAP2-independent anti-MITF peaks
-log10(Binomial p value)
0
20
40
60
80
100
120
140
160
organelle organization
cellular response to stress
cell cycle
organic substance catabolic process
cellular catabolic process
regulation of cell cycle
intracellular transport
macromolecule catabolic process
cell cycle process
establishment of protein localization
protein transport
cellular macromolecule catabolic process
regulation of organelle organization
negative regulation of protein metabolic process
protein modification by small protein conjugation or removal
DNA metabolic process
negative regulation of cellular protein metabolic process
cellular response to DNA damage stimulus
mitotic cell cycle process
mitotic cell cycle
protein modification by small protein conjugation
posttranscriptional regulation of gene expression
intracellular protein transport
regulation of catabolic process
interspecies interaction between organisms
viral process
symbiosis, encompassing mutualism through parasitism
RNA processing
multi-organism cellular process
cell division
regulation of mitotic cell cycle
protein ubiquitination
mitochondrion organization
regulation of cellular amide metabolic process
regulation of cellular catabolic process
protein catabolic process
cellular protein catabolic process
negative regulation of cell cycle
regulation of cell cycle process
regulation of translation
protein localization to organelle
proteolysis involved in cellular protein catabolic process
DNA repair
negative regulation of protein modification process
regulation of apoptotic signaling pathway
regulation of cellular protein localization
mRNA metabolic process
modification-dependent protein catabolic process
ubiquitin-dependent protein catabolic process
ncRNA metabolic process

Fig. S16

TFAP2-activated enhancer at intron 2 of MITF

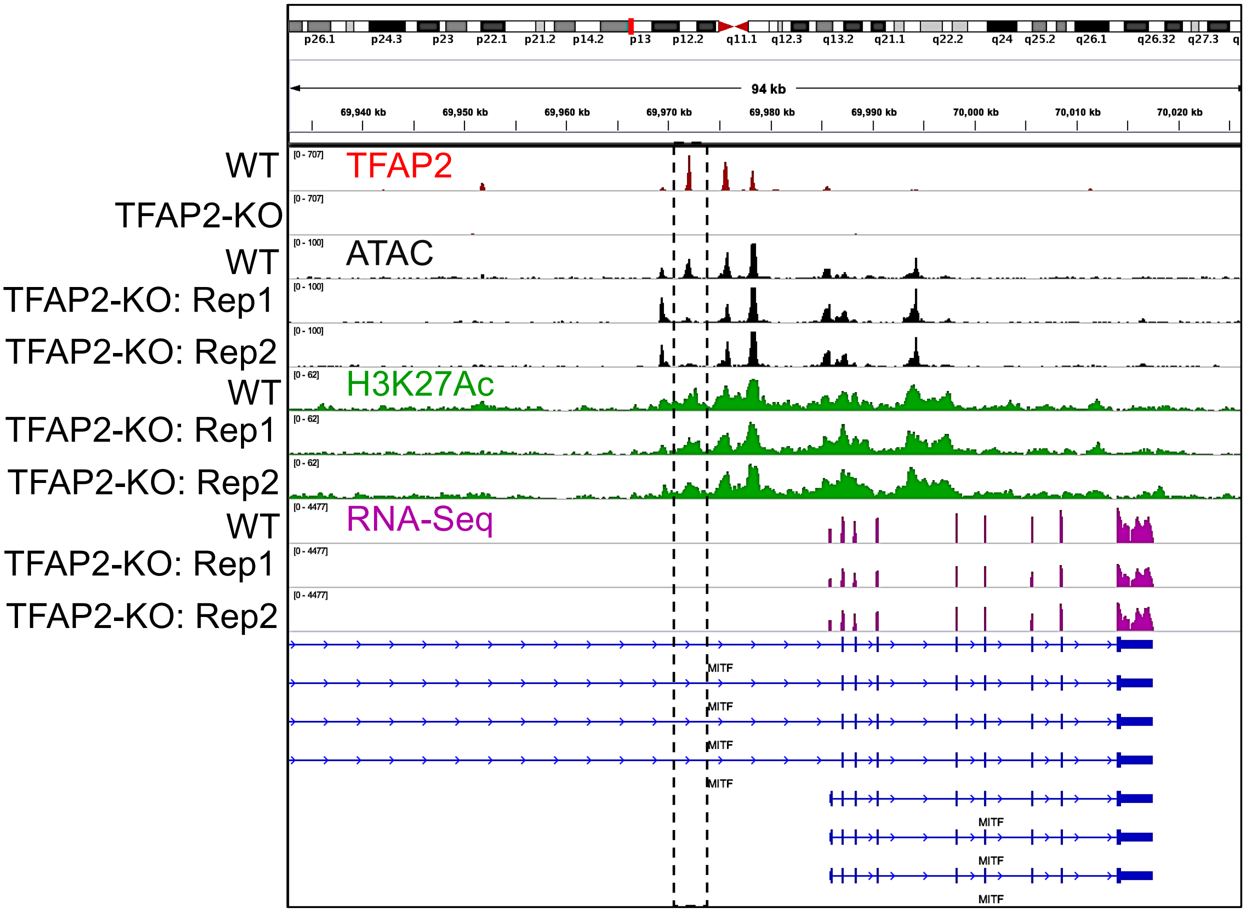

Fig. S17

A

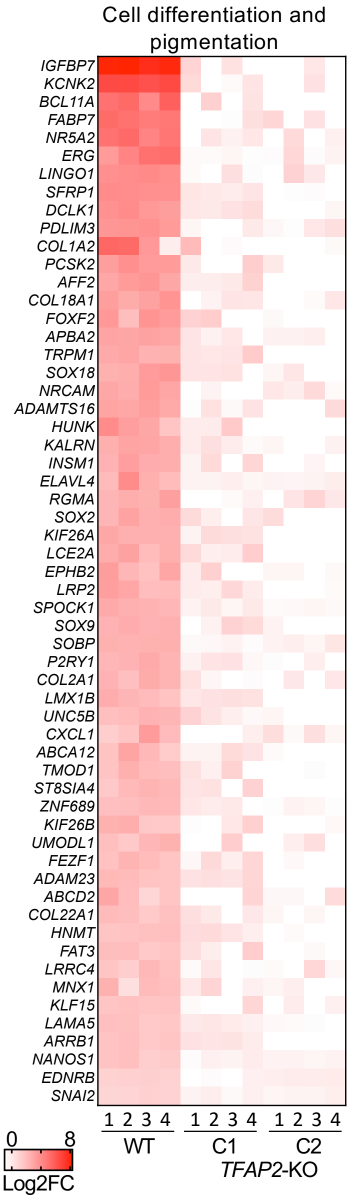


C1: Clone 4.3
C2: Clone 2.12

B

Directly TFAP2 activated genes

| GO biological process | Enrichment | FDR |
|---|---|---|
| Pigmentation | 4.5 | $8.8 \times 10^{-04}$ |
| Epithelial/ mesenchymal transition | 4.42 | $6.0 \times 10^{-03}$ |
| Cardiac septum development | 3.22 | $4.6 \times 10^{-02}$ |
| Kidney epithelium development | 3.07 | $3.0 \times 10^{-02}$ |
| Mesenchymal cell differentiation | 3.07 | $7.1 \times 10^{-03}$ |
| Cell morphogenesis | 2.36 | $1.3 \times 10^{-06}$ |
| Phosphorylation | 2.17 | $3.6 \times 10^{-06}$ |
| Neuron differentiation | 2.14 | $2.6 \times 10^{-07}$ |
| Cell differentiation | 1.62 | $1.0 \times 10^{-09}$ |
| Cell proliferation | 1.49 | $2.4 \times 10^{-02}$ |

C

Directly TFAP2 inhibited genes

| GO biological process | Enrichment | FDR |
|---|---|---|
| Angiogenesis | 6.93 | $4.9 \times 10^{-02}$ |
| Cell junction | 2.96 | $3.0 \times 10^{-08}$ |
| Leukocyte migration | 2.77 | $5.5 \times 10^{-03}$ |
| Morphogenesis of an epithelium | 2.77 | $4.0 \times 10^{-06}$ |
| Response to growth factor | 2.54 | $8.6 \times 10^{-06}$ |
| Cell-cell adhesion | 2.45 | $2.2 \times 10^{-04}$ |
| Cell migration | 2.41 | $1.5 \times 10^{-08}$ |
| Tube morphogenesis | 2.36 | $9.2 \times 10^{-06}$ |
| Cell motility | 2.32 | $4.6 \times 10^{-08}$ |
| Locomotion | 2.3 | $2.4 \times 10^{-08}$ |
| Programmed cell death | 1.9 | $7.6 \times 10^{-07}$ |
| Cell communication | 1.78 | $6.5 \times 10^{-14}$ |

Fig. S18

A

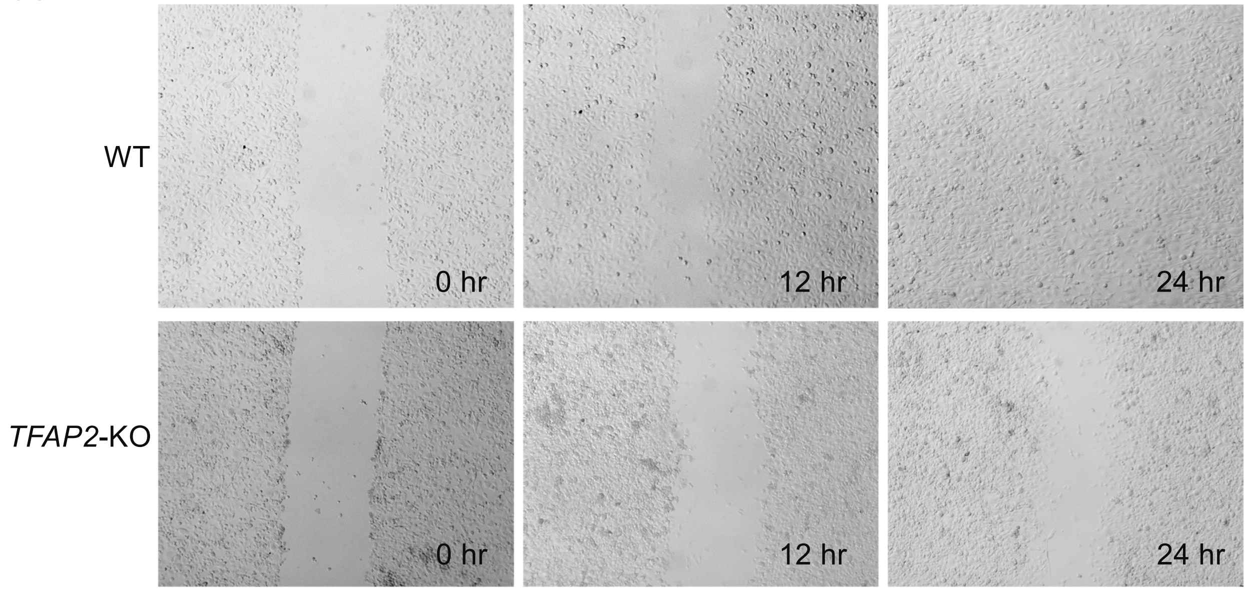
